# Comprehensive multi-omics profiling of a healthy human cohort

**DOI:** 10.1101/2024.11.07.622407

**Authors:** Casper de Visser, Anna Niehues, Lukáš Najdekr, Reka Toth, Petr V. Nazarov, Jana Vrbková, Jarmila Stanková, Sara Ekberg, Elisa Conde Moreno, Bishwa Ghimire, Bhagwan Yadav, Pirkko Matilla, Maija Puhka, Val F. Lanza, Jolein Gloerich, Purva Kulkarni, Hans Wessels, Udo Engelke, Emanuela Oldoni, Toni Andreu, Gary Saunders, Arnaud Muller, Michaela Bendová, Zuzana Rozankova, Petr Dzubak, Petr Pavlis, Bronislav Siska, Aikaterini Spantidaki, Daniël Beisly, Josef Srovnal, Viktorie Moravíková, Petr Šesták, Matěj Gazdarica, Andreas Scherer, M Laura Garcia Bermejo, Jessica Nordlund, Alain J. van Gool, Marián Hajdúch, Peter A.C. ’t Hoen

**Author notes:** ^†^Shared 2nd author.

## Abstract

Multi-omics approaches can offer powerful insights into personalized biomarker profiles relevant for disease diagnosis, prognosis, and therapeutics. However, separating meaningful biological variability from technical noise remains a major challenge. The EATRIS-Plus consortium analyzed blood samples from 127 healthy adults across six omics layers using twelve platforms, resulting in one of the most comprehensive multi-omics profiling datasets of healthy individuals available to date. We applied reproducible workflows to analyze and integrate these data, revealing several key findings. Sex significantly influenced all omics layers, emphasizing the importance of sex-balanced study designs. Age could be accurately predicted using epigenetic clocks, achieving high performance with our high-resolution enzymatic methylation sequencing data (*R*^2^ = 0.90), whereas candidate aging biomarkers were identified across all omics layers. The resulting dataset provides reference ranges in healthy individuals for abundance and variability of omics features, enabling robust power analyses, sample size estimations, and benchmarking of multi-omics integration methods. This resource can guide future biomarker discovery and personalized health research and was made FAIR-compliant and publicly available via the ClinData Portal (https://clindata.imtm.cz) and a Zenodo repository (https://doi.org/10.5281/zenodo.17514796).

## Introduction

Multi-omics studies have a large potential for uncovering biomarker profiles that are relevant for disease diagnosis, prognosis, or prediction of the efficacy of medical interventions. Compared to single-omics analysis, multi-omics interrogations of biological systems provide more comprehensive molecular data, deeper insights into biology, and a better understanding of mechanisms underlying health and disease. They also contribute to studying the impact of lifestyle on well-being at the molecular level, which is useful for identifying individuals or populations at risk and for the stratification of participants in clinical trials. However, these complex studies are subject to many different sources of technical and inter- and intra-individual variation. Capturing biological variability in healthy individuals and reducing unwanted technical variation can help to plan and interpret future translational disease and interventional studies in precision medicine^1–3^. Currently, there are many open questions regarding the design and analysis of multi-omics studies: What are the challenges in multi-center multi-omics studies? Which biological and technical confounders should be considered? How are the different omics layers related and functionally connected? Which omics layers capture complementary biological information, and which are largely repetitive and redundant?

To answer these questions, we designed the EATRIS-Plus project (https://www.eatris.eu/projects/eatris-plus), executing one of the most comprehensive multi-omics studies in a human healthy population cohort. In this reference study, we investigated six omics layers with twelve different omics data platforms to uncover sources of technical and biological variation. Mapping the variability in the resulting omics datasets facilitates sample-size calculations for future single and multi-omics studies, as demonstrated previously^4,5^. Assessing the variability of individual features in this reference cohort will help to select robust biomarkers for disease or lifestyle in future diagnostic or prognostic studies. Through testing a range of multi-omics data integration methods, we investigated the complementarity of the different omics datasets. This should facilitate the selection of the best combination of omics platforms in future multi-omics studies, where budgets or technical implementation in clinical diagnostics may not allow the profiling of multiple omics layers. Finally, we investigated the association of the molecular features with the participants’ phenotypes, such as sex, (biological) age and body mass index (BMI), and explored the biological nature of the observed variation through pathway analyses.

## Results

### Study Design

This study profiled blood samples obtained from a cohort of 127 healthy individuals. These individuals were selected from a larger cohort of blood donors who were clinically examined and whose samples were bio-banked at the Institute of Molecular and Translational Medicine of Palacký University and University Hospital Olomouc (see Methods). We selected a group of individuals with equal representation of the sexes and a representative adult age distribution between 21 and 61 years (Table S1-2). All samples were collected using standardized sampling procedures and underwent comprehensive analysis using 12 different omics platforms, including **genomics** (whole genome sequencing (WGS) and array comparative genomic hybridization (aCGH) on blood cells), **epigenomics** (DNA methylation using enzymatic methylation sequencing (EM-seq) on blood cells)^6,7^, **transcriptomics** (mRNA and miRNA sequencing on blood cells and qPCR-based miRNA profiling on heparin or EDTA plasma), **proteomics** (shotgun mass spectrometry (MS)-based proteomics and NGS-based proteomics (Illumina Protein Prep [IPP]) on heparin plasma), **metabolomics** (liquid chromatography (LC)-MS-based targeted metabolomics (amino acids, very long chain fatty acids (VLCFA) and acylcarnitines) on heparin plasma), and three **lipidomics** datasets generated using the same platform (LC-MS-based untargeted lipidomics on heparin plasma) (Figure 1, Table S3). This extensive setup allows us to compare different omics layers, and multiple datasets for the same omics type. Samples were distributed to EATRIS facilities across Europe with relevant expertise for profiling the different omics layers. Quality control and processing of the individual omics data were performed locally. A subset of samples in the sequencing-based methods did not meet quality control standards, *e.g.* hemolysis grade for qRT-PCR (see Methods). Moreover, two individuals showed discordance between genotypes in the different omics layers or discordance between genotype and blood group due to potential sample-swaps (see Methods) and were removed from all analyses. From the remaining 125 individuals, all omics datasets are available for 53 individuals, with 108 individuals having at least nine different omics datasets (Figure 1). All processed and quality-checked data are available via doi.org/10.5281/zenodo.17514796 and https://fdp.radboudumc.nl and provides a rich resource for benchmarking multi-omics data integration methods. Uncorrected raw data and GDPR-sensitive data are available upon request in machine-readable formats in ClinData Portal (https://clindata.imtm.cz).

**Fig 1.**
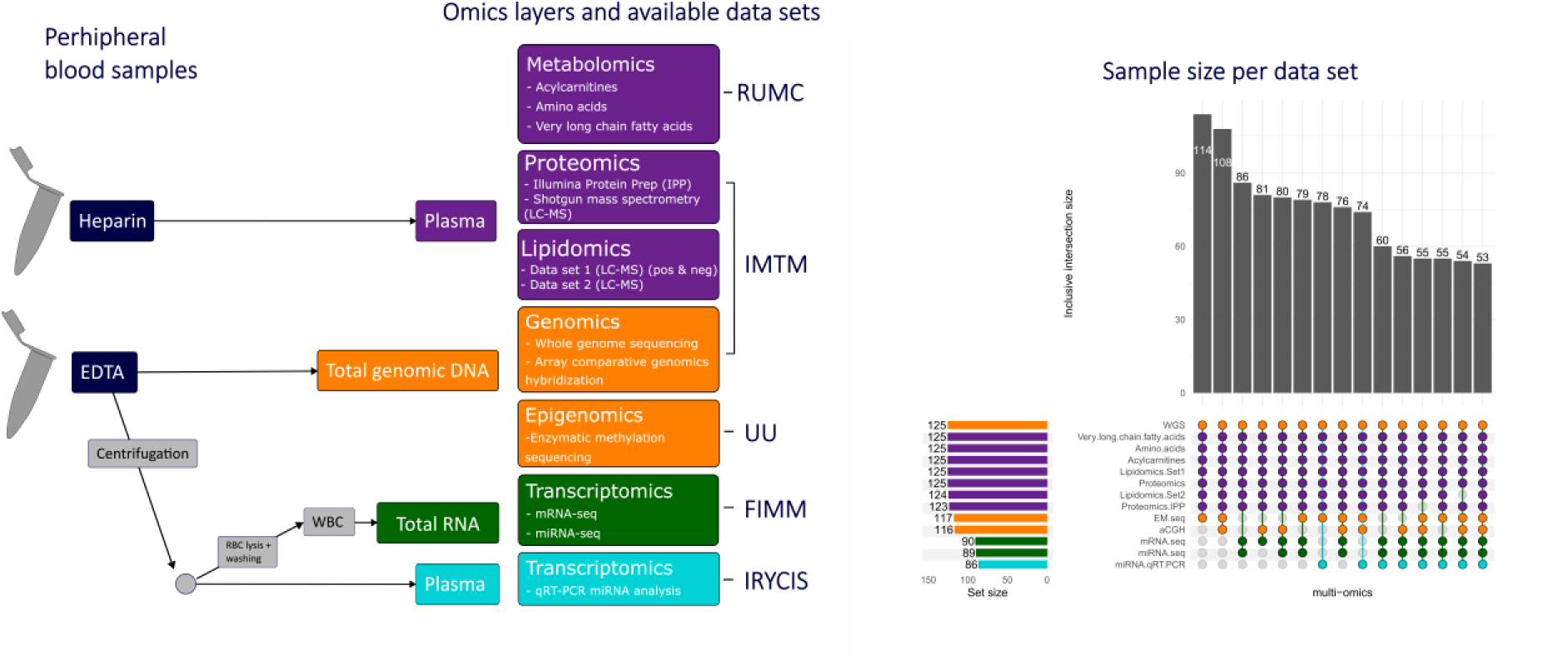
Study design. Different omics layers are analyzed using different platforms, resulting in distinct 13 omics datasets. Sample preparation for the different –omics sets are shown. Colours represent the different blood sample preparations. The EATRIS facilities that analysed the samples are indicated: Radboud University Medical Center (RUMC), Institute of Molecular and Translational Medicine, Olomouc (IMTM), Uppsala university (UU), Institute for Molecular Medicine Finland (FIMM) and Ramon y Cajal Health Research Institute (IRCYS). On the right the sample size per –omics dataset are shown, and the sample intersections of different combinations of –omics layers are plotted.

### Genomic characterization

Since the genetic makeup of individuals may directly affect other omics layers, we first assessed gross chromosomal aberrations. Through the assessment of the whole-genome sequencing, aCGH (Figure 2), and EM-seq data (Suppl Fig S1), we identified one participant with an undiagnosed Klinefelter syndrome (XXY sex chromosomes). This is not surprising given the estimated incidence of 1:500 and because this syndrome does not always lead to evident clinical symptoms. Moreover, aCGH analysis revealed several losses and gains of chromosomal regions in individual participants (Figure 2, Suppl Table 1). We hypothesize that these are the consequences of both genetic heterogeneity in the population and somatic aberrations in clonal expansions of hematopoietic cells^8^. In six genes (*ANKRD36B*, *ANKS1A*, *BTNL3*, *BTNL8*, *DUSP22* and *UGT2B17*), we observed significant differences in the mRNA levels among subjects with gene gains or losses (Suppl Fig S2). Since we regarded these chromosomal aberrations as common genetic variation present in the normal population, we included all subjects in subsequent multi-omics profiling.

**Fig 2.**
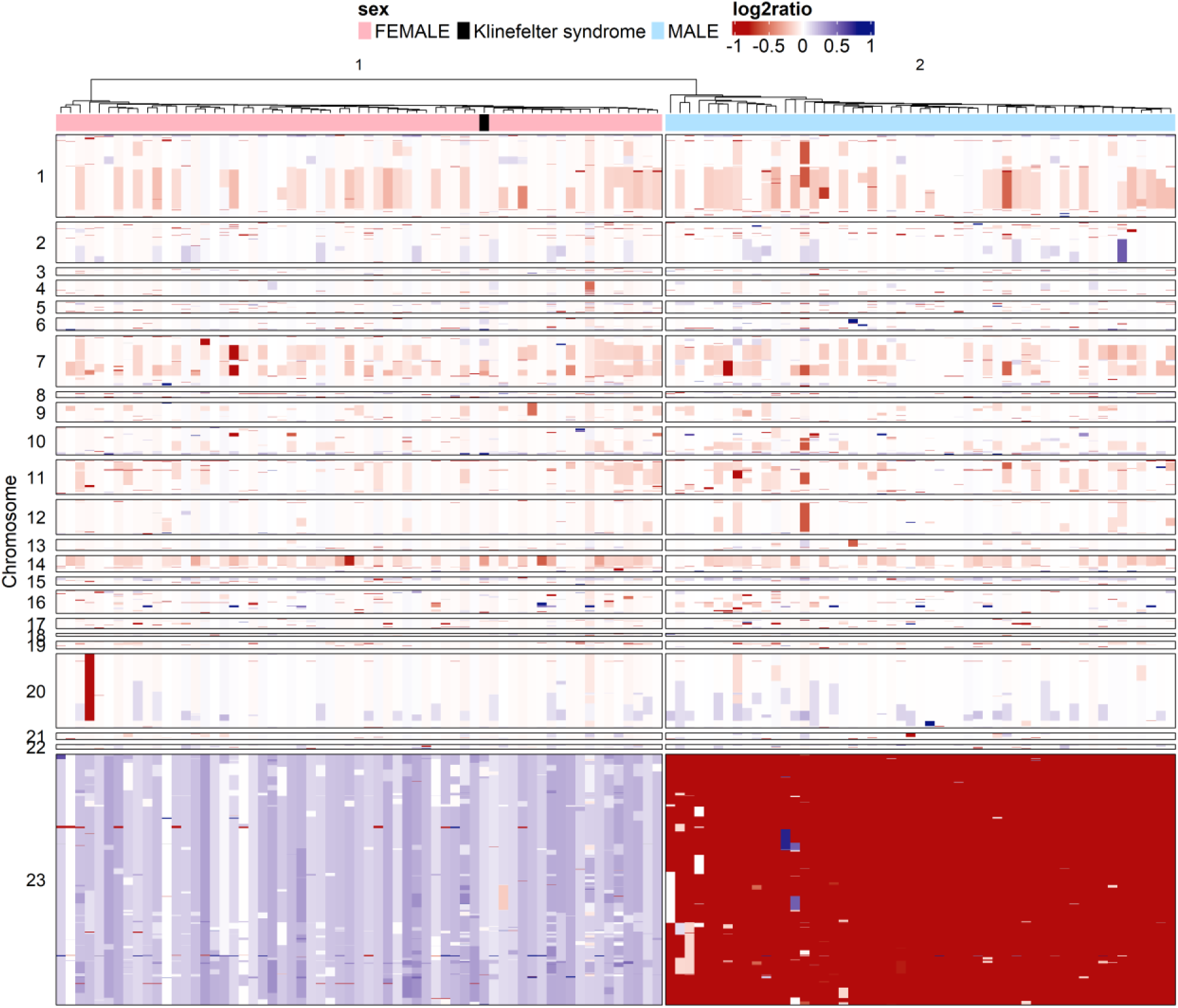
Heatmap of log2 ratio values for genes with LOSS (log2 ratio < -0.5) or GAIN status (log2 ratio > 0.5) grouped by chromosome number. Only 3,707 genes on chromosomes 1-22 and X (labeled as 23) with at least one sample with LOSS/GAIN status presented. The number of selected genes per chromosome can be found in Suppl Table 1. The aCGH data was available for 118 samples, hierarchical clustering (complete agglomeration method, Euclidean distance) was performed in this heatmap.

Furthermore, we examined the effect of single nucleotide polymorphisms (SNPs) overlapping CpG sites on DNA methylation profiles. Notably, we identified over 8,000 SNPs that not only disrupted DNA methylation at the overlapping CpG site, but were also associated with a change in methylation levels of nearby CpG sites (within a 1 kb window) (Suppl Fig S3). For 295 genes closest to these SNP-CpG sites, SNPs were linearly associated with differences in the measured mRNA levels (Suppl Table 2). For two genes, *NQO2*and *TPSB2*, SNPs overlapping with CpG sites were significantly associated with both mRNA and protein levels (Suppl Fig S4). With the high resolution provided by EM-seq, we can assess the effects of SNPs on methylation with high precision. We find that SNPs affect the methylation of larger regions within a CpG island. This may provide a mode-of-action for SNPs affecting gene expression (eQTLs). However, due to our sample size, we did not perform genome-wide association studies and therefore did not exhaustively explore other direct links between the genomics layer and other omics layers, such as eQTLs.

### Variability in omics values

To provide researchers with insight into the number, identity, and abundance of the features detected in the different omics layers, we provide the mean, median, standard deviation and coefficient of variation (CV) for each molecular feature in Suppl Table 3. We stratified these values by sex, because we observed large differences between males and females. Figure 3 illustrates the CV for each feature, sex-stratified and categorized by omics dataset. CVs demonstrated notable differences among the omics datasets. For example, miRNA sequencing exhibited larger variation compared to other data types. This variation can be attributed to differences in leukocyte cell type proportions, ranging from 10%-50% lymphocytes and 40%-80% neutrophils (Suppl Fig S5). Since these differences primarily affect the transcriptomics and epigenomics layers, we adjusted the corresponding sequencing data for white blood cell proportions in all our subsequent analyses (Suppl Fig S6). Moreover, based on the CVs we conclude that platforms within the same omics layer can exhibit considerable differences. For example, the protein levels measured by the IPP platform were more variable (15%) than those measured by the LC-MS proteomics (3%). In addition, variability differed between female and male subjects. For example, females exhibited larger CVs in proteomics-IPP, VLCFAs, and mRNA-seq.

**Fig 3.**
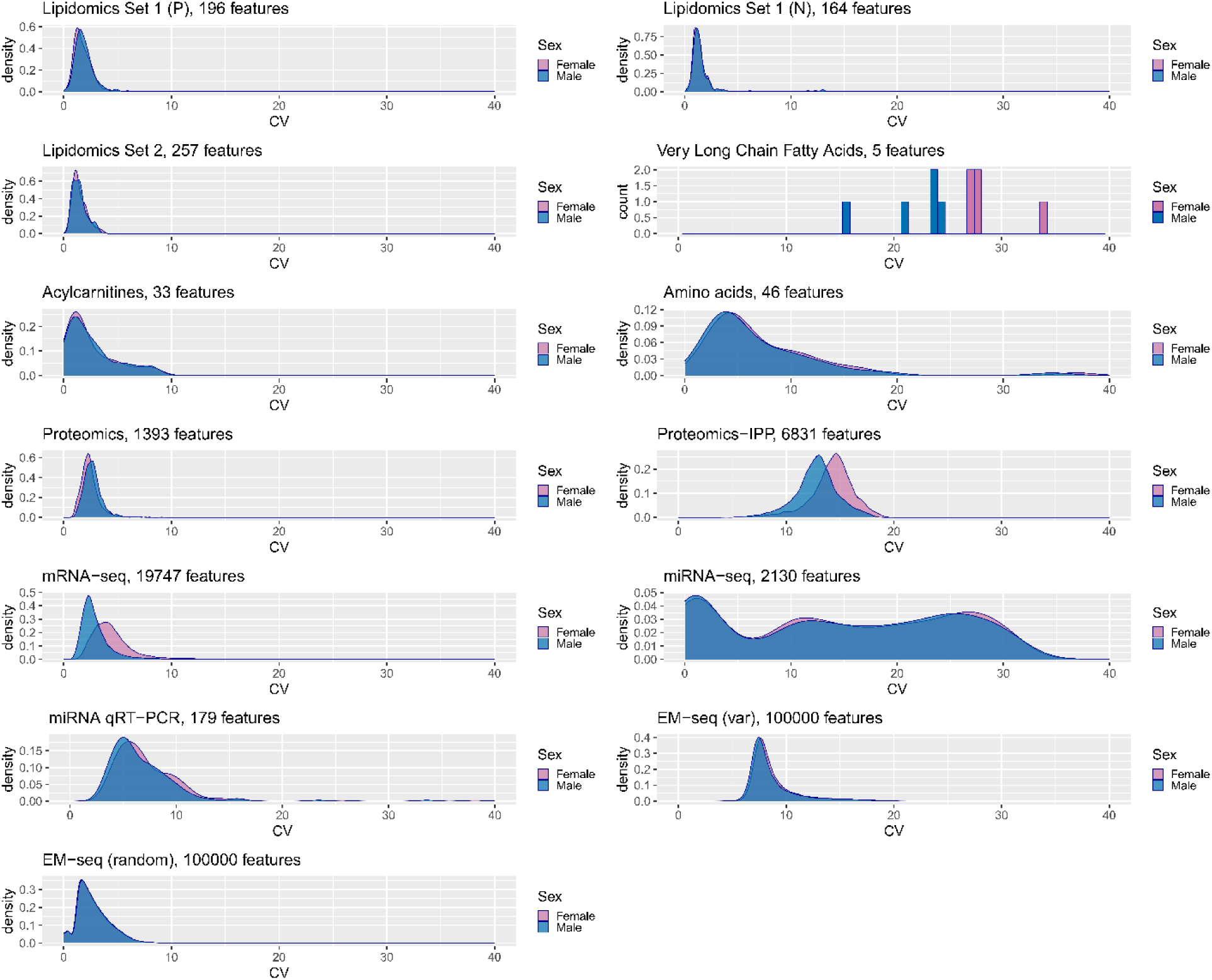
Variability among the –omics data datasets. The Coefficient of Variance (CV) is visualized for every feature in each dataset using density plots. The CV is calculated by taking the ratio of the standard deviation to the mean, and then multiplying the result by 100 to express it as a percentage. CVs are plotted separately for female/male. Missing metabolomics and proteomics data are imputed. A histogram is used or the VLCFA data, as this data contains only 5 features.

As expected, EM-seq revealed that most CpGs were constitutively methylated or non-methylated across healthy individuals. However, there was a subset of CpGs with dynamic methylation in the population. Many of the variable sites overlapped with SNPs, which were excluded from further analysis (Suppl Fig S7). Since the number of CpGs assessed by EM-seq was very large (> 26 million), and that this large number of features may dominate multi-omics analyses, we decided to continue our analysis with the 100,000 most variable CpG sites.

Phosphatidylcholines were found to be the most variable class of lipids. This was particularly true for the negative ionization mode. Pristanic acid and its metabolite phytanic acid were the most variable metabolites in human plasma. This is not surprising since they are heavily influenced by dietary intake^9^. Among the most variable plasma proteins were ApoL1, Alpha-1-antitrypsin, and Immunoglobulin heavy chains, which is possibly due to variable food intake before blood sampling, allelic^10^ and/or antigenic variability of those proteins.

The provided normal ranges for abundance and variation can help to guide power calculations in future biomarker studies and to identify and prioritize biomarkers for disease or lifestyle. Next, we analyzed the omics features in detail by single-omics and multi-omics analysis methods, including Robust Linear Models, Principal Component Analysis and Multi-Omics Factor Analysis.

### Association of single molecular features with phenotypes

We characterized healthy individuals to strengthen the basis of precision medicine studies. Robust linear models (RLM) were applied to discover linear associations between each of the single omics features and a number of phenotypic traits including hematological measurements. This analysis included four different models: linear models with and without sex as a covariate, and two separate models for male and female individuals. Note that the number of molecular features varies widely between the different data types (*e.g.* 100,000 methylation sites *vs.* 5 measured VLCFAs). Figure 4 summarizes the results of these different models by plotting the proportion of statistically significant associations (after multiple testing correction) between omics features and phenotypic traits. All significant associations can be found in Suppl Table 4.

**Fig 4.**
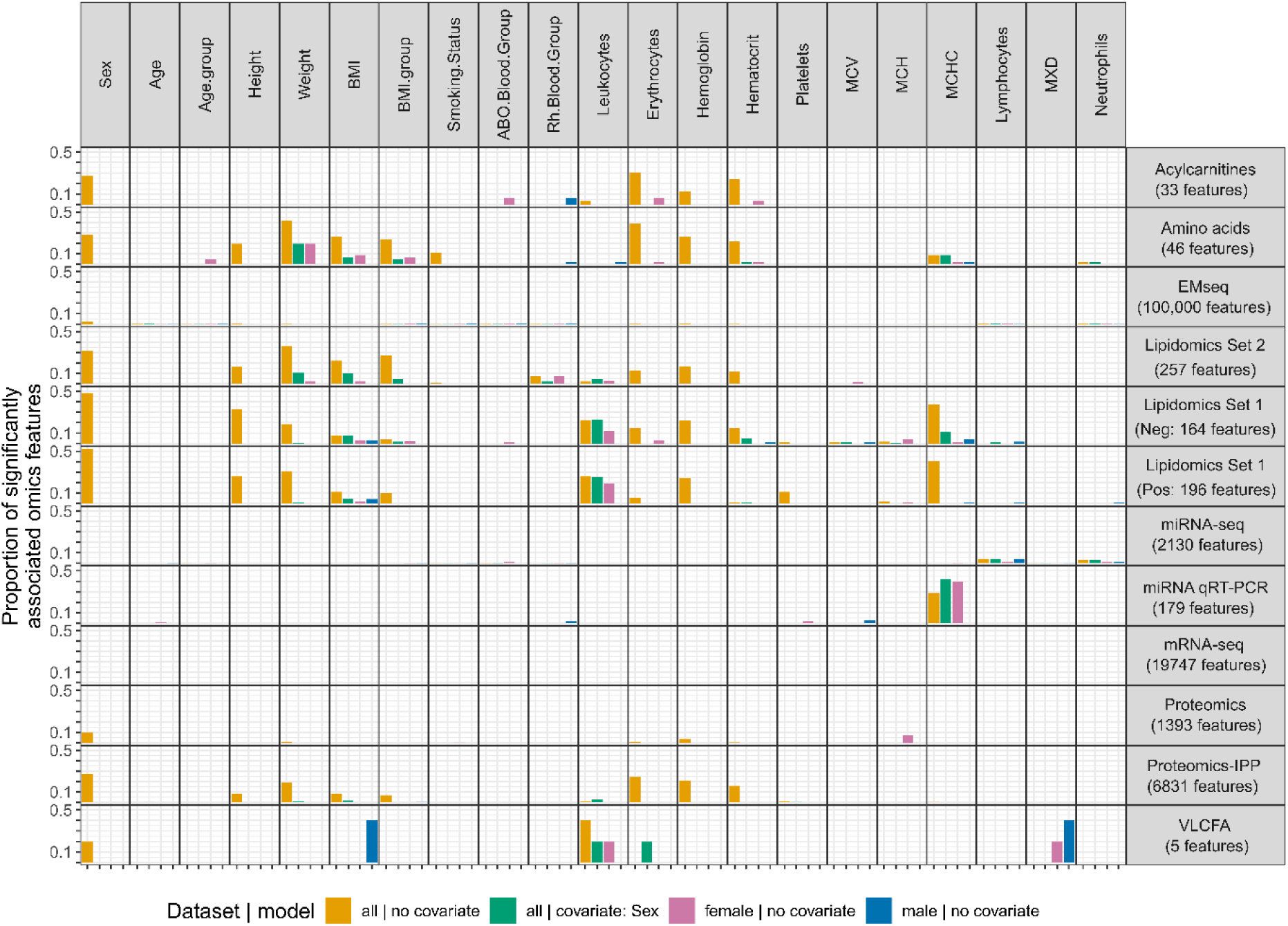
Robust linear models. The proportion of significant associations between -omics features and phenotype variables is depicted, per -omics and per linear model. The maximum value on the y-axis is in all cases 0.5, corresponding to 50% of all features assessed. Numbers of features per omics dataset are indicated on the right. Associations with FDR-corrected p-values below 0.05 are considered significant. Missing values in the metabolomics and proteomics data are imputed, and mRNA-seq, miRNA-seq and EM-seq data are corrected for cell-type compositions. For EM-seq, we only considered the 100,000 most variable CpG sites.

A substantial proportion of targeted metabolomics (amino acids, acylcarnitines), proteomics (IPP) and lipidomics (both datasets) analytes were associated strongly with sex, height, weight, BMI and several blood cell types. For example, root mean squared errors (RMSE) of 0.96 (tiglylcarnitine *∼* blood group) and 0.88 (phosphoetanolamine *∼* sex) were reached.

Associations between these platforms and erythrocyte, hemoglobin and hematocrit levels can be largely explained by sex, as they disappeared when sex was added as a covariate and in the sex-stratified models. Notably, some omics features showed associations only within male or female individuals: VLCFAs (pristanic acid and phytanic acid, Suppl Fig S8) were associated with BMI in male but not in female individuals, whereas pristanic acid and a number of other lipids were associated with leukocyte numbers only in females.

Surprisingly, across all the omics layers, the majority of features did not show significant associations with chronological age. For proteomics (IPP), miRNA-qRT-PCR, miRNA-seq and EM-seq data, less than 1% of the features were associated with age, whereas no significant associations were identified in the other omics datasets. We describe associations with biological age in more detail below.

Despite the correction for estimated cell types, the EM-seq and two miRNA datasets showed significant associations with cell type proportions. In both the miRNA-seq and EM-seq data, hundreds of features were associated with lymphocyte and neutrophil levels, conducted for these data types. In addition, mean corpuscular hemoglobin concentration (MCHC) levels showed linear relationships with a large number of plasma miRNAs measured by qRT-PCR, especially in female samples.

The two proteomics datasets exhibited distinct phenotypic associations. For instance, associations with height and BMI were observed exclusively in the IPP dataset. Consequently, we characterized differences between datasets within one omics layer. This was limited to an overlap of 85 proteins between the proteomics platforms and 52 lipids between the lipidomics datasets. Nonetheless, the majority of shared features exhibited positive correlations (Suppl Figures S9-12). In addition, overlapping lipids were consistently linked to the same phenotypic traits. In summary, while overlap across datasets within one omics layer was limited, shared features displayed similar patterns of variation.

In conclusion, RLMs revealed a large number of significant relations between molecular features and phenotypic traits in all omics layers. Many of the associations can be explained by sex. Hence, balancing sex or conducting sex-stratified analysis in future multi-omics and biomarker studies is essential. The different omics layers exhibited distinct associations with other phenotypic traits, underscoring the added value of multi-omics approaches to analyze biology from different viewpoints.

### Principal component analysis (PCA) on single omics

To reveal multivariate associations with phenotypic traits within single omics data, principal component analysis (PCA) was performed on all individual omics datasets. Following this, the correlations between the first ten principal components and the phenotypic traits were evaluated (Suppl Figures S13-14). Figure 5 summarizes these findings by visualizing, for each omics dataset, the cumulative variation explained by components that correlated significantly with the phenotypic traits.

**Fig 5.**
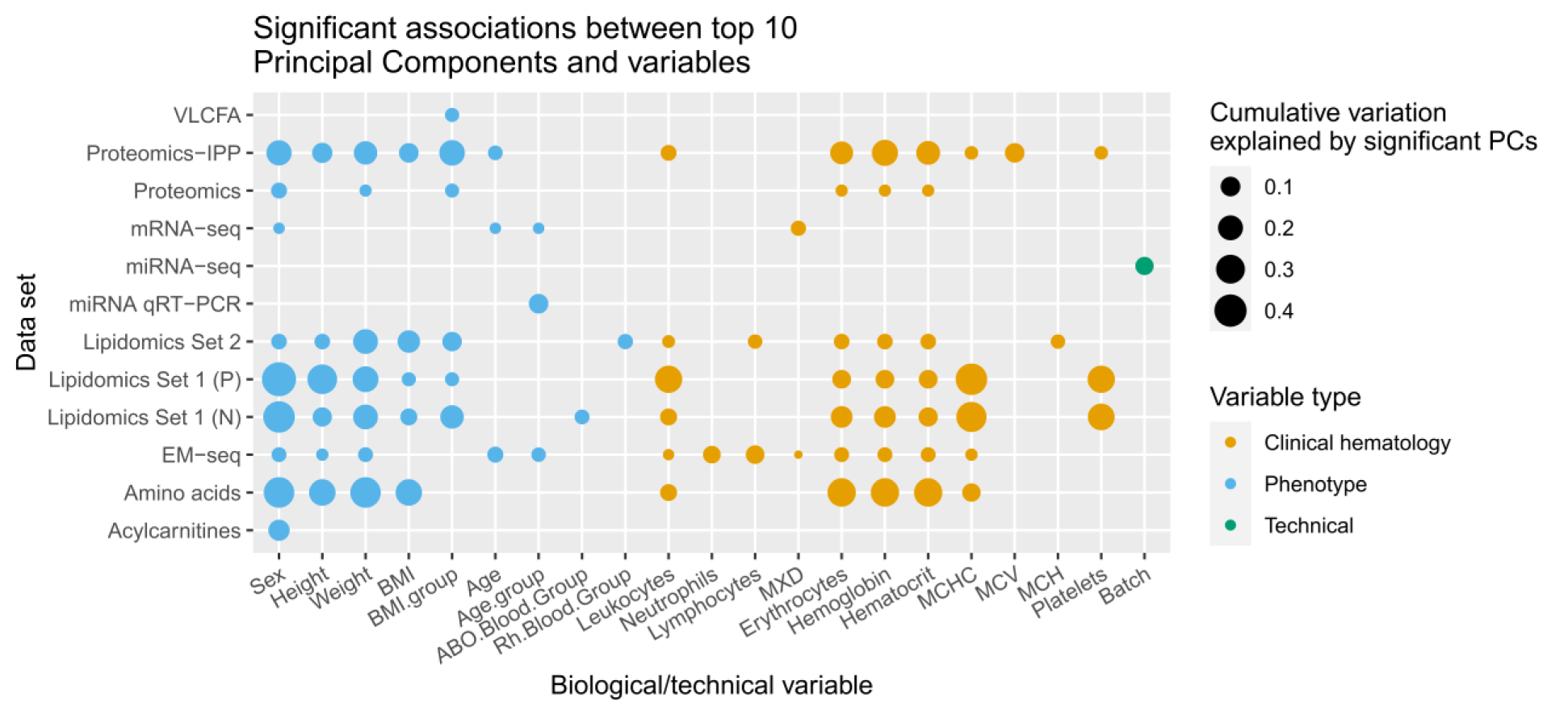
Principal Component Analysis (PCA). Significant associations between one or more PC’s and the variable are visualized for each -omics dataset. Associations with FDR-corrected p-values below 0.05 are considered significant. The size of each dot represents the amount of variation explained by these components combined. As PCA is not able to handle missing values, imputed datasets were used for this analysis.

Patterns observed in the RLM analyses were supported by PCA. For instance, metabolite and lipid levels were mostly associated with sex, body morphology phenotypes and hematological variables. For example, PC1 in the amino acids data associated with weight and BMI, with L-Methionine and L-Leucine as top contributing factors. In contrast to the RLM results, where fewer than 1% of features were associated with phenotypic traits, principal components derived from the EM-seq data showed clear associations, including with chronological age. These components jointly captured up to 7% of the variation in the EM-seq data, highlighting the power of PCA and dimension reduction methods in unveiling hidden patterns in high-dimensional data. Similarly, in mRNA-seq and miRNA plasma levels, associations of PCA with age groups were found, which were not detected by the analysis of single omics features.

We assessed the principal components for influence from technical variation. Since the omics datasets were corrected for batch effects (see Methods), no significant correlations were observed between any of the first 10 principal components and batch, with the exception of the miRNA-seq data (see Discussion). This indicates that batch-adjustment methods have mostly been successful in reducing unwanted technical variation in these omics data. However, for miRNA-seq, results should be interpreted with caution due to a possible residual batch effect.

The PCA results revealed consistent patterns across datasets within the same omics layers. For example, the top components derived from the lipidomics datasets both showed associations with sex-related phenotypes such as weight, height and cell type levels. The proteomics components were linked to the same phenotypes. However, the proteomics (IPP) data were also associated with other sex-dependent traits such as height, weight and hematocrit.

In summary, the PCA of individual omics data confirmed the findings from the linear models. Additionally, it revealed age-related associations that were not detectable when analyzing omics features separately. This indicates the added value of multi-variate analyses in complex multi-omics datasets.

### Multi-omics factor analysis (MOFA)

To uncover joint sources of variation across different omics layers, we applied the unsupervised multi-omics factor analysis (MOFA) method. MOFA learns latent representations of the multi-omics data and summarizes them as MOFA factors, much like principal components for single–omics data, but integrating information across multiple data modalities^11^. We selected only lipidomics data acquired in a single ionization mode (positive) to avoid domination by these highly correlated datasets.

Figure 6A depicts the twelve latent dimensions uncovered by MOFA and quantifies the contributions of the individual omics datasets to these MOFA factors. This figure clearly illustrates how the MOFA factors comprise information from multiple omics levels. For example, factors 1 and 6 captured variance present in metabolomics, lipidomics, epigenomics and proteomics data.

To facilitate the biological interpretation of the MOFA factors, we correlated the factor scores with the continuous phenotype traits (Fig 6B) and performed pathway enrichment analysis (Suppl Figures S15-26) on the weights (Suppl Table 5) of the mRNAs, proteins and metabolites on the MOFA factors. Several significant correlations were identified between MOFA factors and continuous (Figure 6B) and categorical (Figure 6C-E, Suppl Fig S27) phenotypic traits. For instance, factors 1 and 6 were associated with sex, height, weight and blood cell type levels. As the scores for factors 1 and 6 were significantly different between males and females, the associations with the other 3 variables may also be driven by sex. Interestingly, the Klinefelter individual was most similar to females in factor 1 and 6, but appeared to be an outlier in factors 9 and 11, suggesting that this individual is different from XX and XY individuals in multiple molecular aspects.

**Fig 6.**
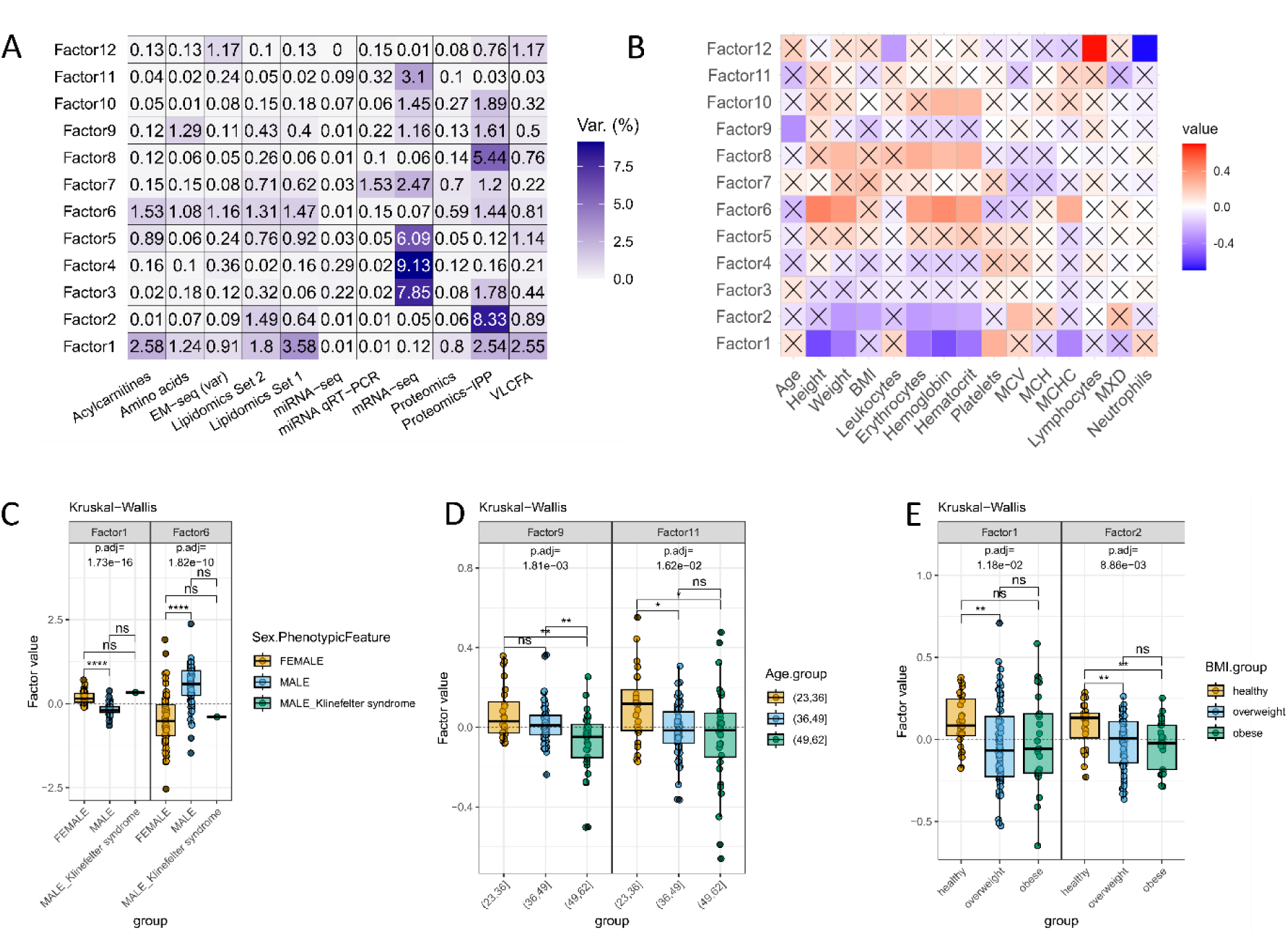
Multi-Omics Factor Analysis (MOFA) results. A) Explained variance across the –omics datasets for MOFA factors 1-10. B) Heatmap showing correlations between the MOFA factors and phenotypic variables; significant associations are open squares (not marked with an X). Comparative analysis of the values of the MOFA factors associated with sex (C), age (D) and BMI (E) groups. Significance was tested using Wilcoxon for sex. Kruskal-Wallis was used for age and BMI, which were post-hoc tested using a two-sided Dunn test. All p-values were corrected using Benjamini-Hochberg’s false discovery rate (FDR).

Factor 2 was dominated by the proteomics (IPP) dataset, with smaller contributions from the lipidomics datasets. This factor seems to reflect variation related to BMI differences (Fig 6E). Moreover, the protein feature weights on factor 2 were associated with metabolism, insulin regulation and inflammatory pathways (Suppl Fig S16, Suppl Table 6). Many of these proteins were low-abundant molecules such as cytokines and chemokines, which were not detected by the LC-MS-based proteomics.

Although MOFA is designed to capture variance across different data sources, factors 3, 4 and 5 represented mainly contributions from the mRNA-seq data. The MOFA sample scores on these three factors were not correlated with the phenotypic traits, but pathway enrichment analysis revealed several significant associations with immune-related biological pathways (Suppl Table 6). Although factors 3, 4 and 5 were not significantly associated with the overall percentage of lymphocytes and neutrophils, we hypothesized that this variance is due to the cell type proportions that could not be compensated by the cell type correction. This hypothesis was supported by the strong involvement of cell type-specific transcripts in these factors (Suppl figures S28-31). B-cell specific genes had large negative weights on factor 4, whereas monocyte-specific genes demonstrated large positive weights on factor 5. Furthermore, granulocyte-specific mRNAs and genes coding for proteins of the large and small subunit of the ribosome had strong contributions to factor 3. Ribosomal protein gene expression differs between cell types in the blood^12^. Thus, the cell type correction that we applied may have worked well for the major cell types, based on cell-precursors, but may have been less efficient for corrections of more granular differences in cell types with a common cell precursor, such as the proportion of B- and T-cells.

The mRNA and protein feature weights from factor 4 were associated with immune-related pathways, for example the complement pathway (Suppl Figures S18). Since the proteomics and mRNA-seq data were derived from different sample types (plasma vs. white blood cells (WBCs)), this variation is less likely to be affected by the cell type proportions.

Factor 9 was indicative of the aging process. Factor 9 primarily accounted for variations observed in amino acids, IPP-proteomics and mRNA-seq data, exhibiting a negative correlation with chronological age. Accordingly, sample scores on factor 9 differed significantly between the age groups (Fig 6C). Additionally, the mRNA-seq and proteomics feature weights were enriched in aging-associated pathways, including chemokine signaling^13^, cytokine activity^14^, and JAK-STAT signaling^15,16^. High contributing mRNA features on these factors are *CCR7*, *IRAK2*, *STAT5A* and *CXCR3* (Suppl Table 6). Remarkably, the two strongest contributing miRNAs (hsa-miR-139-5p and hsa-miR-99a-5p) have already been linked to chronological age or biological processes involved in cell senescence^17–19^. A more detailed analysis of the association between methylation levels and aging is given below.

Finally, MOFA factors did not associate with the chromosomal arrangements observed in the aCGH data (Suppl Fig S32), suggesting that they did not have a major impact on the multi-omics profiles except from the XXY configuration in the Klinefelter individual.

In conclusion, MOFA was able to discern shared variation across the omics layers, which could be linked to different phenotypes and biological processes. While most sex effects were captured by the first and sixth MOFA factor, other MOFA factors reflected non-sex dependent biological variation that was more difficult to detect in single omics analyses.

### Results from other multi-omics data integration methods

In addition to MOFA, we analyzed the multi-omics data with other data integration methods and the results mostly confirmed the results from MOFA and single-omics analyses. We applied consensus independent component analysis (consICA), a method and R package we recently developed to disentangle technical biases and important molecular signals, highlighting active biological processes and cell subpopulations^20^. We applied consICA separately to EM-seq, mRNA-seq, miRNA-seq, LC-MS proteomics, metabolomics, lipidomics, with the latter two combined as single data modality. ICA identified 20 independent components, that we integrated in downstream analyses. In line with the PCA and MOFA results, strong associations with sex were identified. Specifically, ICA component g1 from the mRNA-seq data (Fig. 7A), as well as components from the EM-seq (e5) and metabolomics data (m17), showed significant associations with sex (Fig. 7D, Fig. 8). Additionally, EM-seq component e20 allowed precise recovery of lymphocyte/monocyte abundances through linear regression (Fig 7B), with *R*^2^ = 0.80 and *R*^2^ = 0.76 for lymphocytes and neutrophils, respectively.

**Fig 7.**
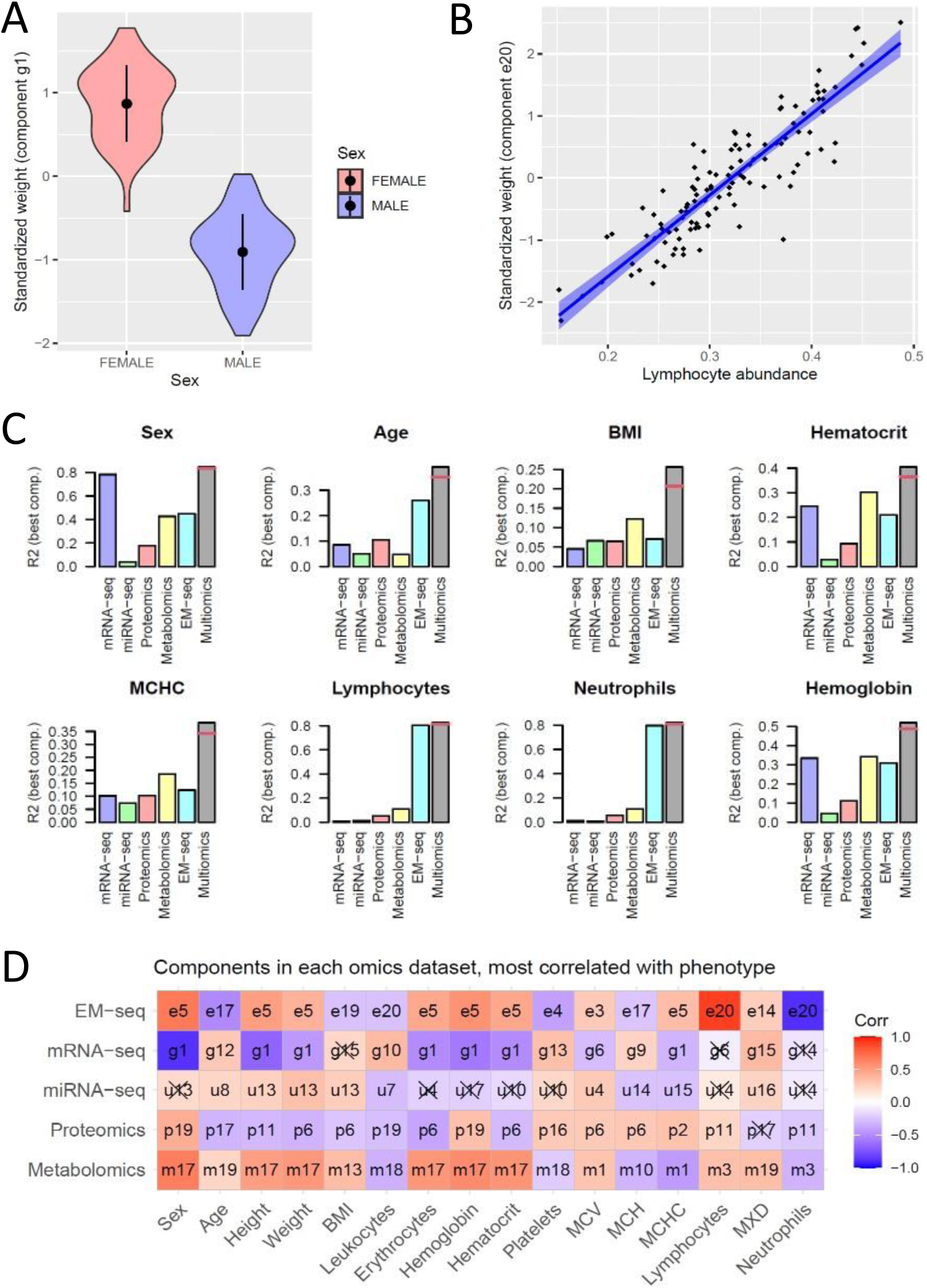
consICA on various -omics layers predicts phenotypes. (A) Violin plot of the weights of mRNA ICA component g1 for males and females (p-value: 4.7e-32; t-test). (B) The weights of the DNA methylation component e20 were strongly linked to lymphocyte abundance in the samples. (C) The top *R*^2^ observed between experimental variables and independent components of each -omics dataset separately, and their combination into a multivariable linear model (“Multi-omics” bar, red lines represent adjusted *R*^2^). (D) The top correlated ICA component for each combination of omics and phenotypic variables. Non-significant correlations are marked by X (FDR-corrected p-values of the visualized correlations).

**Fig 8.**
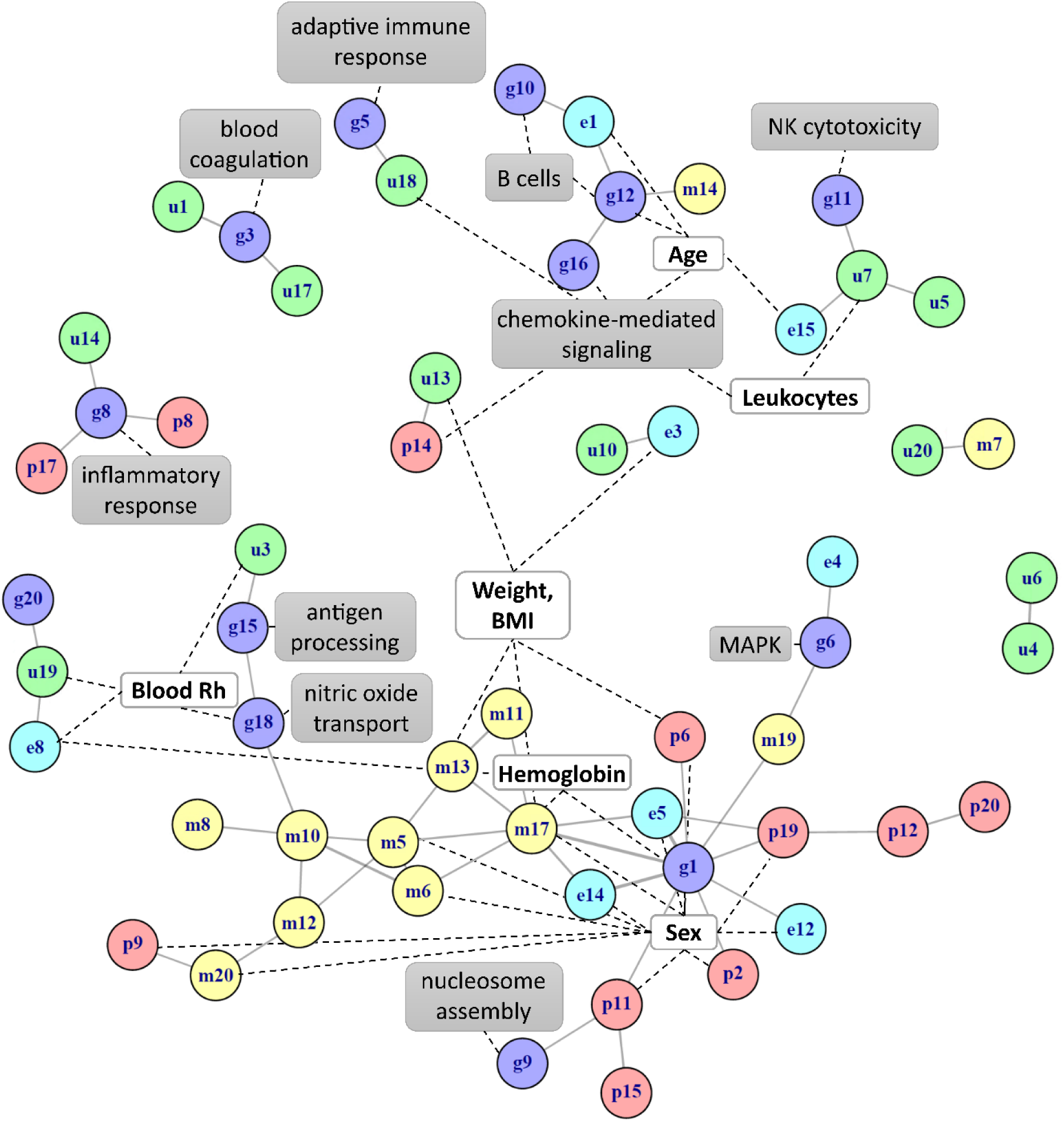
Integration of ICA results as a correlation-based graph. The nodes represent components from epigenomics (e#), transcriptomics (g#), microRNA-omics (u#), proteomics (p#), and metabolomics (m#). The edges show components with correlated weights (*R*^2^ > 0.1, p-value < 0.001). Additional annotation based on clinical data (bold, white box) or the most important gene sets (grey box) is shown.

To demonstrate the added value of multi-omics data integration, we built linear regression models to predict phenotypes using the components’ weights. For the majority of clinical variables (e.g., BMI, age, hematocrit, MCHC, and hemoglobin levels), the multi-omics components explained more of the variance than the individual–omics components (Fig. 7C), demonstrating the complementarity of the information present in the different omics datasets. As expected from the results described above, white blood cell proportions were accurately estimated from methylation components, with minimal additional contribution from other omics components. (Fig. 7C).

By correlating the weights of single-omics consICA results, we constructed a network of multi-omics components and identified the biological functions represented by components from different omics dataset (Fig. 8, Suppl Table 7). The multi-omics components were linked to general blood characteristics, such as blood coagulation, immune cell function, and the blood Rhesus factor (Blood Rh), as well as more specific biological functions, including nucleosome assembly, mitogen-activated protein kinase (MAPK) signaling, and nitric oxide transport (Fig. 8).

As another way to investigate the complementarity of the different omics datasets, we clustered the samples using similarity network fusion (SNF)^21^, a method that fuses multiple sample similarity networks constructed from single omics data into one. Here, we showed that clustering based on multiple omics datasets did not demonstrate stronger clustering of samples than single omics datasets (Suppl Fig S33). Furthermore, partial least squares 2 (PLS2)^22^ analysis demonstrated the strongest covariance between the EMseq data and other omics datasets, which were likely driven by sex (Suppl Fig S34). Moreover, multiple integration methods (MOFA, SNF, and PLS2) revealed strong correlations between VLCFA and lipidomics datasets, likely because VLCFAs represent a specific lipid class.

### Biomarkers for biological age

Methylation is often used to assess biological age^23^. However, other omics layers may reveal biological pathways that are affected by methylation or provide a complementary view on biological aging, thus yielding a more comprehensive and potentially improved assessment of biological age. Therefore, we first predicted the participants’ biological age using our high-resolution DNA methylation dataset. We then correlated these biological age estimates with other omics layers to explore more connections between the omics layers. Notably, these epigenetic clocks were developed on methylation array data, whereas we generated EM-seq data. Therefore, we first demonstrated the performance of the most popular epigenetic age predictors on our cohort and our EM-seq based data. These epigenetic clock predictors will be referred to as: best linear unbiased prediction (BLUP)^24^, Bayesian neural networks (BNN)^25^, elastic net (EN)^24^, Hannum^26^, Horvath^27^, Levine^28^ and skin-Horvath^29^. Age predicted with BLUP and EN clocks was most strongly associated with the chronological age of the participants of our cohort (Fig 9A). The high correlations (*R*^2^ of 0.90 and 0.81, respectively) are remarkable given the differences between the methylation profiling platforms and the different geographical origin of our population. The other age predictors, including Levine, performed significantly worse, suggesting they are less generalizable across profiling platforms and/or population. In line with the observed associations between MOFA factors 9 and chronological age, we observed significant associations between this factor and predicted age based on the BLUP but not Levine clock (Fig 9B). This confirmed that BLUP was superior in predicting chronological age in our cohort.

**Fig 9.**
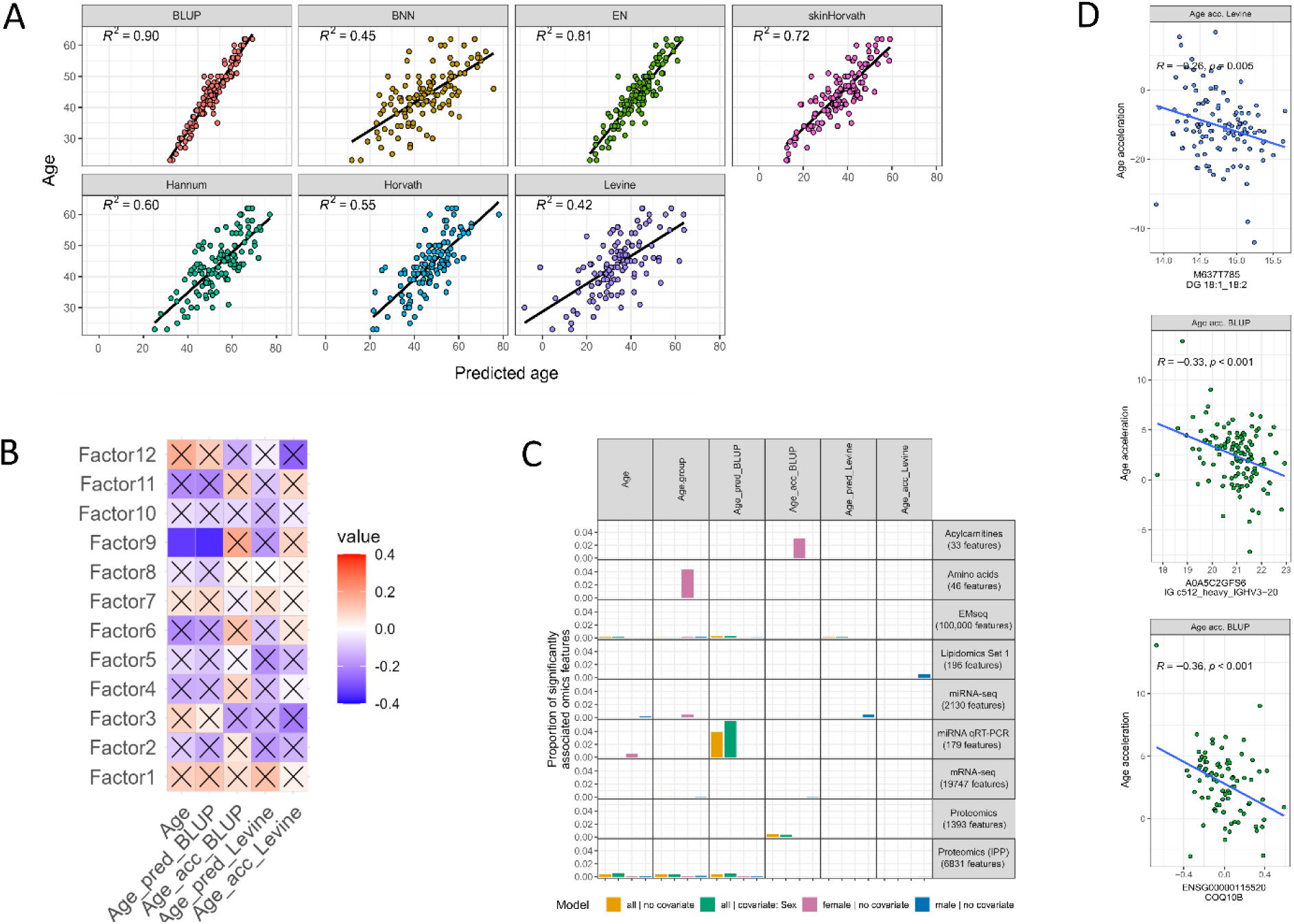
Associations with biological age. A) Performance assessment of the different epigenetic clocks, by comparing them with the chronological age. B) Robust linear model with the age prediction and age acceleration (biological age - chronological age) from BLUP and Levin clocks as outcomes. C) MOFA factor correlations to the same age prediction and age acceleration. D) Individual features correlating with age acceleration.

We then tested for linear associations between BLUP- and Levine-predicted age and all omics features (Fig 9C). Associations of omics features with chronological age were often recapitulated by associations with BLUP-predicted age, which was expected due to the high correlation of chronological age with BLUP-predicted age (Suppl Fig S35). However, we also observed a considerable fraction of miRNAs in the plasma with more significant associations with BLUP-predicted age than with chronological age (Suppl Table 4). Furthermore, associations with the Levine-predicted age were mostly not overlapping associations with chronological age, suggesting that the Levine-predicted age reflects distinct molecular patterns of aging.

To identify candidate omics biomarkers for biological aging, we calculated the age acceleration, *i.e.* the difference between chronological age and age predicted by BLUP and Levine, after adjusting for the calculated cell type composition. Notably, lipids like CDP-diacylglycerol, transcripts such as *COQ10B*, *SH3GLB1*, and *MFSD14A* and diverse immunoglobulin V-region peptides were significantly associated with accelerated age but not with chronological age (Fig 9D, Suppl Figures S36-S38). We observed only weak associations between age acceleration and phenotypic traits such as blood group and smoking status (Suppl Fig S36), making it unlikely that associations with age acceleration are due to confounding of other biological factors. The identified molecules may thus represent candidate biomarkers of biological processes that contribute to the aging process throughout the lifespan or biomarkers of accelerated aging, relevant for pathological contexts like degenerative diseases or cancer.

## Discussion

We reported one of the most comprehensive multi-omics studies in a cohort of healthy volunteers performed so far, integrating six omics layers derived from twelve platforms applied to 127 blood samples, surpassing the scope of other large-scale studies^30–33^. Our dataset can be used as a benchmark, providing normal ranges for abundance and variation of tens of thousands of omics features in the healthy population and guiding the design of reproducible omics studies investigating specific (disease) phenotypes and pharmaceutical or lifestyle interventions. Our study enables power analyses, sample size estimations and sex composition balancing for future multi-omics studies. Moreover, our dataset can be used to benchmark computational methods for multi-omics data integration, as we did in this study. The best-performing multi-omics integration techniques would optimize the number of identified molecular associations with phenotypes that are known to influence molecular omics levels such as sex, age and BMI.

With our benchmark study, we set out to answer the following questions guiding the designs of future multi-omics studies: **Question 1: What are the challenges in multi-center multi-omics studies? Answer:** From our study, it is evident that running a multi-omics cohort analysis in a controlled manner in five different laboratories with highly standardized procedures (available through the EATRIS multi-omics toolbox: motbx.eatris.eu) is feasible. In establishing reference protocols and materials for multi-omics studies^34–36^, we faced several challenges that needed to be addressed when implementing these techniques in multi-center omics analyses. For example, designing multi-omics studies with respect to sample sizes can be challenging, also due to costs. Our dataset provides a valuable resource for estimating the minimum required sample size to draw statistically valid conclusions. For instance, power calculations can be performed using the tables of normal ranges and abundances (Suppl Table 3), as well as the effect sizes and standard deviations derived from linear associations with phenotypic traits (Suppl Table 4).^37^ Another critical challenge is technical noise, which introduces sources of unwanted variation that can obscure true biological signals. These include differences in reagent batch quality, instrument drift, and variability between operators.^7,38–40^. If not addressed either through experimental planning or stringent execution of the study along with careful documentation of experimental steps, such unwanted noise will lead to misinterpretation of results. Experimental design strategies can help alleviate the influence of technical noise. These include clinical trial design with appropriate inclusion and exclusion criteria, standardization of biospecimen sampling, planning of the pre-analytical phase, inclusion of reference material, and stratification of samples across experimental batches. Finally, statistical modeling can reduce unwanted variation^41^. For instance, we concluded that batch correction with ComBat, originally developed for mRNA-seq data, worked equally well for other omics datasets (Suppl Figures S39-47). Moreover, Combat allows inclusion of covariates, helping to preserve biologically relevant signals while reducing the risk of false positive associations. Therefore, we included sex as a covariate to avoid eliminating sex-dependent effects from the data. However, the partial confounding of batch and sex in some omics datasets should have been avoided by running equal numbers of male and female samples per batch. Lastly, one major challenge is missing assays due to quality issues and missing values due to detection problems. These missing values complicate the analysis and integration of omics data. Data imputation is often necessary, in particular for LC-MS-based methods (see Suppl Figures S48-50).

**Question 2: Which biological and technical confounders should be considered? Answer:** Sex-dependent variation was observed through all omics layers, with strongest effects observed in lipidomics and metabolomics, as was also discovered by Mittelstrass et al.^42^ and Beyene et al.^43^ This suggests that future studies where sex is not a study parameter, irrespective of the omics technique applied, should be balanced in the number of females and males included. Actually, studies may benefit from analyzing the females and males separately, because many diseases progress differently between sexes^44,45^. Separate analysis can reveal more specific or better prognostic biomarkers for each sex. Cell type composition differences were a major confounder for the omics techniques performed on blood cells but, as expected,^46^, not so much for plasma-based assays.^47^ Our MOFA analyses and linear associations with cellular transcriptomics levels demonstrated that there were residual effects from differences in cell type composition, despite our efforts to correct based on the cell type proportions estimated from the EM-seq data. This can be partially explained by the different sample processing per omics dataset (Fig 1). Moreover, we did not consider low abundant subtypes of leukocytes in the estimation of cell type proportions. We suggest using independent analytical technologies such as multiparametric flow cytometry or CyTOF for measuring instead of inferring cell type proportions.

**Question 3: How are the different omics layers related and connected, which –omics layers capture complementary biological information, and which are largely redundant? Answer:** From the different multi-omics integration techniques performed, we conclude that the omics layers are only weakly connected and contain mostly complementary information. Nevertheless, MOFA and ICA both identified latent dimensions across the omics layers that reflect distinct sources of biological variation (sex, age and cell-type proportions). Moreover, analysis of genetic variants from WGS and aCGH revealed specific links between the genomics layer and other omics layers (Suppl Figures S2-4, S28). The strongest associations were observed between SNP-CpG pairs and corresponding mRNA levels. These findings could explain some of the shared variation identified through integration methods. However, other omics layers did not strongly correlate with each other nor show significant associations with similar biological pathways. Even protein and RNA levels of the same genes were only weakly correlated (Suppl Fig S51). This is partially because they profiled molecules from different sources: EM-seq, RNA-seq and miRNA-seq were performed on white blood cells, whereas peripheral tissues - along with blood cells - contribute lipids, proteins and metabolites to circulation. We therefore recommend selecting omics layers in future omics studies based on specific biological hypotheses, taking into account the relevant tissues, cell types and their secretions.

We compared our results to those from other large multi-omics cohort studies, Swedish CArdioPulmonary bioImage Study (SCAPIS) in Sweden^48–50^, Biobanking and Biomolecular Resources Research Infrastructure - Netherlands (BBMRI-NL) in the Netherlands^51,52^, integrative Personal Omics Profiling (iPOP) in the USA^53,54^ and the UK Biobank^55,56^. This revealed consistencies, complementarities and potential discrepancies.

The SCAPIS and iPOP studies focused on longitudinal fluctuations of omics features within an individual, whereas we focused on the cross-sectional variation of omics features across individuals. SCAPIS ranked proteins as most stable omics features, followed by lipids and metabolites. However, in our study proteomics could be as variable as other omics layers, depending on the platform. Furthermore, the iPOP study identified a considerably higher percentage of proteins with seasonal fluctuations than transcripts or metabolites. Analytes with large (environmentally induced) fluctuations and large variation between individuals are not likely to serve as reliable biomarkers and including them in biomarker panels may lead to reduced power. We therefore recommend consulting our tables with CVs when designing (disease) biomarker assays by deprioritizing analytes with large CVs.

Associations between omics features and phenotypes in other studies were re-discovered in our study. For example, some of the highlighted sex-associations in the SCAPIS study (L-leucine, protein Prokineticin-1 and mRNAs *ZFX* and *KDM6A*) were also found in our study. However, we observed different trends in the fraction of omics features associated with sex (SCAPIS: transcripts > proteins > metabolites > lipids; our study: lipids > metabolites > proteins > transcripts). Similar inconsistencies were found for associations with BMI (SCAPIS: proteins > transcripts > metabolites > lipids; our study: metabolites > lipids > proteins). This may be attributed to age differences between the two cohorts, with our Czech cohort on average younger than the Swedish SCAPIS cohort, to other differences between the cohorts and the use of targeted antibody-based proteomics techniques *vs.* LC-MS-based and somamer-binding-based proteomics. In addition, our results confirmed some of the strongest associations of plasma protein levels with sex, age and weight found in the UK Biobank resource^56^, which used a different platform for detecting protein levels (Olink). For example, sex hormone-binding globulin (I3L145) and Apolipoprotein F (Q13103) strongly associated with both weight and sex in both cohorts.

In line with results from the iPOP study, we found a number of omics features other than methylation sites associated with age. However, the metabolite analytes, transcripts, and proteins identified in our study did not overlap with those reported in the iPOP study.

Our two miRNA profiling platforms, miRNA sequencing on white blood cells and qRT-PCR profiling of plasma, have not been applied in SCAPIS, the Biobank-based Integrative Omics Study (BIOS) from BBMRI-NL and iPOP. The two miRNA profiling methods provide complementary data, because the miRNAs in the plasma usually originate from peripheral tissues (bound to lipoproteins^57^, and/or extracellular vesicles^58^), with only a minor contribution from blood cells. Still, we observed that miRNA sequencing on white blood cells and qRT-PCR on plasma demonstrated co-variance (*i.e.* a correlation of 0.83 in PLS2).

Methylation profiling was only done in BIOS, and not in SCAPIS or iPOP. However, connections between methylation and miRNA levels, metabolomics, proteomics or lipidomics were not established in BIOS. On the contrary, BIOS detected many links between genetic variants and mRNA expression^51^ and CpG methylation^52^ (expression and methylation quantitative trait loci) that we could not identify due to our smaller sample size.

Most notably, we identified candidate miRNA, mRNA, protein, metabolite, and lipid biomarkers for biological age and age acceleration predicted from methylation data. These biomarkers underlie many relevant diseases and have not been observed before.

In conclusion, the total overlap of significant associations was limited, compared to the large number of omics features. This highlights the need for more reference materials and standardized guidelines on multi-omics studies for assuring their reproducibility and further implementation in clinical practice.

### Biologial age

Extensive methylation profiling using EM-seq is unique to our cohort but is remarkably consistent with the microarray-based methylation profiling performed for the BLUP age predictor^24^. The correlation of the BLUP-predicted age with the chronological age in our cohort reaches 0.95, with a slope close to 1. This is even more remarkable since the BLUP predictor is based on methylation array measurements, whereas we used a sequencing-based method. This confirms the superior precision and accuracy of the BLUP predictor over other biological age predictors and the high quality and accuracy of our EM-seq methylation level assessment. Obviously, the much higher resolution and coverage of CpG sites in the EM-seq platform provide more opportunities for fine mapping of causal methylation sites and methylation quantitative trait loci over more commonly used array-based technologies. The age predictions using Levine^28^ correlated less with the chronological age. This is not surprising, since this epigenetic clock has been specifically trained on a surrogate healthy age, determined based on clinical measurements. Therefore, age predictions based on Levine are more indicative of a healthy lifespan.

Biological age can also be estimated using other omics data. For instance, proteomic aging clocks have demonstrated predictive power of age-specific mortality and disease^59^, similar to the Levine epigenetic clock. Moreover, different omics age predictors can be integrated into a single multi-omics age predictor, which could be more accurate in predicting healthy lifespan and accelerated aging^60^. Such a predictor could potentially be enriched with the omics features associated with accelerated age in this study (Fig 9).

Interestingly, transcripts that we associated with accelerated aging, such as *COQ10B*, *SH3GLB1*, and *MFSD14A*, are closely linked to mitochondrial functions. These findings reinforce the well-established role of mitochondria in the aging process, as previously reported in the literature^61,62^.

Some of the miRNAs associated with aging in our study have also been linked to age-dependent disease progression in other studies. However, it is important to note that miRNA profiles associated with age or disease may be tissue-dependent^63^ and may not always be reflected in blood. Several miRNAs were significantly associated with age in our models. In the qRT-PCR data, predicted age correlated with miR-223-5p, which has also been linked to liver fibrosis.^64^And BLUP-predicted age was associated with hsa-miR-425-3p, hsa-miR-191-5p, and hsa-miR-485-3p, which are previously implicated in Alzheimer’s disease^65,66^. In miRNA-sequencing data, chronological age and Levine-predicted age were respectively associated with miR-6762-5p, miR-20a-3p, and hsa-miR-6876-3p, each also connected to Alzheimer’s^65,67^. Furthermore, in the age-associated MOFA factor 9, the top contributing miRNAs were also previously associated with aging. For instance, miR-17-5p has been linked to cellular senescence processes, including apoptosis and cell cycle regulation^68–70^. Similarly, miR-139 has been shown to influence senescence in hepatocellular carcinoma^71^, and miR-221-3p has been implicated in brain aging and age-related diseases^72^.

Conversely, none of the 25 miRNAs associated with chronological age identified by Huan *et al*. were associated with age in our study. This is not surprising because Huan *et al*.^73^ analyzed miRNAs in whole blood, where platelet levels can alter the miRNA profiles. Nevertheless, we linked a number of these miRNAs to neutrophil and lymphocyte levels. These age associations may be driven by age-dependent variation in blood cell composition^74^.

### Limitations

Sample QC steps in large-scale omics studies are essential. This includes sample concordance checks to eliminate possible sample swaps that can affect the research output. While sequencing-based methods allow for such sample checks (sex-matching, SNP-profiling, see Methods), other data types such as LC-MS-based metabolomics and lipidomics cannot be used for such sample checks. Consequently, the risk of sample swaps in the final multi-omics dataset cannot be fully eliminated.

We analyzed omics data from 125 healthy individuals, although not all datasets were available for every sample. The miRNA qRT-PCR data had the lowest coverage, with 86 samples (Figure 1). While this is less critical for single-omics analyses, integration across omics layers is limited by the reduced number of complete data available for fewer than 60 individuals. This may partly explain the modest contribution of miRNA data in MOFA, and suggests that higher sample coverage could further enhance the detection of cross-omics relationships.

We applied cell composition correction methods that also reduce sex-related variation as the number of specific cell types differs between male and female samples (Suppl Fig S5). Nevertheless, sex was the most dominant source of variation in all our analyses, highlighting the strong influence of sex on omics profiles We recognize that we are faced with a huge multiple-testing problem in these multi-omics studies. While we applied careful multiple testing correction within each omics dataset, we did not address multiple testing issues arising from analyzing multiple omics datasets and phenotypes. This may have led to type I error rates higher than the intended 5%. Moreover, we have sought the validation of individual features mostly by comparing to the literature or by comparing with other multi-omics studies on healthy human volunteers. Validation of these findings will require additional studies. Hence, the value of our current study is mainly in guiding more accurate future studies rather than in drawing conclusions on individual omics features.

In our studies, we mostly used computational methods (robust linear models, PCA, MOFA) that identify linear associations between omics profiles and phenotypes. There are alternative methods for finding latent dimensions across different omics layers. For example, more advanced techniques such as variational autoencoders are promising methods that capture non-linear relations across the different omics layers^75^. Potentially, such methods could reveal more complex interactions across the omics layers that have not been found here.

### Concluding remarks

The presented multi-omics study was undertaken within the context of the EATRIS-Plus project. EATRIS-Plus aimed to provide innovative scientific tools to the research community in the field of Personalized Medicine and increase the quality of the services provided by EATRIS (European Infrastructure for Translational Medicine). Best practices, guidelines and standard operating procedures arising from this and other studies were brought together in EATRIS’s multi-omics toolbox (MOTBX at motbx.eatris.eu), a public resource for multi-omics researchers. Moreover, we put significant efforts into making both our data and computational workflows FAIR (see data availability section), facilitating the use of this data for comparative analyses in future multi-omics studies in precision medicine, and studies developing or benchmarking multi-omics integration algorithms.

## Methods

### Clinical trial design

The project used a well-characterised Czech cohort of 1,109 healthy volunteers recruited during blood donations who were clinically examined and whose samples were bio-banked within the clinical trial “Assessment of Multi-omics Profiles in Health and Disease (ENIGMA)", NCT04427163. The inclusion criteria for participants in the study were: (a) age between 18-68, (b) absence of severe genetic diseases or a family history of such diseases, (c) absence of premature onset of civilization diseases such as hypertension, diabetes, autoimmune and cancer diseases, ongoing infectious or cancerous diseases, clinically manifest cardiovascular or pulmonary disorders, (d) not on long-term medication at the time of collection of biological material. An exclusion criterion was considered a failure to meet any of the inclusion criteria mentioned above. Participation in the study involved a visit for sample collection, including peripheral blood (BD Vacutainer EDTA and heparin blood collection tubes). All medical examinations are considered safe and minimally invasive.

### Sample collection

Samples have been collected at the University Hospital Olomouc and biobanked at the Institute of Molecular and Translational Medicine (IMTM, Palacký University Olomouc). Blood plasma was separated by centrifugation and stored at *−*80*^◦^*C until analysed. Blood leukocytes were isolated after osmotic lysis of red blood cells, lysed in TRIzol™ Reagent (ThermoFischer Scientific) and stored at *−*80*^◦^*C until the total RNA or DNA extraction and analyses. The average time from blood sampling to processing did not exceed one hour. All samples were distributed from the IMTM biobank to EATRIS-Plus partner sites on dry ice.

A representative subset (age and sex balanced) of 127 samples was selected (Table S1-2). The samples were selected according to the parametric or non-parametric distribution (including box-cox transformation strategies, 95% confidence interval). The subset cohort was sex balanced, with an age span ranging from 21 to 61 years. The IMTM distributed these samples to EATRIS-Plus centres for further omics measurements.

### Inclusion and Ethics statement

The multi-omics cohort used in EATRIS-Plus project was approved by the Ethics Committee of the University Hospital in Olomouc and the Faculty of Medicine of the Palacký University in Olomouc under the reference number 119/18 on 9 July 2018. The EATRIS-Plus project is supervised by EATRIS-Plus Ethics Committee. EATRIS-Plus Ethics Committee ensures that the EATRIS-Plus project’s ethical compliance is guaranteed, and that the project meets the highest ethical standards while maintaining compliance with national regulations. More information can be found in EC Annual Reports (https://eatris.eu/eatris-annual-reports/).

### Genomics - WGS

#### Sample preparation

WGS data was acquired at the Institute of Molecular and Translational Medicine, Palacky University. The TruSeq DNA PCR-Free LT Kit (Illumina) was used to prepare samples for whole genome sequencing. WGS libraries were prepared according to the manufacturer’s protocol. The input DNA amount for WGS library preparation with genomic DNA for TruSeq-PCR-free libraries was 1 µg. The same fragmentation conditions for all samples on Covaris instrument were used with a targeted size of 350 bp. The concentration of the TruSeq DNA PCR-Free libraries for WGS was measured by fluorometry using Qubit 2.0 fluorometer with HS dsDNA Assay kit (ThermoFisher Scientific, Q32854). The samples were equimolarly pooled based on the measured concentration and final quality check of sample pool was measured by qPCR with the KAPA Library Quantification Complete Kit (Universal) (Roche, KK4824); by fluorometry using Qubit 2.0 fluorometer with HS dsDNA Assay kit (ThermoFisher Scientific, Q32854); and by capillary electrophoresis using 2100 Bioanalyzer instrument (Agilent) with the High Sensitivity DNA Kit (Agilent, 5067-4626).

For the WGS library preparation, the sequencing was performed using Illumina platform NovaSeq 6000 at 2 x 161 bases read length using the S4 configuration (cat 20028312). Sequencing was performed following the manufacturer’s instructions.

#### Data processing

Raw sequencing data were processed to obtain sequence information. Quality control (QC) was performed using *FastQC (v0.11.9)*. The filtering of low-quality reads and adapter contamination trimming was performed using *fastp (v0.20.1)*. The remaining reads were aligned to the reference human genome hg38/GRCh38 primary assembly with the software package *BWA (v0.7.17-r1188)* using default parameters. This was followed by the task of marking duplicate reads and fixing mate information using *Picard tools (2.27.5)*^76^.

QC steps and the coverage were checked using the in-house software Genovesa https://www.bioxsys.eu/#/genovesa (developed by Bioxsys, s.r.o). Single-nucleotide variants (SNVs) and insertion/deletion variants (indels) were called using the VarScan v2.4.4^77^ (with parameters –min-coverage 20; –min-var-freq 0.1; —p-value 0.5, —min-avg-qual 15).

The variant calling process is based on a Fisher’s Exact test (applying algorithms Samtools, Varscan, Freebayes, and their combinations), a statistical test procedure that calculates an exact probability value for the relationship between two dichotomous variables, as found in a two-by-two crosstable. The program calculates the difference between read counts supporting reference and variant alleles with a p-value threshold 0.05. Only SNVs and indels passing the quality filter (a minimal quality of coverage *≥* 30, base quality *≥* 10, mapping quality *≥* 5) and with an alternative allele frequency *≥* 20% per sample, p-value (Fisher exact test) < 0.05 were included for further variant filtering.

### Genomics - aCGH

aCGH data was acquired at the Institute of Molecular and Translational Medicine, Palacky University. The method is based on the complementary binding of tested DNA to 2.6 million oligonucleotide markers immobilized on microarray cartridge. The obtained fluorescent intensities are statistically processed and sample genotype changes are obtained by comparing to the reference model. The array type CytoScan HD (Thermo Fisher Scientific) was used.

Total genomic DNA (130 ng) was fragmented, labelled with fluorescent dyes and hybridized onto the microarray according to the manufacturer instructions. Arrays were scanned using GeneChip Gene Scanner 3000 7G (Thermo Fisher Scientific), the fluorescent intensity for each oligonucleotide markers was measured and .DAT files (image file) were generated. Each marker has its own known position on the array. Using annotation library, the fluorescent intensity for each marker was calculated and .CEL files were subsequently generated. The scanning process and generation of .DAT and .CEL files were performed using Affymetrix GeneChip Command console version.

The generated .CEL files were annotated and analyzed using Chromosome analysis suite (CHAS) (Applied Biosystems by Thermo Fisher Scientific), version 3.3.0.139 (r10838) and genome version hg38. The generated .CYCHP files were viewed using CHAS. Table of Array Data Quality Control Metrics and a probeset table were created for all available 126 processed CELL files corresponding to the Czech Cohort samples based on the text exports from CHAS. For a subset of 118 samples that passed the QC thresholds, separate files with signal restored log2 ratio values of CN probes were created. These files served as input to the segmentation process.

The log2 ratio values were processed separately for each sample using the *rCGH R/Bioconductor*^78^ package with default parameter settings, circular binary segmentation algorithm (CBS)^79^ and Expectation-Maximization algorithm (EM) for genomic profile centralization^78^. Each resulted segmentation table was converted into by-gene table and finally combined into one data matrix (2 discordant samples were excluded). The missing value imputation by the Random Forest method implemented in the missForest R package^80^ was applied to the complete by-gene table consisted of 34,899 gene log2 ratio values for 116 samples. For multi-omics integration process only the subset of 3,990 genes with at least one sample log2 ratio value outside the interval for normal gene status <-0.5,0.5> was selected (Suppl Table 1).

### Epigenomics – Enzymatic methylation sequencing

EM-seq data was acquired at the SciLifeLab, Uppsala University. Sequencing libraries were generated from 100 ng of DNA using the NEBNext Enzymatic Methyl-seq sample preparation kit (cat E7120S/E7120L, NEB), incorporating unique dual indexes. Library preparation strictly followed the manufacturer’s instructions (manualE7120), inclusive of the addition of control DNAs (pUC19 and Lambda). Quality control assessment was conducted using the Fragment Analyzer equipped with the DNF-910 dsDNA kit (Agilent) to ensure library integrity. Libraries were quantified by qPCR, utilizing the Library Quantification Kit for Illumina (KAPA Biosystems) on a CFX384 Touch instrument (Bio-Rad).

Sequencing was performed on an Illumina NovaSeq6000 system, with a paired-end 151 bp read configuration (Illumina, San Diego, CA, USA). All libraries were sequenced to a minimum of 300 M paired-end reads per sample with read length 2x151 bp. Sequencing data was demultiplexed and converted from base calls into FASTQ files using Illumina bcl2fastq2 Conversion Software v2.20.0.422. Quality control after sequencing was performed using the in-house quality control tools CheckQC www.github.com/Molmed/checkQC and Seqreports www.github.com/Molmed/seqreports as well as FastQ Screen from Babraham Bioinformatics (www.bioinformatics.babraham.ac.uk/projects/fastq_screen/). Primary methylation sequencing data analysis was performed with nf-core analysis pipeline methylseq (version 1.6.1), the reference genome used was Ensembl GRCh38, release 105. Pipeline output includes alignment to the reference genome, methylation calls and quality metrics. The controls pUC19 and lambda were aligned using Bismark version 0.23.0 (Babraham Bioinformatics) and the methylation rate was compared against threshold provided by vendor. Quality control after primary analysis consisted of PCA (Fig S1) on the sites (overlapping (3000) a set of array probes selected for sex classification^81^.

We used the *methrix (v1.10.0)* R/Bioconductor package for data processing using BSgenome. Hsapiens.NCBI.GRCh38 as reference genome. Samples that failed on prior QC measures for sequencing for low conversion rate (n=2) or low alignment rate (n=3) were excluded from further analysis. Information from the different strands on the same CpG site was merged. The resulting object was further filtered by removing all CpG sites covered by < 5 reads in >50% of the samples (n= 1,525,790). CpG-level DNA methylation measurements were indicated as missing values (NA) if the coverage was <5 reads for a given CpG site and sample. The CpG sites not covered by any samples were removed from the dataset (n = 12,775). The sites overlapping with common SNPs as defined by a minor allele frequency > 0.01 in the 1kGenomes based on *MafDb.1Kgenomes.phase3.hs37d5* data package were removed using the *remove_snps* function as implemented in the methrix R package (n = 3,452,226). Cell type composition was assessed using an in-house implementation of the reference-based Houseman algorithm^82^ using DNA methylation from the recently published DNA methylation atlas^83^ (Fig S2).

To avoid the large number of CpG sites dominating the multi-omics analyses, we selected the 100,000 most variable CpG sites. To adjust the dataset for cell type composition, we first fitted a linear model between the cell type composition and the beta values for each CpG site. If the given site was associated with the cell type composition with a p < 0.01, we exchanged the beta values with the residuals of the model. If not, we used the centered beta value.

### Transcriptomics – mRNAseq

mRNAseq data was acquired at the Institute for Molecular Medicine Finland. Library preparation from 800 ng of total RNA was performed according to the Illumina stranded mRNA prep Reference Guide (Illumina, San Diego, CA, USA). Library quality check was performed using LabChip GX Touch HT High Sensitivity assay (PerkinElmer, USA). Normalized and pooled libraries were quantified using KAPA Library Quantification Kit (KAPA Biosystems, Wilmington, MA, USA) and sequenced on the Illumina NovaSeq6000 system to a minimum of 50 M paired-end reads for each sample (Illumina, San Diego, CA, USA). Read length for the paired-end run was 2x151 bp. Sequencing data is demultiplexed and converted from base call files into FASTQ files with Illumina *bcl2fastq2 Conversion Software (v.2.20)*. Sequencing was performed in multiple batches, each with a reference sample Brain RNA for normalization purposes (Human brain total RNA (Thermo Fisher AM7962). Primary RNA-sequencing data analysis is performed with in-house Nextflow-based FIMM-RNAseq data analysis pipeline version 2.0.1. Pipeline outputs include alignments to reference genome and various quality metrices. The gene counts that were produced by the pipeline underwent a variance stabilization transformation from R/Bionconductor’s package *DESeq2 (v 1.36.0)*, which also included correction for size and normalization factors and provided log-transformed counts.

### Transcriptomics – MicroRNAseq

miRNAseq data was acquired at the Institute for Molecular Medicine Finland. A clean up of the samples was performed using the miRNEasy micro kit from QIAGEN (cat. no. 74004). Total RNA was eluted using 15 *µ*L RNase-free water. Library preparation from 600 ng of total RNA was performed in batches of 6-24 samples according to Lexogen Small RNA-Seq library prep Reference Guide (Lexogen GmbH, Vienna, Austria). Library quality check was performed using Agilent Bioanalyzer High Sensitivity DNA assay (Agilent Technologies, Waldbronn, Germany) and libraries were pooled based on the concentrations acquired from the assay. The pooled libraries were subjected to size selection targeting fragments that were 130-180 bp in size using BluePippin (Sage Science, Beverly, Massachusetts, USA). The size-selected library pools were quantified using KAPA Library Quantification Kit (KAPA Biosystems, Wilmington, MA, USA) and sequenced on the Illumina NovaSeq6000 system for 100 cycles using the S1 flow cell with a lane divider to a minimum read depth of 10 M reads for each sample (Illumina, San Diego, CA, USA). Read length for the single-end run was 101 bp.

Sequencing data is demultiplexed and converted from base call files into FASTQ files with Illumina *bcl2fastq2 Conversion Software (v.2.20)*. Primary miRNA-sequencing data analysis is performed with Nextflow-based FIMM-smRNAseq data analysis pipeline. General output includes a sample-specific read count file describing the number of reads mapping to each miRNA and an alignment visualization file.

### Transcriptomics – microRNA qRT-PCR

The microaRNA qRT-PCR data was acquired at the TTranslational Genomics& Bioinformatics Core Facility of Ramon y Cajal Health Research Institute. Total RNA enriched in miRNAs was isolated using miRNeasy Serum/plasma advanced kit (217204, QIAGEN) starting from 250 *µ*L of plasma and eluted in a final volume of 25 *µ*L, following the manufactures instructions. Before RNA isolation, 1 *µ*L of synthetic RNA Spike-in was added to each sample (miRCURY RNA Spike-in kit). A spike in amplification serves as a technical control of RNA extraction efficiency and homogeneity. We used the Universal RT miRNA PCR System (Qiagen) for cDNA synthesis. Briefly, 16 *µ*L of RNA was used as a template for RT in a final volume of 80 *µ*L. An external RNA (cel-miR-39) was added and further amplified to cDNA synthesis efficiency control. Sample hemolysis was checked by miR-23a-3p and miR-451a amplification. Samples in which the subtraction of the expression of these miRNAs was higher than 7.5 were discarded due to elevated levels of hemolysis. 179 miRNA expression was evaluated by qRT-PCR using microRNA Array Profiling (miRCURY LNA miRNA Focus Panels; YAHS-106YG, Qiagen). All reactions were carried out in triplicate using a Light Cycler 480 instrument (Roche, Basel, Switzerland), and Cq values were calculated using a 2nd derivative method (Light Cycler 480 Software 1.5, Roche). Raw Cq values were transformed to Z-scores and normalized among plates, replicates and samples. Replicates with missing values higher than 18 miRNAs ( 10% of the miRNAs) were removed from the analysis. Internal controls (mean of UniSP2 *>* 25, mean of cel-miR39 *>* 30 and N° of missing values > 40%) and hemolysis were used as quality controls to remove samples from the study.

### Proteomics (LC-MS)

LC-MS proteomics data was acquired at the Institute of Molecular and Translational Medicine, Palacky University. Plasma peptide mixtures were prepared from plasma samples by the FASP protocol adapted from^84^. The protein concentration was determined by BCA assay, and 100 *µg* of total proteins were processed. All processed plasma samples were diluted into a 0.1 *µg/µl* concentration. Dilution was based on the results from the peptide concentration assay (see MOTBX protocol "Sample preparation for plasma analysis"^85^). During dilution, PROCAL peptides (JPT, cat. n. RTK-1-100pmol) are added to obtain a final concentration of 10 fmol*/µl*. Each sample was analyzed in two technical replicates. The 1 µg of peptides and 100 fmol of PROCAL peptides in injection volume 10 µl for 1 technical replicate were used. Peptides were separated by liquid chromatography (UltiMate 3000 RS series, Thermo Scientific™) and detected by mass spectrometry (Orbitrap Fusion™ Tribrid™, Thermo Scientific™). Measurements of all 127 Czech Cohort samples consisted of 4 batches. Each batch started with an SST Test (5 injections of BSA standard); samples were released for analysis based on the results of the SST. The 10 injections of plasma peptide samples (5 samples in duplicate) and 1 BSA standard were running repeatedly until all plasma samples in the batch were measured. All detailed parameters of liquid chromatography and mass spectrometry settings are described in MOTBX protocol "LC-MS (HCD OT) Plasma Analysis"^86^. Raw data of each technical replicate were searched in Proteome Discoverer (PD) software, (v. 2.5) (Thermo Scientific) in two subsequent steps -the processing and consensus steps. The processing workflow included a search against the complete human UniProtKB database, including Swiss-Prot reviewed and TrEMBL computationally analyzed but unreviewed proteins (https://www.uniprot.org/uniprot/, downloaded in July 2020). In the consensus step, the normalization of abundances was done on specific protein amounts of PROCAL peptides. Detailed settings of both steps have been described in the SOP_Search_PD2.5_Plasma analysis. Results with identified and quantified proteins were exported from PD 2.5 in tab-separated format and processed with software R.

The raw abundances were log2 transformed to make data distribution close to normal. Replicates abundances were aggregated by mean and three filters after exclusion of QC feature were applied: 979 proteins completely missing in at least one batch were excluded, and 29 low FDR confidence (q-value *≥* 0.05) proteins were removed. Only features with no more than 30% of missing values were kept in the dataset. After filtering the dataset consisted of 1393 proteins (features) abundances in rows and 127 samples in columns with 7.9% of missing values in total. A local similarity random forest method implemented in missForest R package^80^ was chosen for imputation and a dataset with complete observations was created.

### Proteomics (IPP)

IPP proteomics was acquired at Institute of Applied Biotechnologies a.s., Science and Technology Park, Palacký University Olomouc. Analysis was performed using the Illumina Protein Prep 6K kit, plasma, 96 reaction (Illumina Inc., San Diego, CA, USA, catalog #20137827) , Illumina DNA/RNA UD Indexes, Tagmentation 96 Indexes, 96 Samples Set A, Set B (Illumina Inc., San Diego, CA, USA, Set A catalog #20091654, Set B catalog #20091656) according to the manufacturer’s standard protocol. Human plasma samples were processed together with internal calibrators and control samples (plate controls and SOMAmer reagent controls) to minimize technical variability. The entire workflow is fully automated on the Illumina Protein Prep Automation System built on the Tecan Fluent 780 platform (Illumina Inc., San Diego, CA, USA, platform catalog #20116818), allowing robotic execution with minimal manual handling (Illumina, 2025). An input volume of 55 *µl* of plasma was used per sample (Illumina Support). Proteins were incubated with SOMAmer reagents, enabling selective binding to target proteins using Quad Microplate Shaker-Incubator Thermoshaker (VWR, catalog #97025-554). After incubation, a series of steps followed, including streptavidin bead capture, UV photolysis of bound complexes, transfer, washing, and elution (Illumina, 2025). Eluted SOMAmer–protein complexes were converted into barcoded sequencing libraries through over-night hybridization PCRmax Alpha Cycler 4 Thermal Cycler, Quad 96-Well Capacity (Cole-Parmer, catalog # EW-93945-12) and adapter ligation, followed by PCR amplification C1000 Touch Thermal Cycler with 96–Deep Well Reaction Module (Bio-Rad, catalog #1851197) (Illumina, 2025). Libraries were sequenced on Illumina NovaSeq X Plus using NovaSeq X Series 10B Reagent Kit (100 cycles) (Illumina Inc., San Diego, CA, USA, catalog #20085596) and data were processed using the DRAGEN Protein Quantification pipeline (version 1.8.33), which automatically performs BCL → ADAT conversion, normalization, quantification, and QC metric calculation (Illumina, 2025). Data normalization utilized internal controls and correction for intra – and inter-plate variability. Resulting ADAT files containing quantitative values for 6000 proteins were further subjected to statistical analysis, including outlier removal, log2 transformation, and multiple testing correction using the Benjamini–Hochberg method. The detailed Illumina Protein Prep 6K protocol, including reagent specifications, incubation steps, times, and volumes, is available from the manufacturer (Illumina Protein Prep protocol).

### Targeted Metabolomics

#### Data acquisition

Quantitative targeted metabolomics data were acquired at Radboud University Medical Center (Radboudumc) and Maastricht UMC+ (MUMC+). Plasma acylcarnitine analysis (ACRN, 44 metabolites) was performed at Radboudumc according to a protocol previously described^87^. Plasma amino acid analysis (AA, 53 metabolites) and very long chain fatty acid analysis (VLCFA, 5 metabolites) were performed at MUMC+ according a previously described^88^ and an in-house protocol, respectively. Metabolite levels were reported as µM concentrations.

The 127 plasma samples from the Czech cohort were measured in multiple batches: four batches for ACRN, two batches for AA, and six batches for VLCFA analysis. For ACRN analysis, a mixture of compounds with known concentrations (quality control / QC sample) was measured along with each batch. For AA analysis, two different mixtures of compounds with known concentrations were measured along with each batch. For ACRN analysis, the limit of detection (LOD) is 0.01 µM. For AA analysis, the limits of quantification (LOQ) are below 1 µM, except for ethanolamine (hmdb:HMDB0000149) where LOQ is 2.5 µM, and for formiminoglutamic acid (hmdb:HMDB0000854), 3-methylhistidine (hmdb:HMDB0000479), 1-piperideine-6-carboxylic acid (chebi:49015), and allysine (chebi:17027) for which LOQ were not determined. Metabolites that could not be detected (true value < LOD) were reported with a concentration of 0 µM.

#### Data pre-processing and preparation

Metabolites missing in *≥* 30% of the samples were removed. Since metabolite concentrations follow a log-normal distribution more closely than a normal distribution, concentrations were log-transformed using the natural logarithm after adding a constant value to each value (log(x+1)). For downstream analyses, the metabolite features were mean-centered to zero and scaled to unit-variance. Missing values were imputed using Low abundance resampling (Table S4, Suppl Fig S39-41).

### Untargeted Lipidomics

#### Sample preparation

Before the analysis, 50 *µ*L of plasma (thawed on ice) was mixed with 150 *µ*L of ice-cold LCMS isopropanol containing Avanti SPLASH internal standard mixture, FA 20:4(d8), Cer d18:1(d7)/15:0 and vortexed for 20 s. Samples were centrifuged (22,400g, 20 mins at 4*^◦^*C), and 150 *µ*L of the supernatant was loaded into a low recovery volume HPLC vial with insert. Blank samples were prepared using the same procedure instead of gradient fraction LCMS water. Pooled quality control samples (QC) were prepared by mixing 10 µL of plasma from every sample and thoroughly vortexed (5 min). Each QC (n=18) sample was prepared individually in the same way as the biological samples. QC samples were injected as every 6th sample through the analytical batch. The standard reference material samples (SRM NIST 1950) were prepared the same way as other biological samples and injected at the beginning and end of the analytical batch. Samples were randomised for the first time before the sample preparation. Second, separate randomization was done for the data acquisition.

#### LC-MS Analysis

Lipidomics data was acquired at the Institute of Molecular and Translational Medicine, Palacky University. Separation of non-polar compounds was done using reversed-phase chromatography separation mechanisms on an Accucore C30 column (150 mm ×2.1 mm, 2.6 *µ*m; Thermo Fisher Scientific, MA, USA). The constitution of mobile phase A was 20 mM ammonium formate in 60:40 (v:v) acetonitrile:water and mobile phase B consisted of 20 mM ammonium formate in 85.5:9.5:5 isopropanol:acetonitrile:water (v:v:v).

The gradient was as follows: t= 0.0, 20% B; t= 2.5, 20% B, t= 2.6, 55% B; t= 12, 60% B; t= 12.1, 80% B; t= 19.0, 90% B; t= 21.0, 100% B; t= 23.0, 100% B; t= 23.1, 20% B; t= 30.0, 20% B. All changes were linear (curve = 5), and the flow rate was 0.400 mL/min. The column temperature was 55*^◦^*C, and the injection volume was 2 *µ*L. Data were acquired separately in positive and negative ionization modes (150–2000 m/z) with a resolution of 60,000. Ion source parameters: sheath gas = 48 arbitrary units, aux. Gas = 15 arbitrary units, sweep gas = 0 arbitrary units, spray voltage = 3.5 kV (positive ion)/2.5 kV (negative ion), capillary temp. = 350 °C, aux. gas heater temp. = 400 °C. The full scan data were acquired separately in files without any MS/MS scans.

The detailed settings were as follows: mass resolution =17,500 (FWHM at m/z 200); isolation width =3.0 m/z; stepped normalised collision energies (stepped NCE) = 20, 40, 100 for positive ionization mode and (stepped NCE) = 40, 60, 130 for negative ionization mode. Spectra were acquired with different precursor m/z ranges in different segments of the chromatogram: t =0–6 min, 150–1100 m/z; t =6–12 min, 550–1250 m/z; t =12–17 min, 550–1500 m/z; t =17–30 min, 550–1100 m/z in case of 30 min elution gradient and for the 50 min elution gradient the segments were as follows: t =0–10 min, 150–1100 m/z; t =10–20 min, 550–1000 m/z; t =20–25 min, 550–1100 m/z; t =25–30 min, 550–1500 m/z; t =30–40 min, 550–1050 m/z; t=40–50 min, 550–1500 m/z. Thermo Tune Plus (2.7.0.1112 SP2) software controlled the instrument. All data were acquired in profile mode^89^.

### Lipidomics set 1 data processing

Data were processed and evaluated using Compound Discoverer 3.3 SP1 software (Thermo Fisher Scientific, MA, USA). The workflow was set as follows: Chromatograms were aligned using the self-learning function “ChromAlign” using a QC sample as a reference. Peak detection & Feature grouping was done directly from Thermo Fisher Scientific profile data (*.raw). The missing values were filled using the “Fill Gaps node – Real Peak detection” option. Signal/Batch correction was performed by SERRF QC Correction (Systematical Error Removal using Random Forest, n=200)^90^. The data matrix was filtered using criteria as follows: Compounds with > 50% missing in QC samples and with relative standard deviation (RSD) > 30% were removed; Compounds with corrected areas with RSD > 25% were removed. Blank substruction was done with the ratio of intensity sample/blank = 20/1. Finally, the data matrix was normalised using Median Absolute Deviation (MAD). For the purposes of statistical evaluation, the data matrix was log10-transformed, and univariate scaling was performed. The complexity of metabolite identification is depicted in Suppl Fig S52.

The identification of the compounds was performed on various levels using mzVault and LipidBlast (V68) spectral libraries as well as ChemSpider and LipidMAPS online databases.

### Lipidomics set 2 data processing

Data were processed in R 4.3.1 (released June 2023) using XCMS 3.18.0. Peaks were detected with the centWave algorithm (ppm = 14, snthresh = 10, peakwidth = c(4, 40), mzdiff = 0.0045, prefilter = c(3, 100)). Retention-time alignment (bw = 0.25) and feature grouping (mzwid = 0.00835) were performed using default XCMS workflows^91^.

Data pre-processing was performed in R 4.3.1 with Bioconductor 3.18, using the structToolbox (v1.14.0)^92,93^ and pmp (v1.14.1)^94^ packages. Signal drift and batch effects were corrected using the QC-Robust Spline Correction (QC-RSC) implementation (sb_corr, spar_lim = c(0.6, 0.8)), with run order (run_order), batch (Batch) and sample type (SampleType, QC label = “QC”) as inputs. Features that failed correction were removed by excluding variables with more than three missing values per batch. Blank-related artifacts were filtered using blank_filter (fold-change threshold = 20, QC vs. blanks), followed by exclusion of samples annotated as Blank. The first eight conditioning QC injections were excluded from downstream analyses. Multi-batch feature filtering included a Kruskal–Wallis test across batches (*α* = 1 *×* 10*^−^*^4^, FDR-adjusted) and a Wilcoxon signed-rank test (*α* = 1 *×* 10*^−^*^14^, FDR-adjusted) to remove unstable and non-representative features, respectively, and an RSD filter to remove features with QC RSD > 20%. Missing-value filters were then applied to retain only features present in at least 80% of QC samples and 50% of all samples. Finally, data were normalized using Probabilistic Quotient Normalization (PQN) (QC-based), missing values were imputed by k-nearest neighbors (k = 5), and a generalized log (glog) transform was applied before statistical analysis.

### Batch effect adjustment

In some omics datasets, batch effects originating from the running of samples on different days needed correction (Suppl Figures S42-50, Tables S3-5). Batch effect correction for lipidomics was performed based on pooled quality control (QC) samples applying the SERRF QC Correction algorithm^90^. For all other omics datasets where an internal standard was not available, batch correction was done in a consistent manner using Combat^95^, implemented with R/Bioconductor’s *sva (v3.44.0)*. The parametric method was applied using sex as covariate to retain information related to sex in an unbalanced batch design and without covariate. For the mRNA-seq data, Combat was applied on the gene counts table with the *Combat_seq* function, which is designed specifically for bulk RNA-seq count data. Combat optimally reduced associations with technical confounders (Suppl Figures Suppl Figures S42-50.

### Cell type correction

The estimated cell type proportions derived from the EM-seq data (see Methods Epigenomics), were used to correct for the influence of cell types on the transcriptomics and epigenomics data. This correction process involved employing a linear model where the estimated cell type numbers were used as covariates, after which the residuals of this model were retrieved for downstream analysis. The estimated cell type compositions can be found in Suppl Fig S6.

### Reference values

For reporting of reference values, mean, median, standard deviation, minimum, and maximum –omics levels were calculated across all individuals in the cohort. Additionally, the CV was calculated by dividing the standard deviation by the mean for every feature and multiplying by 100 to derive percentages. The reference values were reported for male and female individuals separately and are available in Suppl Table 3.

### Single-omics data analyses

Single –omics analyses were performed using R version 4.2.1. The robust linear models were implemented using R package *MASS (version 7.3-60)*^96^. For each feature, linear models were run with no covariates, sex as covariate and for only females and males. PCA was performed using the built-in R function *prcomp*. Correlations of the principal component scores with the continuous phenotypic variables were tested using Pearson pairwise correlation. To ensure comparability across omics layers, all p-values were corrected using the Benjamini-Hochberg false discovery rate (FDR), despite differences in data dimensionality.

### Multi-omics data analyses

Multi–omics analyses were also performed using R version 4.2.1. Multi-Omics Factor Analysis (MOFA)^11^ was implemented in R using a combination of the R/Bioconductor library MOFA2 (version 1.6.0) and the conda library *mofapy2 (v 0.6.7)*, which was called upon by the reticulate package (version 1.3.1)^97^. Correlations of the individual’s factor scores with the continuous phenotypic variables were tested using Spearman pairwise correlation, where all p-values were corrected using Benjamini-Hochberg’s false discovery rate (FDR). The factor loadings of the mRNA-seq, proteomics, metabolomics and EM-seq data were used for gene set enrichment analysis. The R library *multiGSEA (v1.11.2)*^98^ was used to find significantly enriched pathways. The multiGSEA function was customized to enable parameter tuning. Standard deviation was used as gene set enrichment analysis (GSEA) score type and the boundary for calculating p-values was set to 1e-100. For the EM-seq data, gene set enrichment was carried out using the *gseGO* function of the R library *clusterProfiler (v4.2.2)*^99^, focusing on the Gene Ontology (GO). CpG sites were mapped to nearest genes using the R/Bioconductor library *biomaRt (v2.50.3)*^100^. As many CpG sites were mapped to the same gene, we selected the CpG site with the highest loading for every gene.

Independent Component Analysis was implemented in R/Bioconductor package *consICA (v.2.0.0)*^20^ in order to find 20 independent components per –omics dataset. Linear models that were used to predict phenotypic outcomes from the independent component, were created with standard R *lm* tool. Overrepresentation analysis was performed with R library *topGO (v 2.48.0)* within *consICA* package, independently for positively and negatively contributing genes from each component. Network of the correlated components was built using the threshold of *R*^2^ *>* 0.1, calculated on component weights over all common patients (corresponding p-value <0.001).

Similarity Network Fusion was implemented in Python version 3.9.5, using the library *SNFpy (v0.2.2)*^21^. A fused network was constructed for every combination of –omics data, with a minimum of 2 and a maximum of 12 omics datasets. The numbers of sample clusters were selected by a built-in function of the library, which finds the optimal number of clusters based on the eigengap method. The goodness of clustering for every combination was quantified with the silhouette score, which was determined with the *SNFpy* library as well. We assessed whether phenotypic differences could be found among the different SNF clusters, using Mann-Whitney U tests (two clusters) or Kruskal-Wallis tests (more clusters).

PLS2 was applied using the R/Bioconductor library *mixOmics (v6.20.0)*^22^ to find pairwise correlations among all –omics data. The sparse Partial Least Squares (sPLS) method was implemented using the canonical analysis algorithm, where 2 latent dimensions (PLS components) were calculated for each model. A maximum of 1,000 features per –omics l was included for each model, which the model selects using LASSO penalization. Pearson correlations were tested across the sample scores on the first PLS component.

For the SNP-CpG analysis, we used the set of methylation sites that were removed for the other analyses (see Methiods EM-seq). CpGs that overlapped with common SNPs (meCpG-SNPs) and that exhibited no more than 20% missing values across all samples were kept. Consequently, the WGS data was filtered for overlap with this specific set of CpG sites (CpG-SNPs). The resulting set of CpG-SNPs was filtered for minor allele frequency (MAF) 0.2 < MAF < 0.8. These were then filtered using the Hardy-Weinberg equilibrium (p-value > 0.05). Next, all neighbouring CpGs, within a range of 1000 bps up- and downstream of the SNp-CpGs, were selected to evaluate genotype-dependent differences in methylation spread across SNP–CpG groups.. Neighbouring sites that were found within this range of two SNP-CpGs were removed. In additon, we assessed differential gene expression across SNP–CpG genotypes using a linear model with genotype as the predictor and RNA transcript abundance as the response variable, assuming an additive allelic effect. Multiple testing correction was performed using the Benjamini–Hochberg procedure with a false discovery rate threshold of 0.05.

### Epigenetic clocks

Epigenetic clocks were calculated using the *methylclock (v1.6.0* Bioconductor package^101^. Since the methylation clocks are based on Illumina arrays, we filtered our dataset and identified the CpG sites for each site on the Illumina array. Performance of the clocks were assessed by correlating (Pearson) estimated age with reported age. Moreover, we calculated the age acceleration, the difference between biological and chronological age, and applied robust linear models as described above.

### Data availability

Both clinical and multi-omics data were collected and are stored in ClinData Portal https://clindata.imtm.cz/. ClinData is a software solution for collecting of GDPR compliant clinical and laboratory data on patients included in research projects and clinical studies. ClinData is based on client/server architecture, the application runs on a server, a user is connected through a web browser. All communication between the client and the server is secured through SSL encryption, an industry standard. The Java 8 programming language is used for the server, which allows sustainability and long-term development. The client interface uses open source technologies – HTML, Java Script, jQuery, Angular, Bootstrap.

Moreover, the processed multi-omics data, has been publicly available through Zenodo (doi.org/10.5281/zenodo.17514796).

However, genomics data has been excluded to preserve participant privacy.

### Code availability

All scripts that were used to perform both the single- and multi-omics analyses, are made available through a Nextflow^102^ pipeline: https://github.com/EATRIS/Multi_Omics_Workflow.git. The pipeline can be re-used for other datasets, as it has been developed following the FAIR4RS principles^103^.

## Acknowledgements

This work was supported by European Union’s Horizon 2020 under grant number 871096 (EATRIS-Plus) and the Nederlandse Organisatie voor Wetenschappelijk Onderzoek (NWO) under grant number 184.034.019 (X-omics). Part of this work was carried out with the support of ELIXIR CZ Research Infrastructure (ID LM2023055, MEYS CR).

## Author contributions statement

**Conceptualization:** C.V., A.N., L.N., J.S., E.O., T.A., G.S., A.S., J.N., A.G., M.H., P.A.H.; **Data curation:** C.V., A.N., L.N., J.V., J.S., S.E., B.G., V.F.L., M.B., Z.R., P.D., P.P., B.S., J.Sr., V.M., P.S., M.G., J.N., M.H., P.A.H.; **Formal analysis:** C.V., A.N., L.N., P.V.N., R.T., J.V., J.S., S.E., B.G., B.Y., V.F.L., J.G., P.K., H.W., U.E., A.Sp., D.B., V.M., P.S., M.G., J.N.; **Funding acquisition:** A.G., M.H., P.A.H.; **Investigation:** C.V., A.N., L.N., J.S., S.E., E.C.M., P.M., M.P., V.F.L., J.G., P.K., H.W., U.E., M.B., Z.R., P.D., P.P., B.S., J.Sr., J.N.; **Methodology:** C.V., A.N., P.V.N., R.T., M.P., A.M., M.B., Z.R., P.D., J.Sr., A.G., M.H., P.A.H.; **Project administration:** A.N., L.N., E.C.M., P.M., M.P., E.O., T.A., G.S., A.S., M.L.G.B., J.N., A.G., M.H., P.A.H.; **Resources:** M.P., J.G., H.W., U.E., M.B., Z.R., P.D., P.P., B.S., J.Sr., A.G.; **Software:** C.V., A.N., L.N., P.V.N., R.T., J.V., V.F.L.; **Supervision:** A.N., P.M., M.P., E.O., T.A., G.S., A.G., M.H., P.A.H.; **Validation:** P.M., M.B., Z.R., P.D., J.Sr.; **Visualization:** C.V., A.N., P.V.N., R.T., E.C.M., V.F.L., A.M., A.Sp., D.B., M.H., P.A.H.; **Writing – original draft:** C.V., A.N., L.N., P.V.N., R.T., J.V., J.S., M.P., V.F.L., A.M., M.L.G.B., J.N., A.G., M.H., P.A.H.; **Writing – review & editing:** C.V., A.N., L.N., P.V.N., R.T., J.V., J.S., E.C.M., V.F.L., A.M., M.H., P.A.H.

