## Supplementary Figures for "Comprehensive multi-omics profiling of a healthy human cohort"

#### Multi-omics cohort

| Phenotype trait | Mean | Median | Min | Max | SD |
|---|---|---|---|---|---|
| age | 43.91 | 44 | 23 | 62 | 8.63 |
| height | 174.14 | 174 | 156 | 197 | 8.05 |
| weight | 81.36 | 81 | 56 | 110 | 13.04 |
| body-mass.index | 26.78 | 27 | 21 | 37 | 3.35 |
| leukocytes | 6.37 | 6.2 | 3.9 | 10 | 1.36 |
| neutrophils | 0.59 | 0.6 | 0.41 | 0.78 | 0.08 |
| lymphocytes | 0.32 | 0.31 | 0.14 | 0.49 | 0.07 |
| haematocrit | 0.43 | 0.43 | 0.37 | 0.5 | 0.03 |
| erythroctyes | 4.79 | 4.81 | 3.94 | 5.57 | 0.36 |
| hemoglobine | 147.53 | 147 | 125 | 172 | 10.68 |
| mcv | 89.89 | 89.9 | 82.3 | 99.3 | 3.22 |
| mch | 30.84 | 30.9 | 26.7 | 33.5 | 1.2 |
| mxd | 0.09 | 0.09 | 0.04 | 0.18 | 0.03 |
| platelets | 245.96 | 240 | 154 | 410 | 52.54 |
| mchc | 343.22 | 341.5 | 323 | 370 | 9.97 |

**Table S1 Cohort summary statistics of numerical phenotypic traits.**

| **Phenotypic trait: sex** | | |
|---|---|---|
| Value | Count | Percentage |
| FEMALE | 64 | 50.4 |
| MALE | 63 | 49.6 |
| **Phenotype: smoking status** | | |
| Value | Count | Percentage |
| Current Smoker | 26 | 20.5 |
| Former Smoker | 36 | 28.3 |
| Never Smoker | 64 | 50.4 |
| Unknown If Ever Smoked | 1 | 0.8 |
| **Phenotype: blood.type.abo** | | |
| Value | Count | Percentage |
| Blood Group A | 55 | 43.3 |
| Blood Group AB | 15 | 11.8 |
| Blood Group B | 29 | 22.8 |
| Blood Group O | 28 | 22 |
| **Phenotype: blood.type.rh** | | |
| Value | Count | Percentage |
| Rh Negative Blood Group | 22 | 17.3 |
| Rh Positive Blood Group | 105 | 82.7 |

**Table S2. Cohort summary statistics of categorical phenotypic traits.**

| Omics layer | Omics platform (assay/technology) | Dataset |
|---|---|---|
| Metabolomics | Acylcarnitines (electrospray tandem MS) | 33 acylcarnitines |
| Metabolomics | Amino acids (UPLC–MS/MS) | 46 amino acids |
| Metabolomics | Very long chain fatty acids (UPLC–MS/MS) | 5 VLCFAs |
| Lipidomics | LC–MS lipidomics | 164 lipids (Set 1 neg mode) |
| Lipidomics | LC–MS lipidomics | 196 lipids (Set 1 pos mode) |
| Lipidomcs | LC–MS lipidomics | 257 lipids (Set 2 pos mode) |
| Proteomics | LC–MS proteomics | 1,393 proteins |
| Proteomics | Illumina Protein Prep proteomics | 6,831 proteins |
| Transcriptomics | mRNA-sequencing | 19,747 mRNAs |
| Transcriptomics | miRNA-sequencing | 2,130 miRNAs |
| Transcriptomics | miRNA qRT-PCR panel | 179 miRNAs |
| Epigenomics | Enzymatic methylation sequencing | 100,000 CpG sites |
| Genomics | Array Comparative Genomic Hybridization | 2,549 genomic regions |
| Genomics | Whole-genome sequencing | N/A |

**Table S3. Overview of the multi-omics data set.**

### Genomic aberrations

#### Klinefelter Syndrome identification

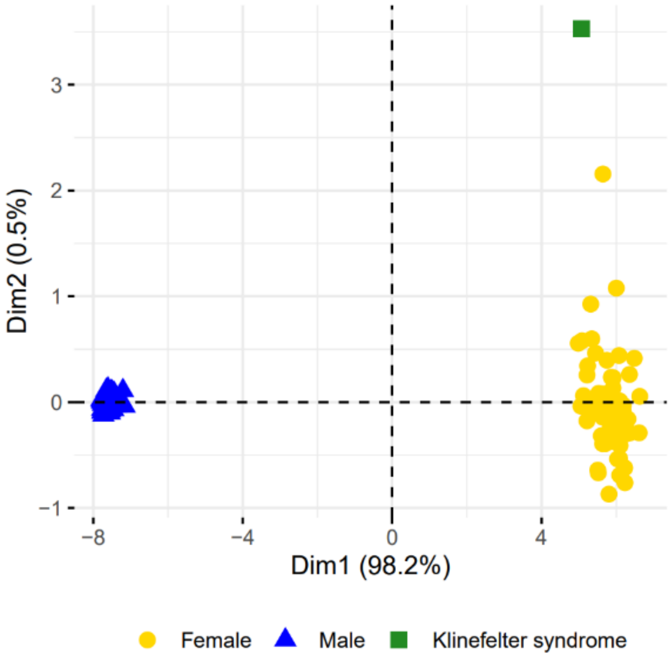

**Figure S1. Principal component analysis (PCA) of the EM-seq data. The sample outlier in this score plot was identified as an individual with Klinefelter syndrome.**

### Effect of gene gain/loss on mRNA levels

For genes with significant loss and/or gain in the healthy subjects, we compared the mRNA levels. We observed six genes with significant transcript levels among groups with gene loss, gene gain or no gene aberrations.

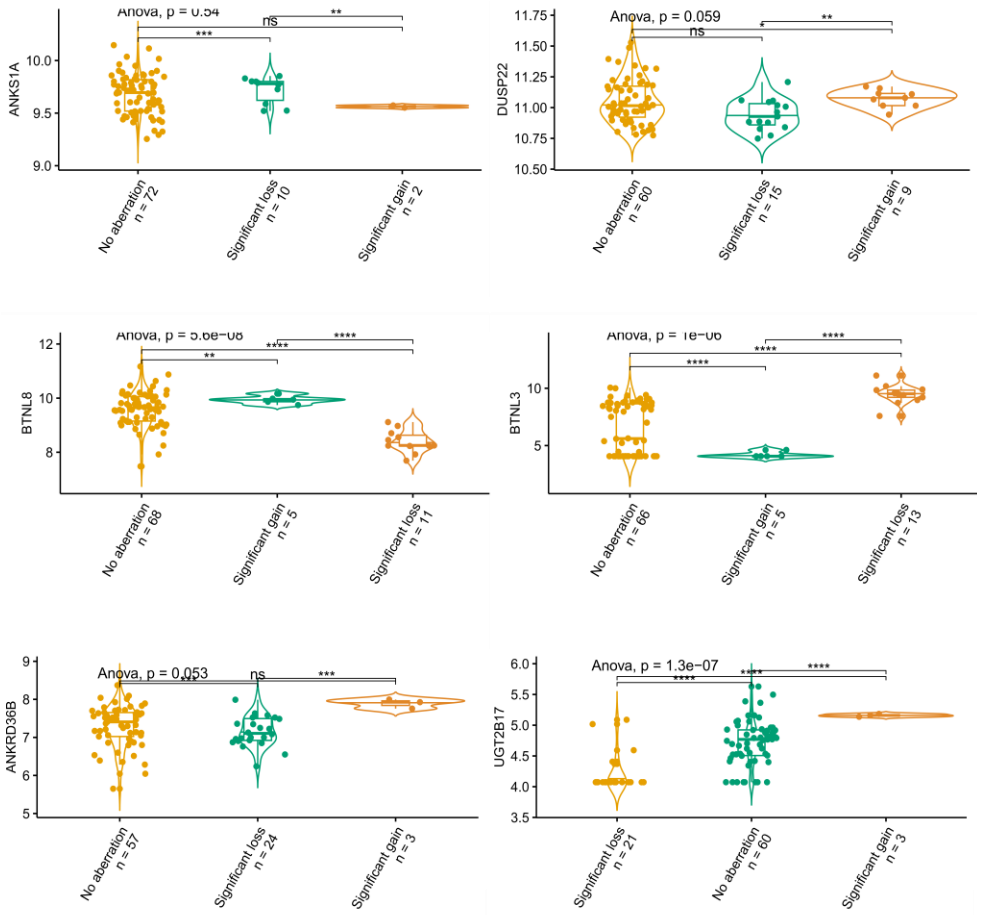

**Figure S2: Levels of mRNAs *ANKS1A, ANKRD36B, UGT2B17, DUSP22, BTNL8* and *BTNL3* were compared among groups with gene loss, gene gain or no gene aberrations.**

#### SNP-CpG analysis

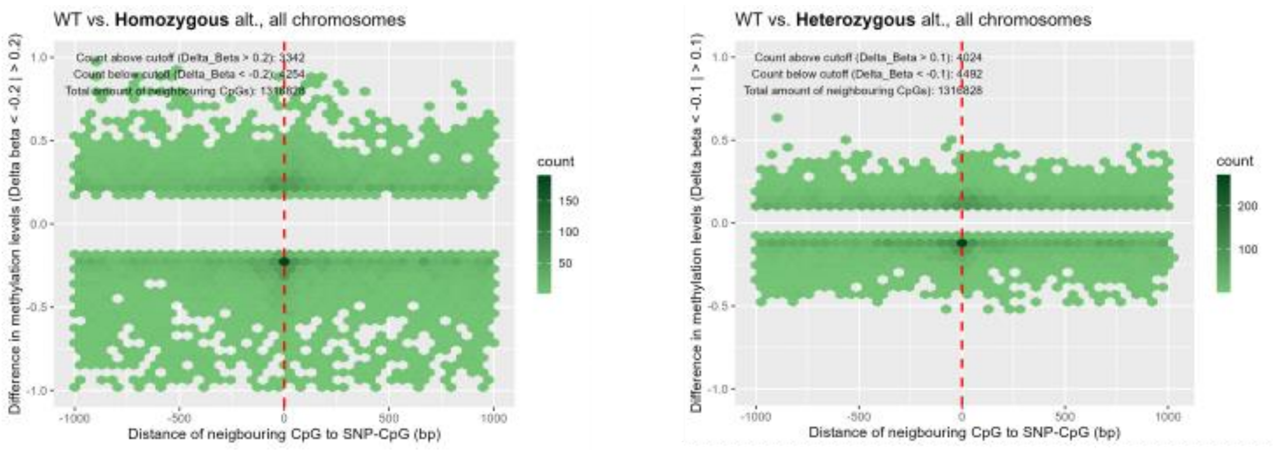

**Figure S3: Hexagon plots showing SNPs overlapping CpG sites that were also associated with a change in methylation levels of neighbouring CpG sites within a window of 1kb around the CpG sites.** The number of CpG sites represented by a single hexagon is indicated by the color scale depicted as side bar. A change in methylation levels was defined as an average change in beta-value of 0.2 for the homozygous, and 0.1 for the heterozygous groups. It can be appreciated that the effects of the homozygous variants are generally larger than the heterozygous variants. A decrease in methylation by the CpG disrupting SNP is more often found than an increase in methylation. Moreover, methylation of CpGs at very close distances to the CpG sites disrupted by the SNP is more frequently affected than methylation of CpGs at longer distances but the effect can extend throughout the window of 1kb and beyond.

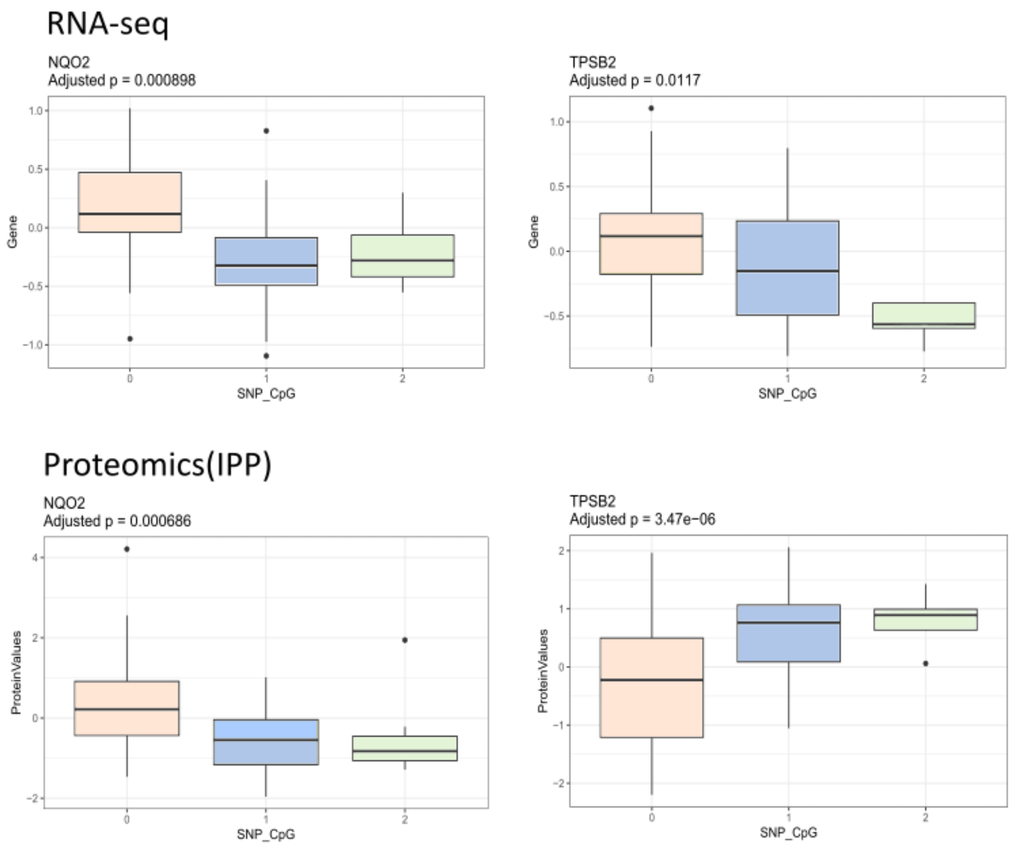

**Figure S4: Protein and mRNA levels of *NQO2* and *TPSB2* associated with SNP-CpG genotypes. SNP-CpG locations are chr6:3015873 and chr16:1277321, respectively.**

#### Hematological measurements

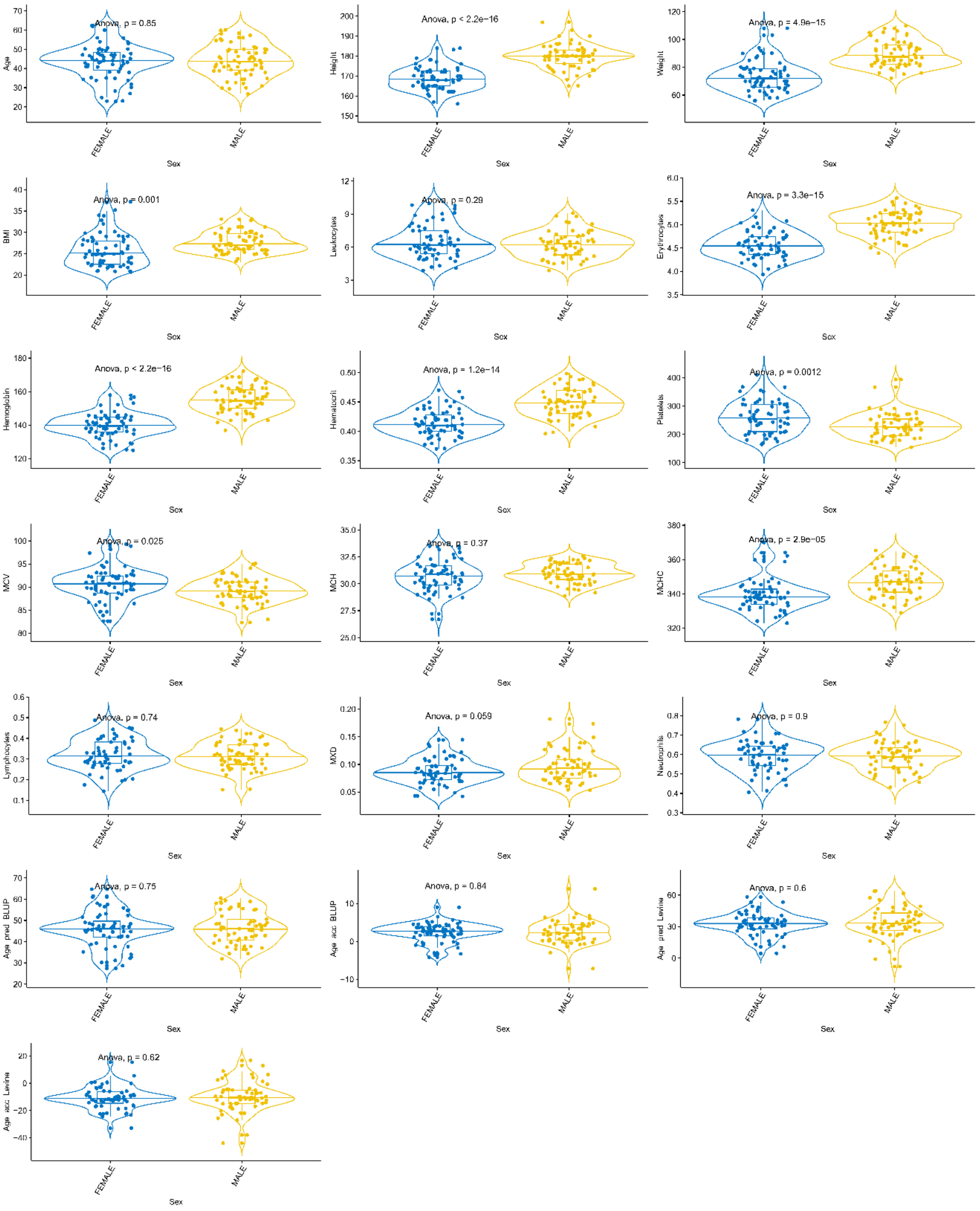

**Figure S5: Haematological measurements.** Female and male individuals are compared. Leukocytes are represented in units as x10E9/L cells; Haematocrit is a volume percentage (vol%) of red blood cells (RBCs) in blood; Erythrocytes are represented in units as [x10E12/L] cells; Platelets units are represented in units as [x10E9/L] cells; Haemoglobin values are represented as [g/L]; MCV = Mean Corpuscular Volume or Mean Cell Volume of RBCs in [fL/cell]; ...; MCH = Mean Corpuscular Haemoglobin in [pg/cell]; MXD = Mixed Cell Count, measures the combined levels of monocytes, eosinophils, basophils in the blood. The results are expressed as MXD (%); MCHC = Mean Corpuscular Haemoglobin Concentration is represented in [g/L]; Neutrophiles values are represented as a percentage (%) of all white blood cells (WBCs).

### EM-seq data

#### Estimated cell-type proportions

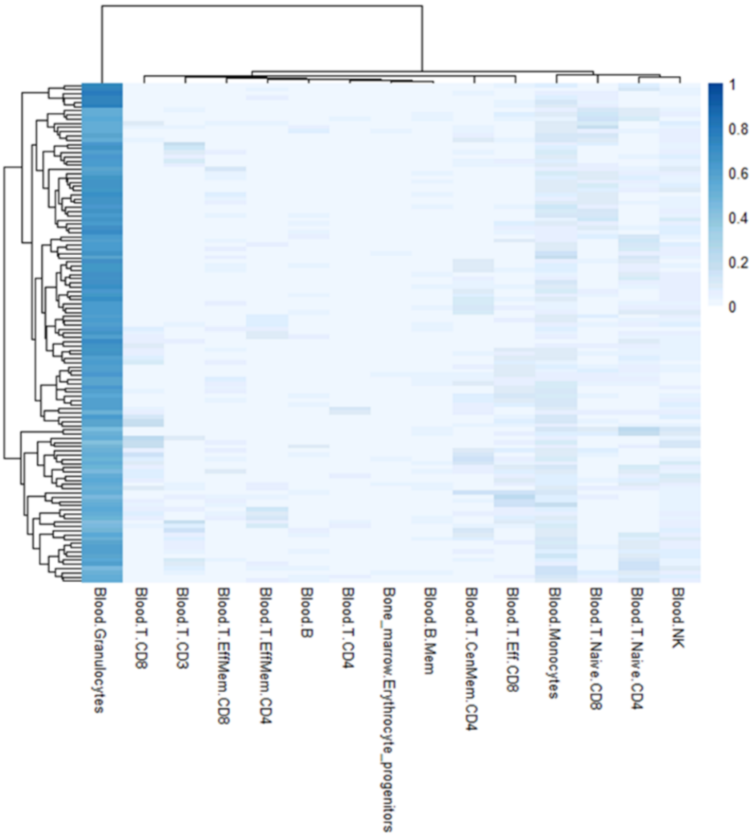

**Figure S6. Estimated cell-type compositions based on methylation levels in the healthy population.**

#### Methylation sites overlapping Single Nucleotide Polymorphisms (SNPs)

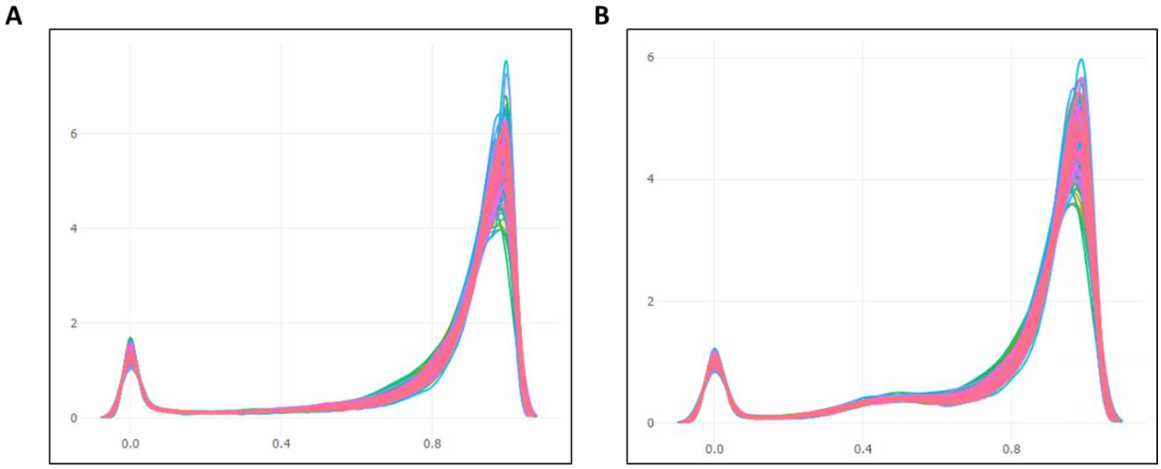

**Figure S7. Beta value distribution of 100.000 randomly selected sites in all samples based on A) all sites and B) sites with possibly overlapping SNPs. Note the peak around beta-values of 0.5 in panel B caused by heterozygous SNPs.**

### Metabolite levels compared in male/female subjects

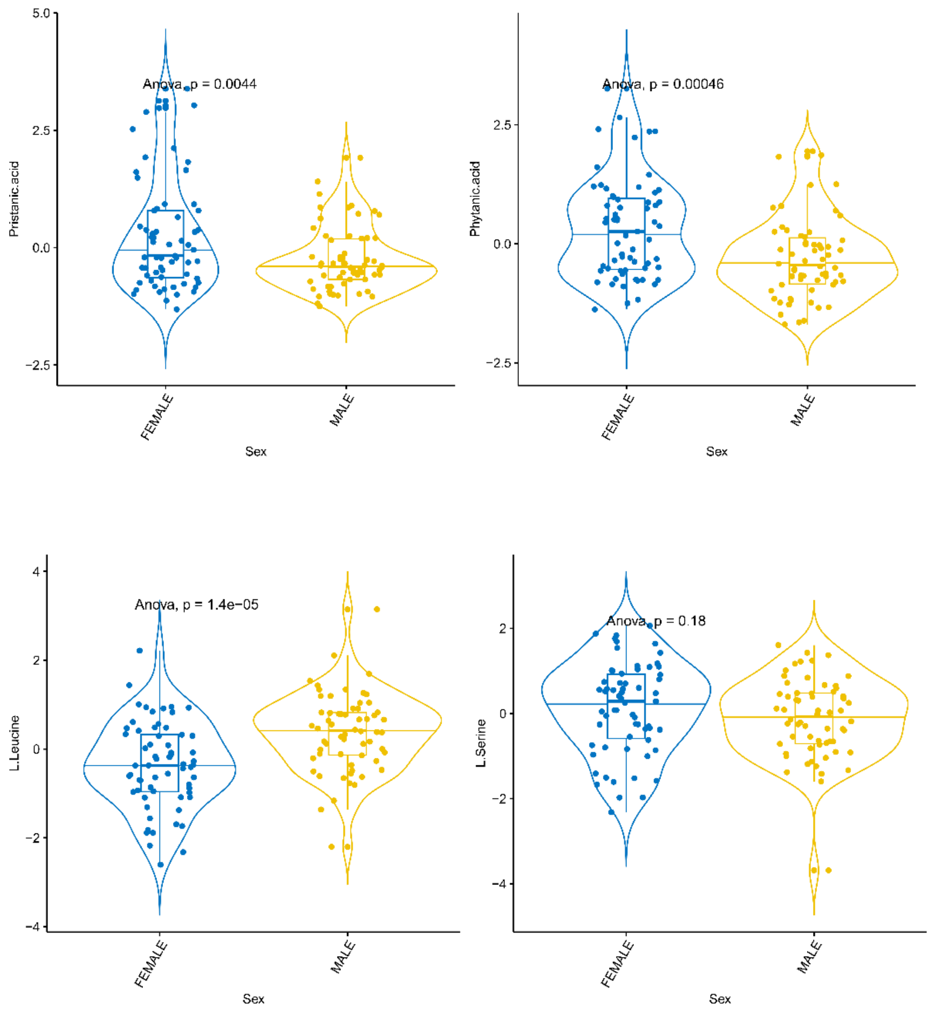

**Figure S8. Levels of phytanic acic, pristanic, leucine and serine compared among male and female subjects.**

#### Comparing platforms within one omics layer

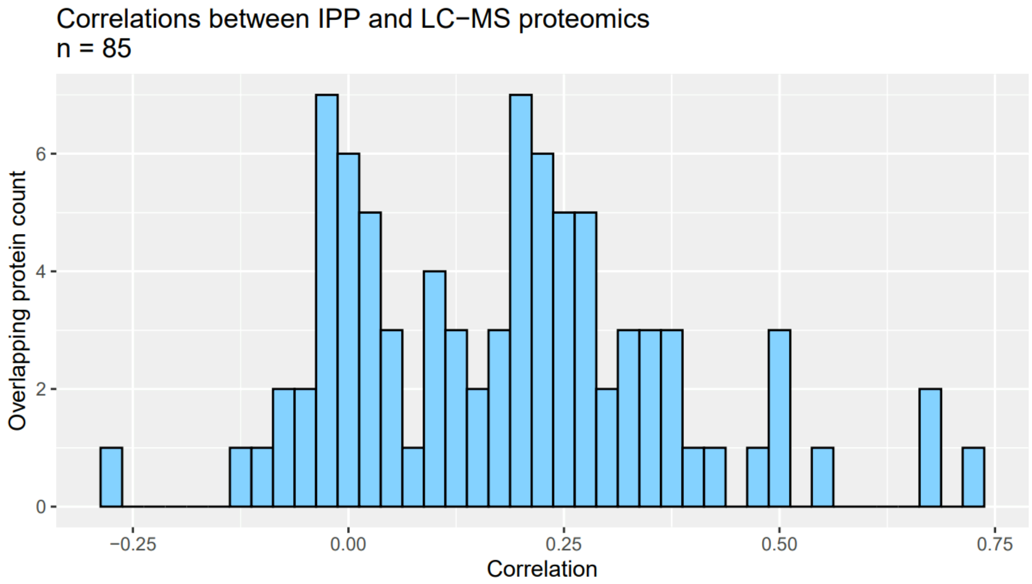

**Figure S9: Pearson correlations of proteins that were measured by LC-MS and IPP platforms.**

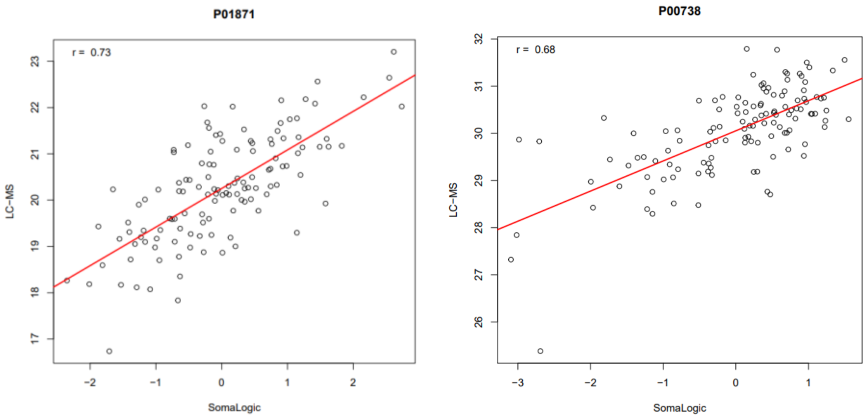

**Figure S10: Two strongest correlated proteins measured by LC-MS and Somalogic technologies.**

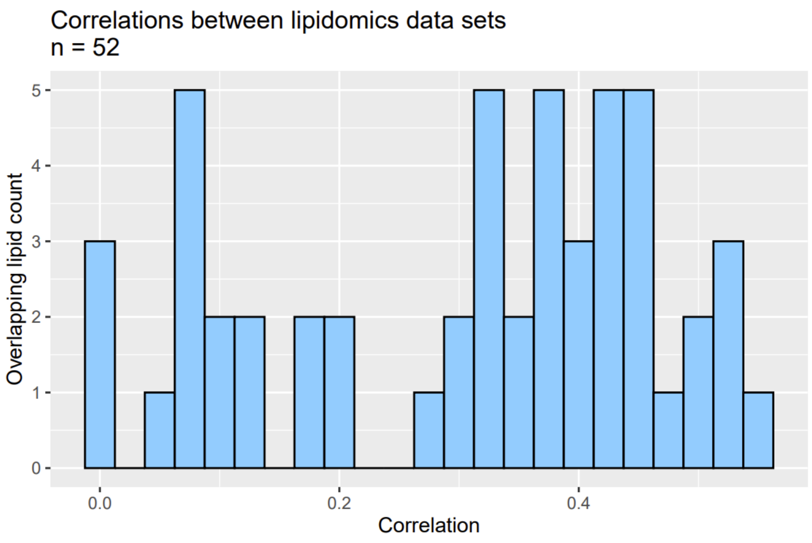

**Figure S11: Pearson correlations of similar lipids in the different lipidomics data sets.**

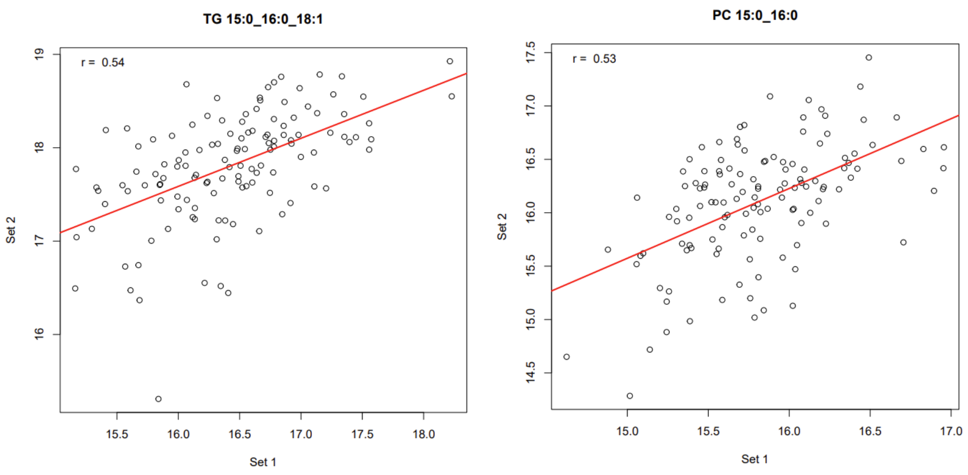

**Figure S12: Two strongest correlated lipids measured in the two different lipidomics data sets.**

### Principal Component Analysis

The correlations (unsignificant crossed out) between the first ten principal components of all –omics layers and phenotypic variables are summarized here.

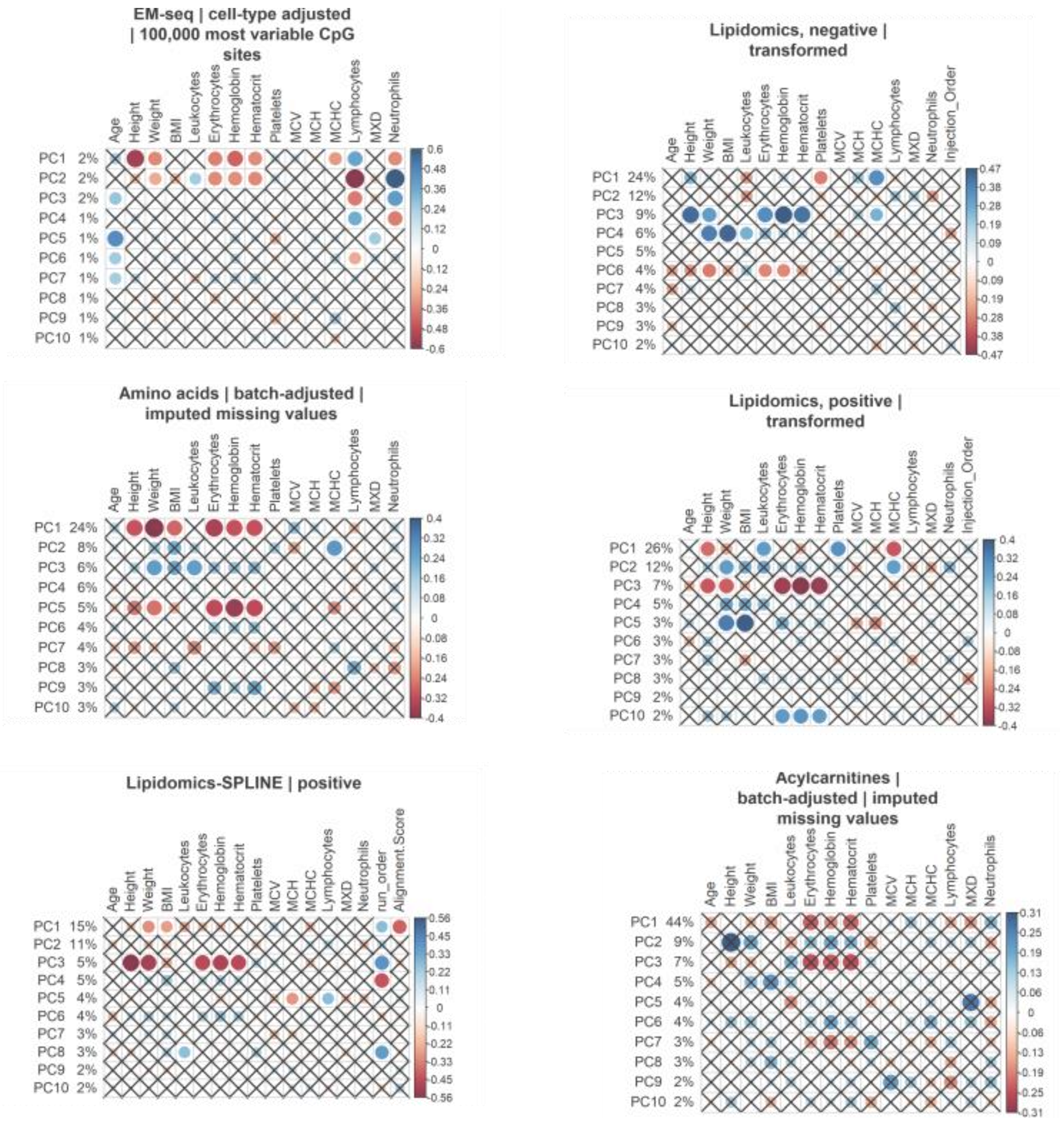

**Figure S13. Correlations between the methylation and transcriptomics data PCS and phenotypic variables**

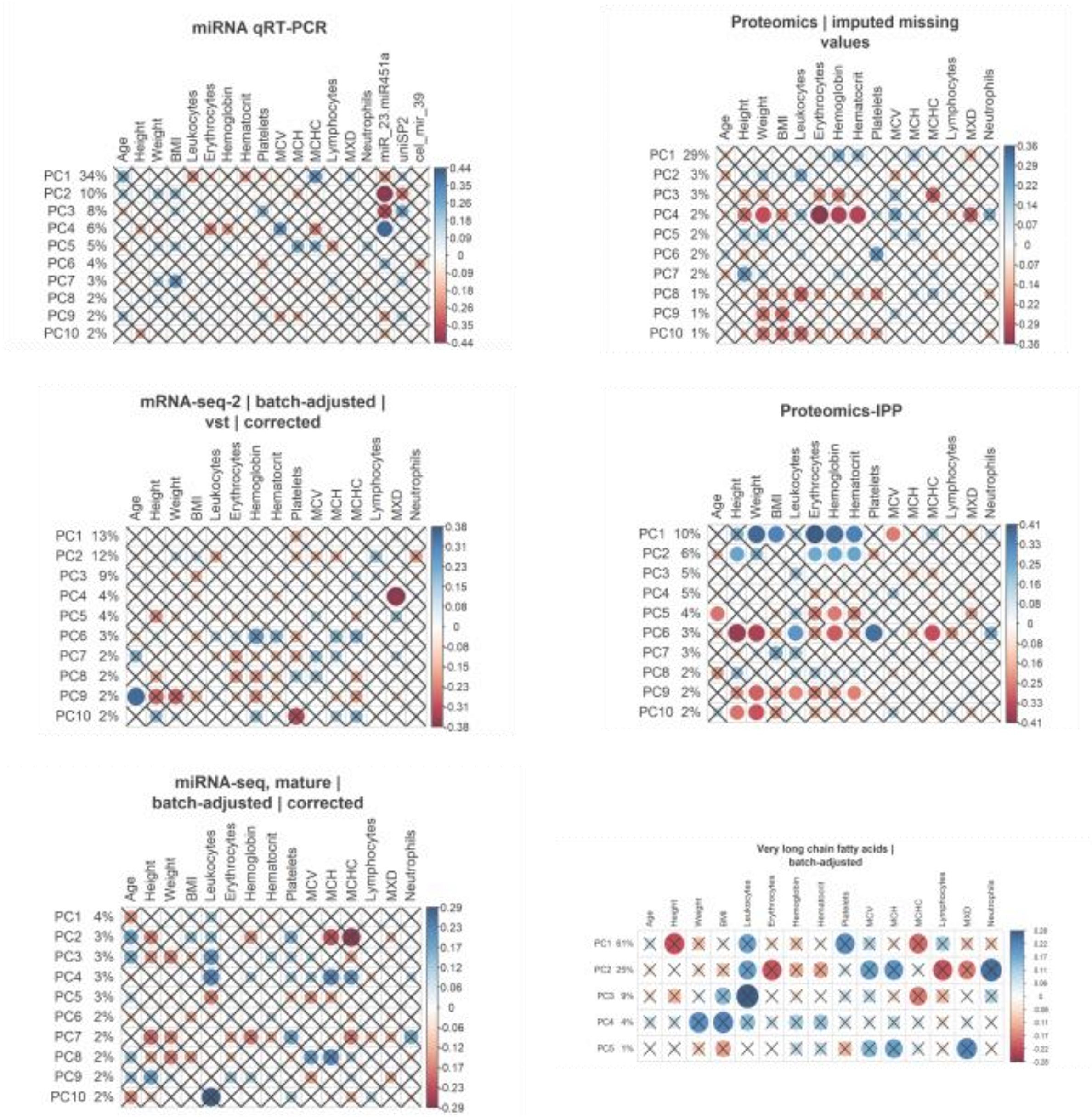

**Figure S14. Correlations between the mass-spectrometry based data PCS and phenotypic variables**

### Multi-Omics Factor Analysis (MOFA)

#### Pathway Enrichment Analysis

The weights of molecular features on the different MOFA factors were used as input for gene set enrichment analysis. Significantly enriched biological pathways, based on the ranked gene lists from mRNA, proteomics and metabolomics data, are shown in figure S17 –28.

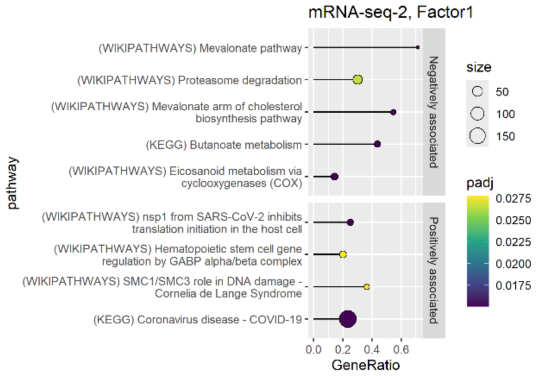

**Figure S15. Enriched pathways based on factor 1 weights.**

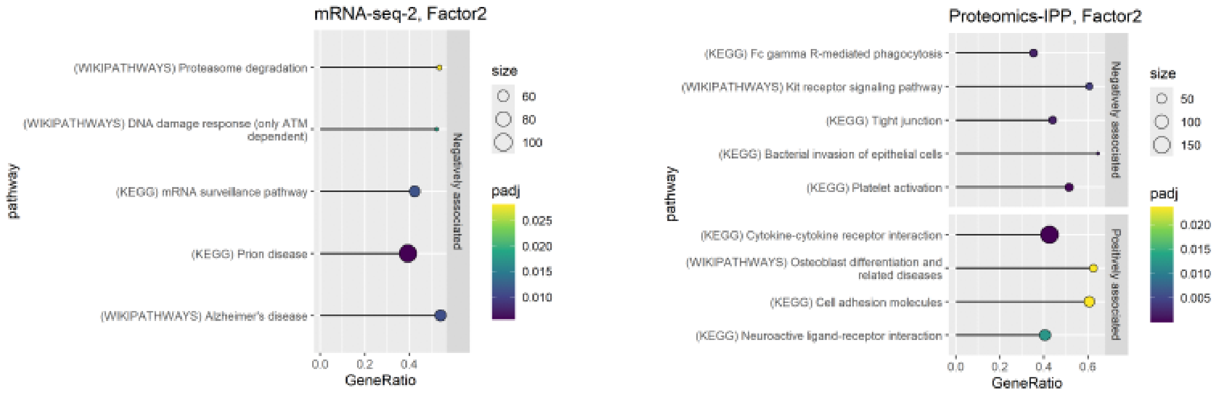

**Figure S16. Enriched pathways based on factor 2 weights.**

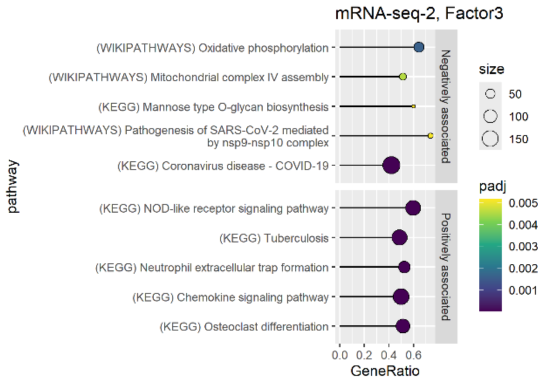

**Figure S17. Enriched pathways based on factor 3 weights.**

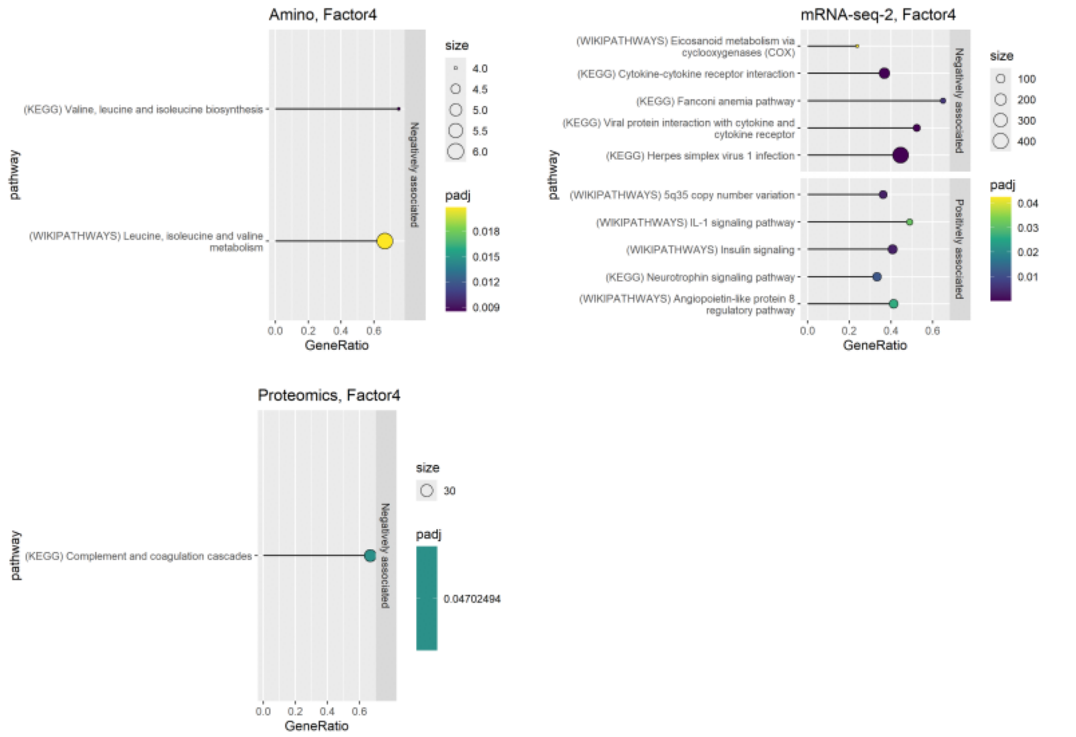

**Figure S18. Enriched pathways based on factor 4 weights.**

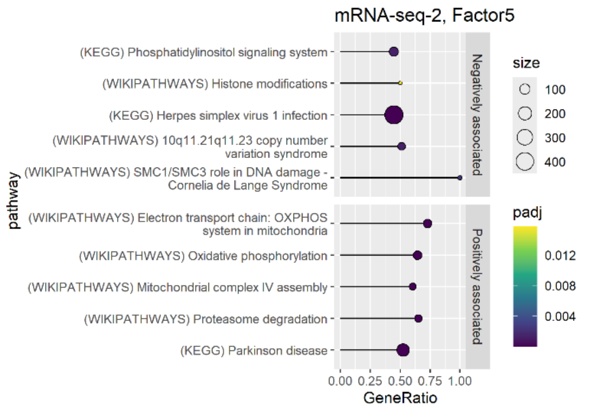

**Figure S19. Enriched pathways based on factor 5 weights.**

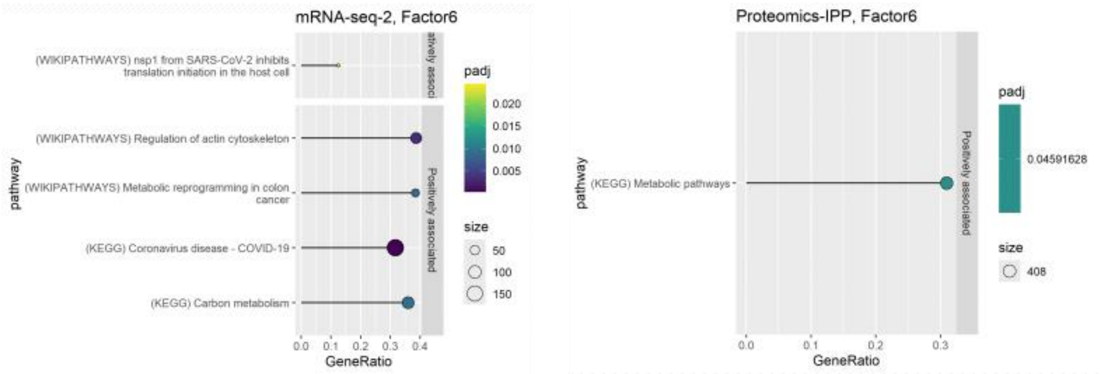

**Figure S20. Enriched pathways based on factor 6 weights.**

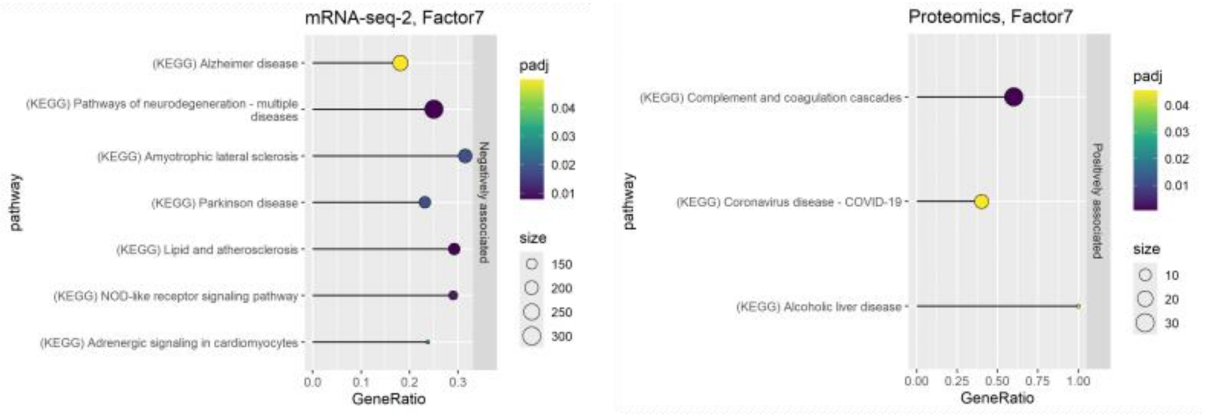

**Figure S21. Enriched pathways based on factor 7 weights.**

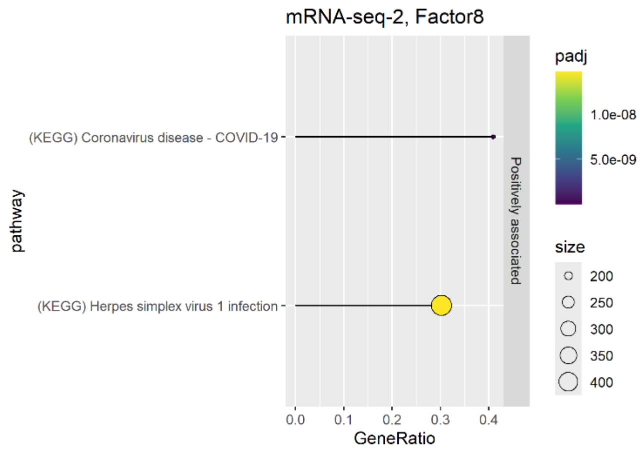

**Figure S22. Enriched pathways based on factor 8 weights.**

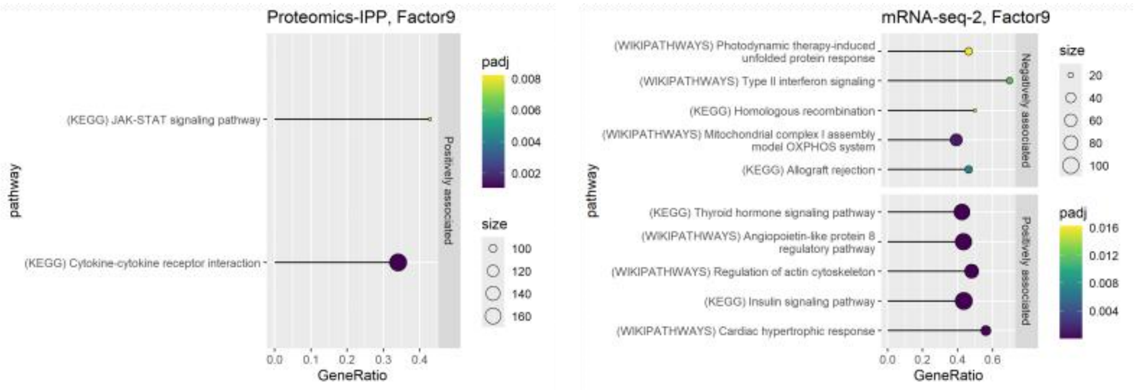

**Figure S23. Enriched pathways based on factor 9 weights.**

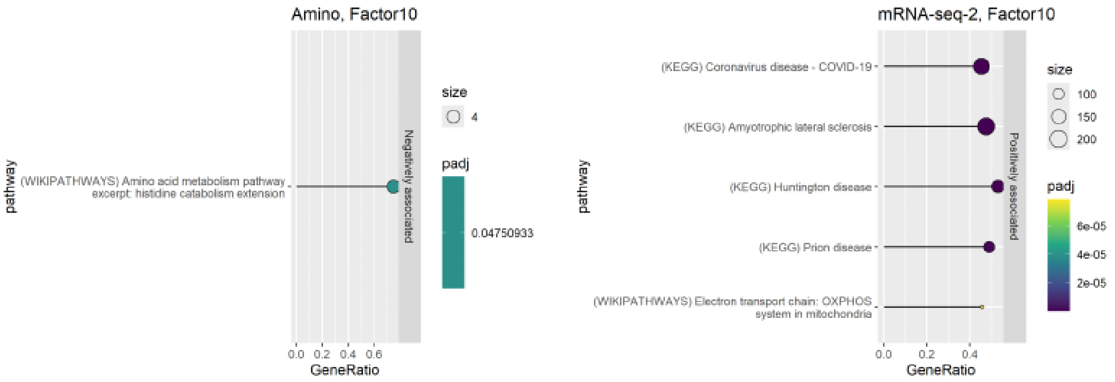

**Figure S24. Enriched pathways based on factor 10 weights.**

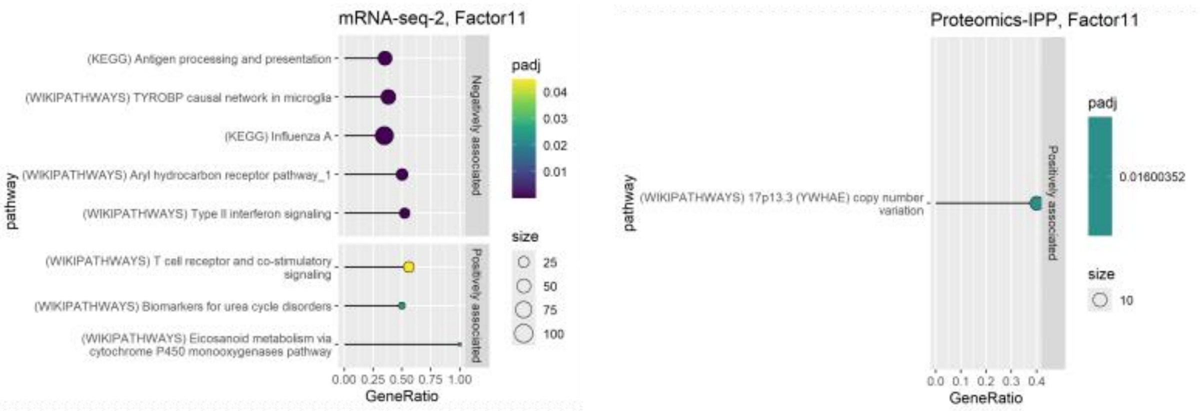

**Figure S25. Enriched pathways based on factor 9 weights.**

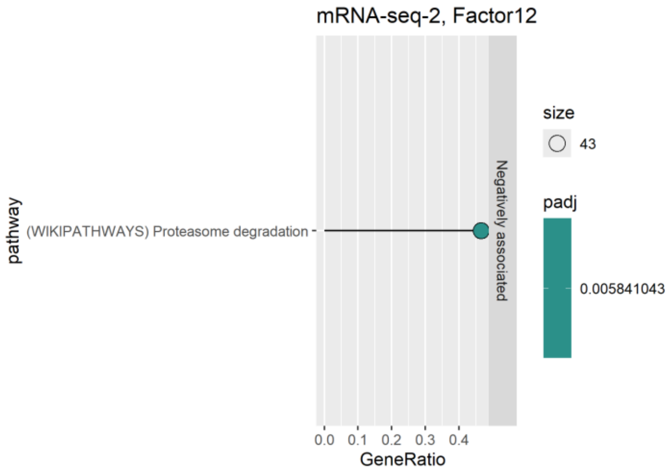

**Figure S26. Enriched pathways based on factor 9 weights.**

### MOFA scores and phenotypes

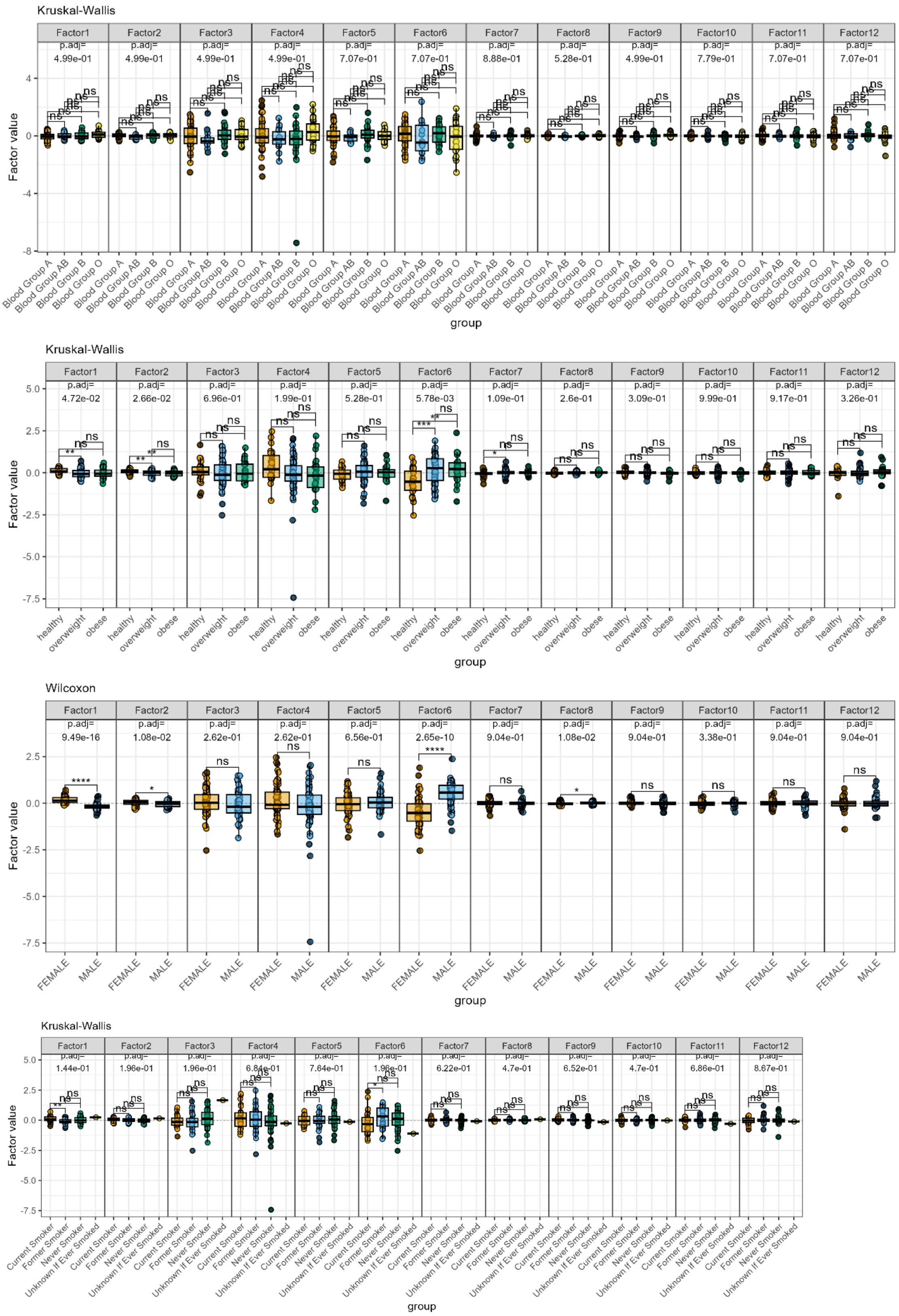

**Figure S27. MOFA factor scores and associations tested with categorial phenotypic variables.**

#### Cell –type specific genes

The mRNA feature weights on the MOFA factors were analyzed for the effect of cell type proportions. For each factor, the feature weights were sorted (ranked gene list) and the rankings of cell-type specific genes were plotted using histograms. Lists of cell-type specific genes were retrieved from The Human Protein Atlas (https://v19.proteinatlas.org) for B-cells, T-cells, dendritic cells, monocytes, granulocytes and NK-cells. Moreover, a list of 73 Ribosomal Protein genes (RPL/RPS) were included in this analysis. This revealed strong (negative or positive) contributions of cell-type specific genes on the MOFA factors are shown here.

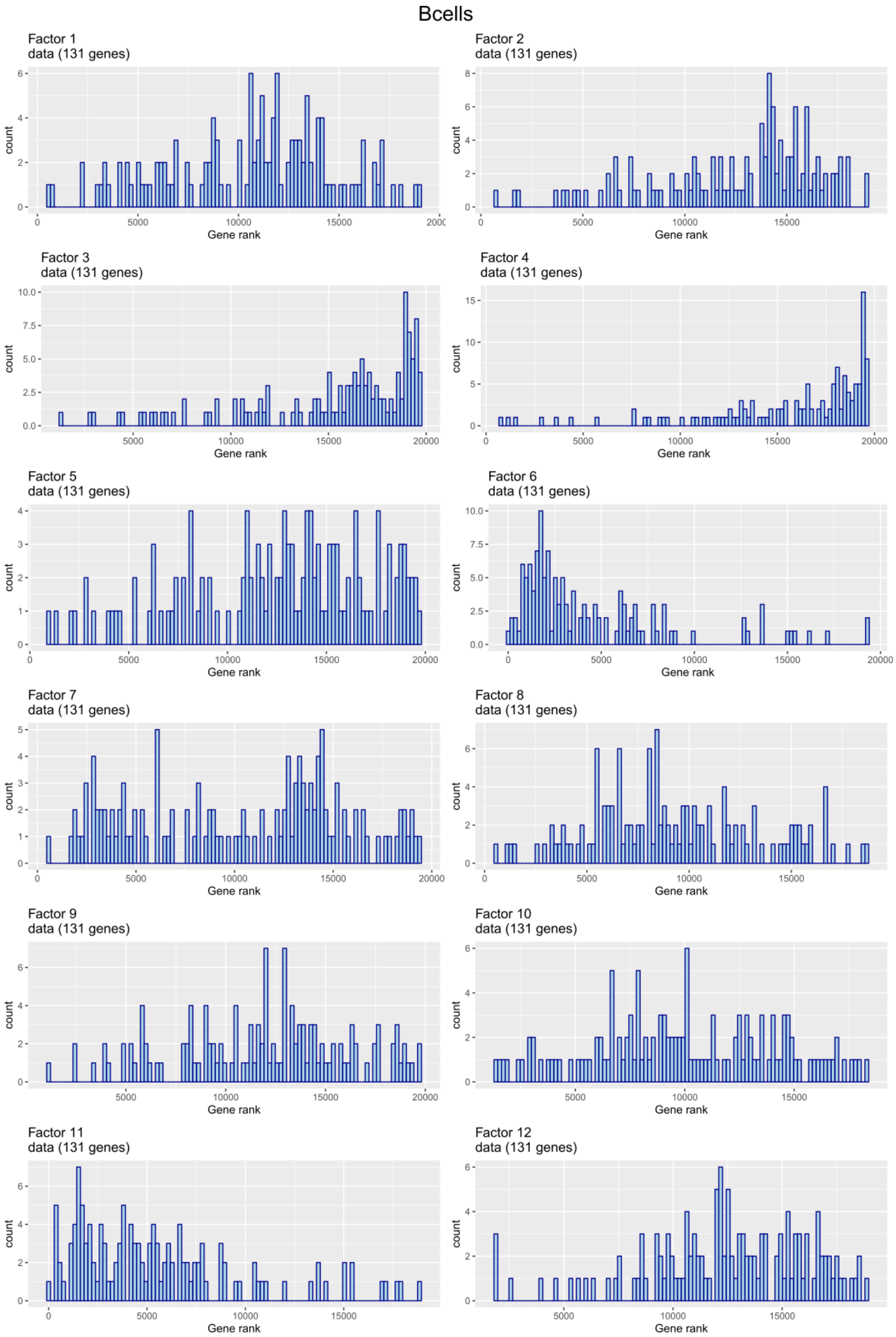

**Figure S28. Histograms depicting the distribution of gene ranks for B-cell specific genes in the mRNA-seq MOFA factor weights.**

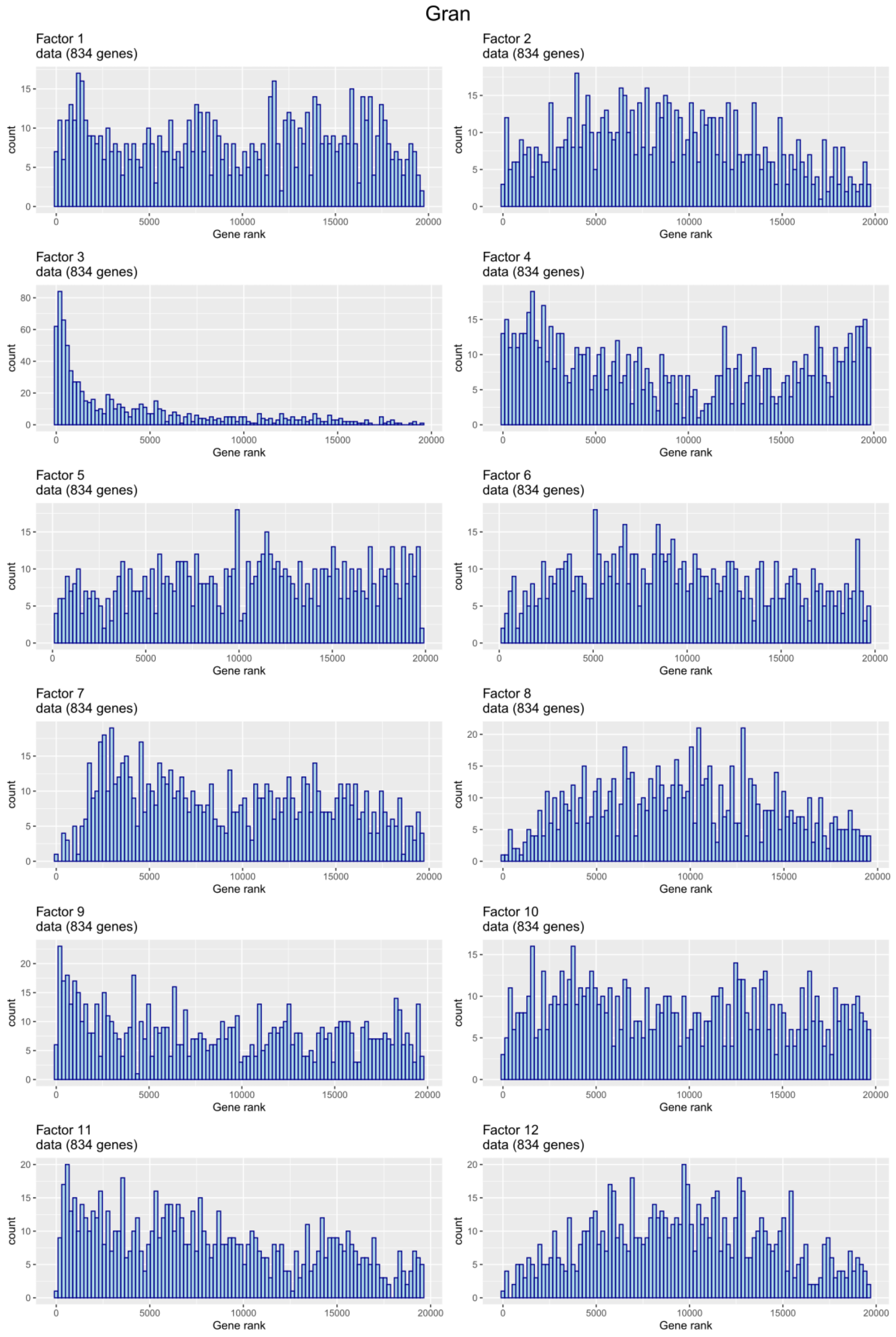

**Figure S29. Histograms depicting the distribution of gene ranks for granulocyte specific genes in the mRNA-seq MOFA factor weights.**

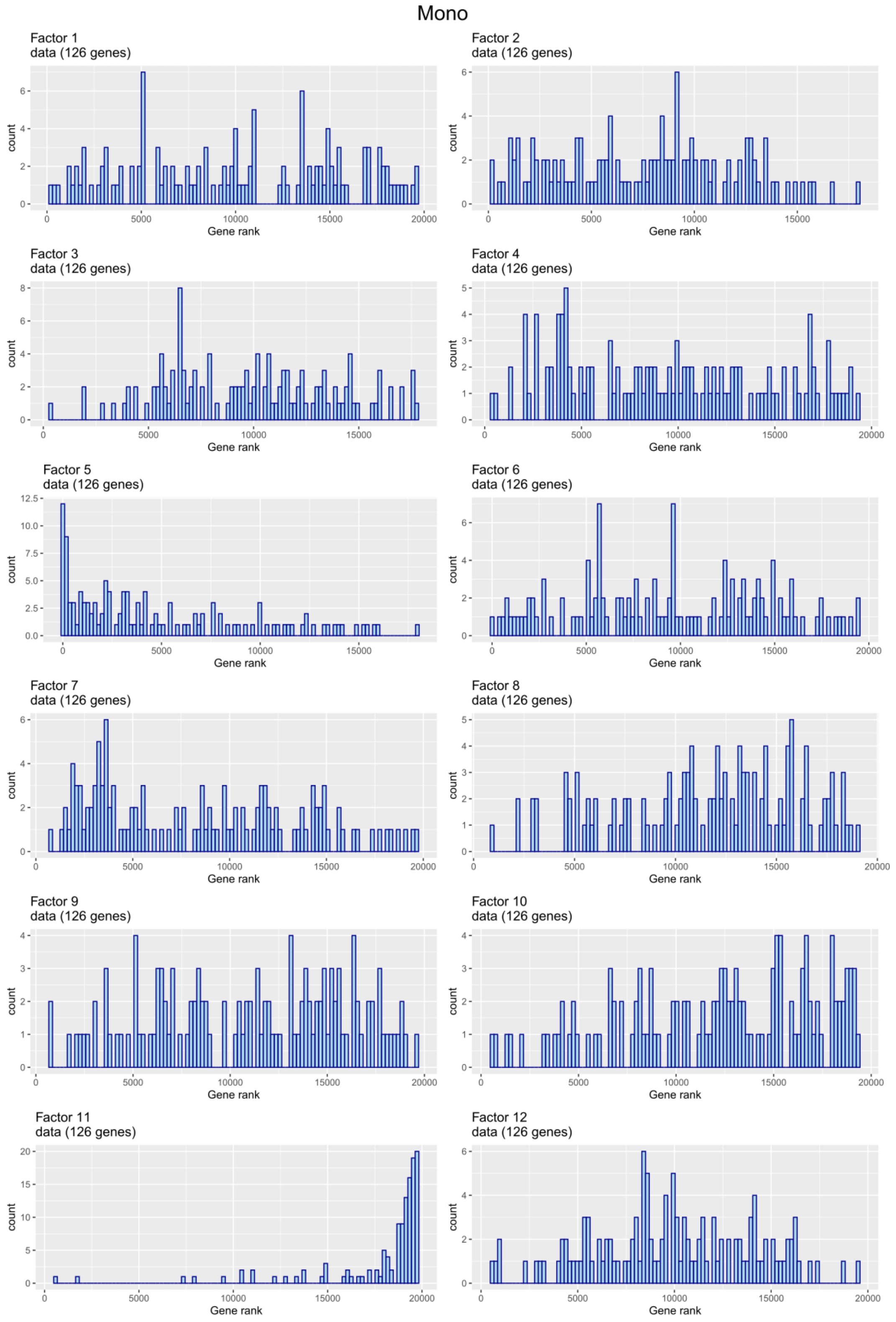

**Figure S30. Histograms depicting the distribution of gene ranks for monocyte specific genes in the mRNA-seq MOFA factor weights.**

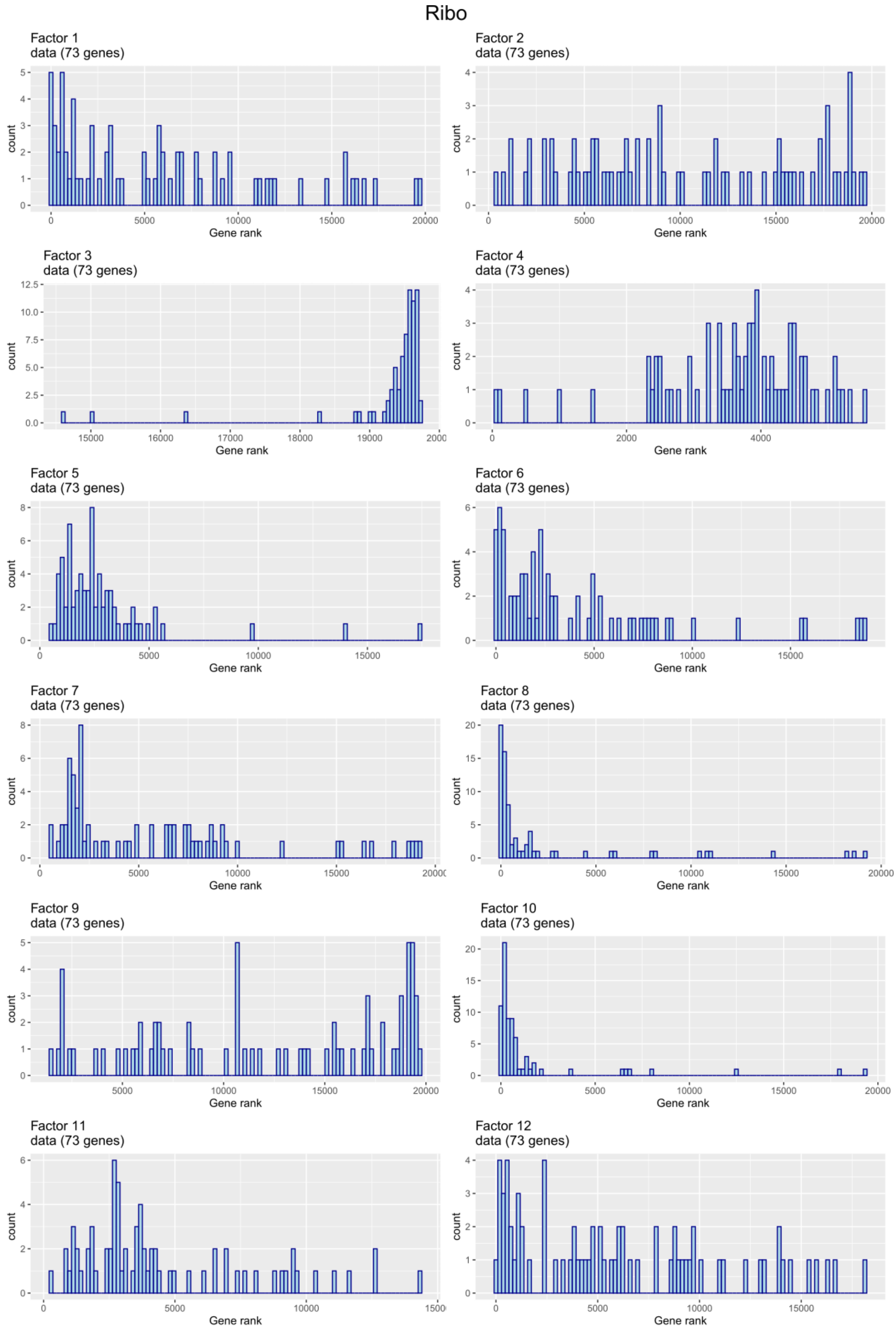

**Figure S31. Histograms depicting the distribution of gene ranks for the ribosomal genes (RPL and RPS) on the mRNA-seq MOFA factor weights.**

#### MOFA scores and gene aberrations

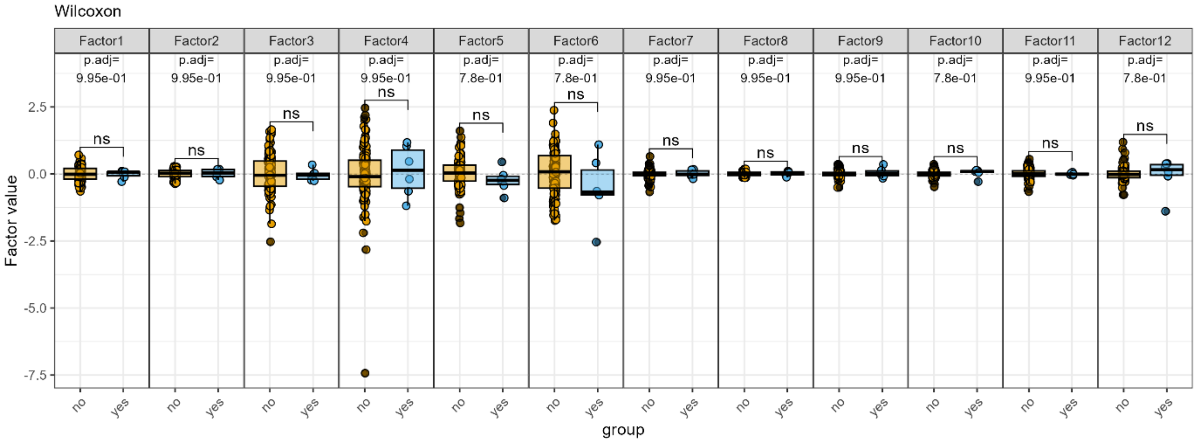

**Figure S32. MOFA factor scores compared among subjects with large numbers of genomic aberrations identified by array Comparative Genomic Hybridizations.**

### Similarity Network Fusion (SNF)

An alternative approach for integrating multi-omics data is to cluster the samples with Similarity Network Fusion (SNF). The SNF algorithm effectively combines information in the different –omics layers, creating a fused network of sample correlations (Wang, B. 2014). This method allows to determine whether the clustering of samples coming from healthy individuals improves when integrating multiple data types, compared to single –omics clustering.

SNF was systematically applied to all possible combinations of data modalities in this study, ranging from 2 to 12 data modalities per combination. Performance of all fused networks was evaluated using the silhouette score (Fig S26A), a metric of data similarity within clusters as compared to other clusters. Interestingly, the box plots show a clear decreasing pattern, suggesting that a higher number of omics combined results in worse clustering. Based on this metric, one fused network clearly outperforms all others, which is the combination of lipidomics negative and lipidomics positive. Since the lipidomics datasets are result of LC-MS analyses of the same biological samples in different polarity modes there are many compounds present in both. This explains the high fusion level. The second-best performance was observed in the fusion of lipidomics (including VLCFA) data, in which two clusters were found by SNF (Fig S26C). To find more biological meaning with respect to these SNF clusters, differences in categorical phenotypes among the SNF clusters were visualized (Fig S26d), of which chi-square tests revealed a significant different among males and females in these clusters. Moreover, differences in continuous variables were tested using Mann-Whitney U tests, revealing no significant relations.

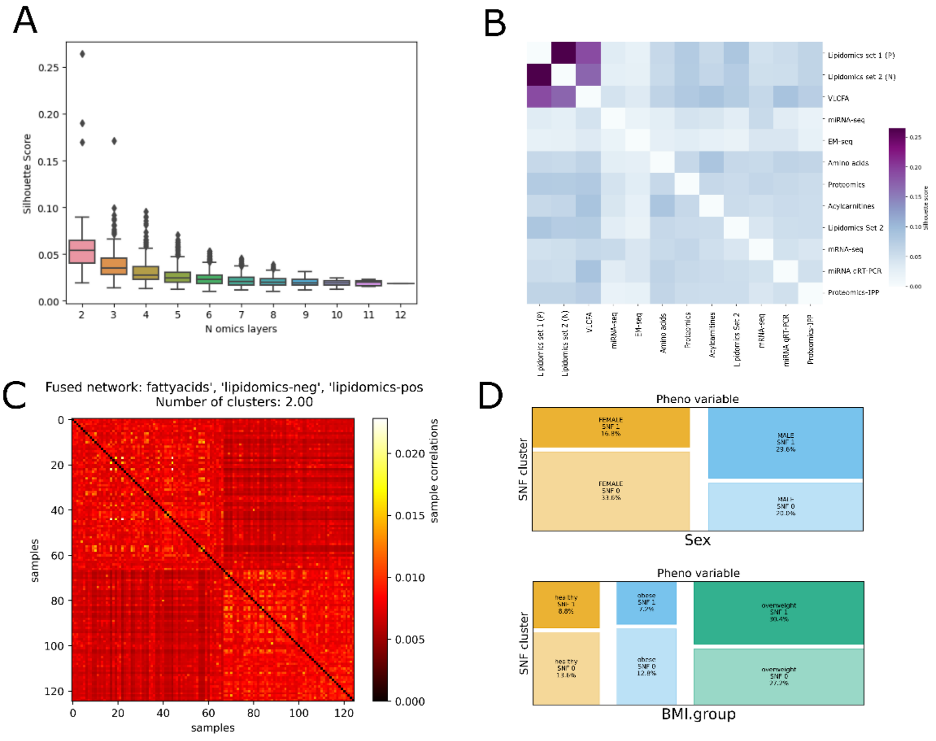

**Figure S33 Similarity Network Fusion (SNF)**: **A) Silhouette scores, reflecting the distance between clusters (1 is complete separation), for every fused network are plotted, with the number of –omics layers that are fused on the x-axis. B) Heatmap of the silhouette scores in pairwise –omics clusterings. C) The sample correlations in the fused network based on VLCFA, lipidomics (neg and pos) data. D) Mosaic plots showing how the categorical phenotypic traits (Sex and BMI group) are distributed over the two sample clusters discovered by SN**F.

#### Partial Least Squares PLS2

Partial Least Squares 2 (PLS2) was applied to evaluate pairwise correlations across the various –omics layers. PLS2 is a multivariate method that seeks linear relations between two continuous data modalities. It has proven to be specifically efficient with data that has a small number of observations as compared to the number of features, such as most –omics data. As illustrated in Figure S9, PLS2 reveals numerous high correlations. The lipidomics data in both negative and positive modes demonstrated strong correlations. This is not a surprise, since there are many compounds ionisable in both polarities. The two miRNA data sets expressed strong correlation correlations as well. Also, the high-dimensional miRNA-seq and EM-seq data displayed strong correlations. This observation may be attributed to the feature selection process implemented in PLS2 (see Methods). The features that correlate highest with the paired –omics layers will be selected for each model; hence this could explain the high correlations with these high dimensional data sets.

These correlations confirmed some of the findings that were done in previous analyses. For example, the high correlations between the methylation levels and both proteomics and metabolomics data were also reflected by similar associations in the linear models and PCA. Moreover, the high correlation of mRNA-seq and VLCFA data was also found by MOFA with MOFA factor 7.

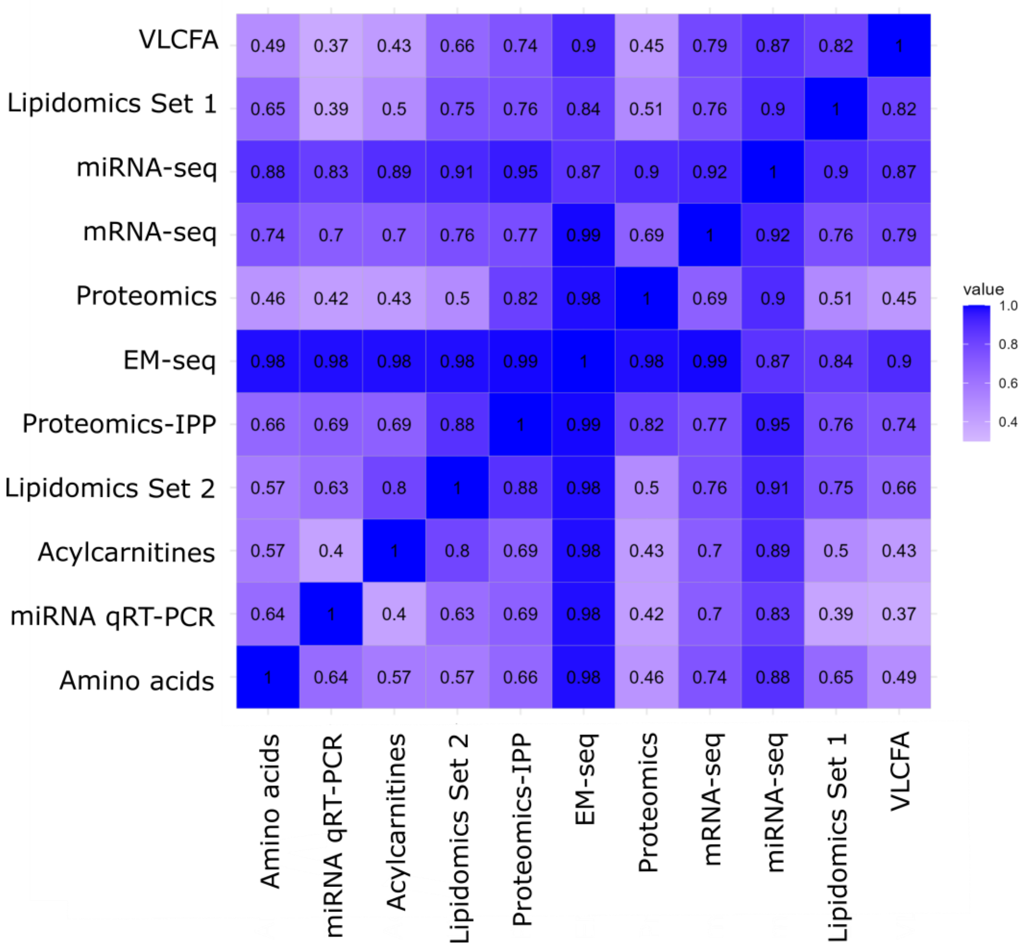

**Figure S34 Partial Least Squares (PLS2). Pairwise correlations of the different data modalities. The –omics layers are clustered using the complete linkage method.**

### Aging

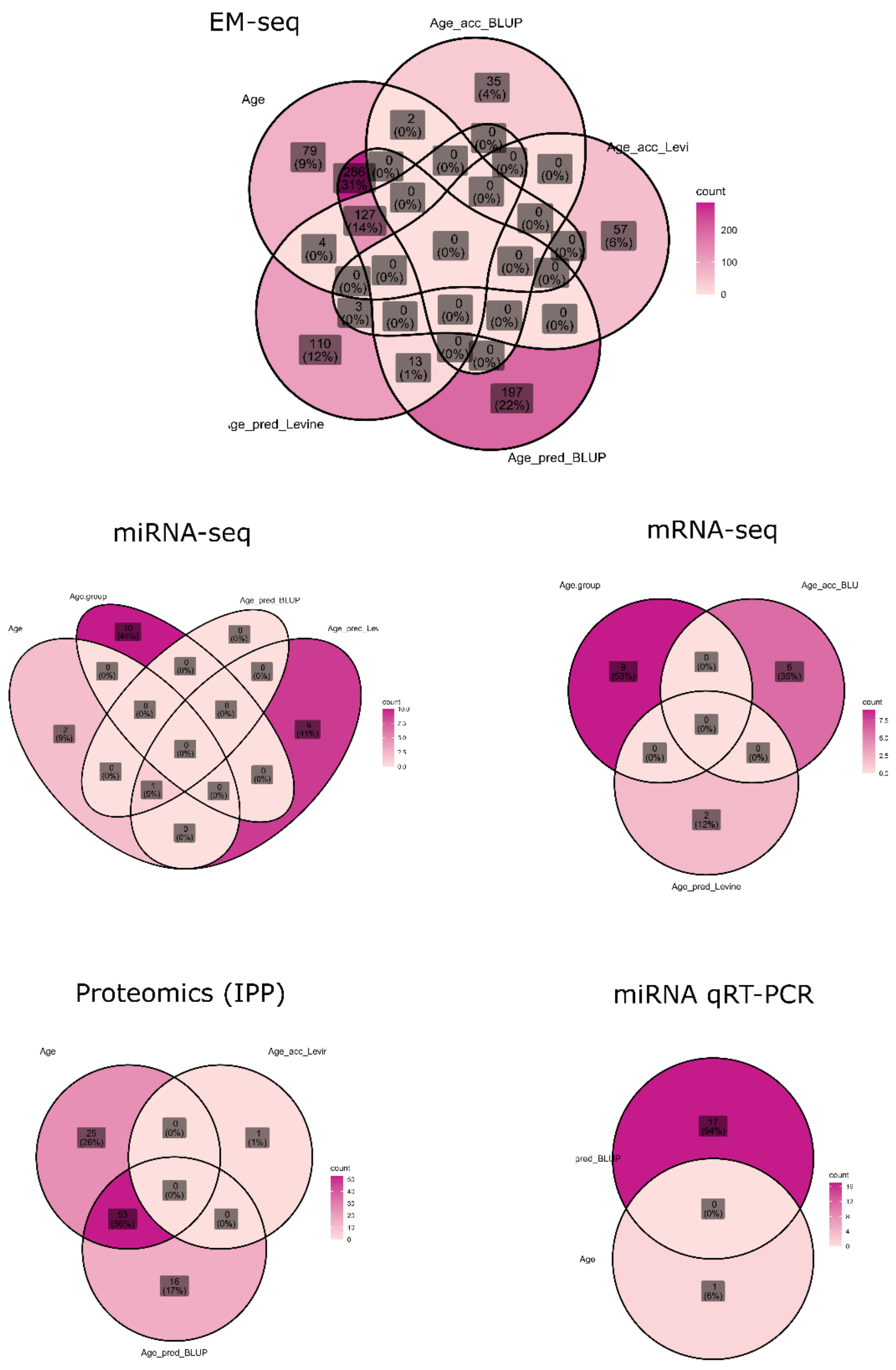

**Figure S35. Overlap between age-associated omics features. Overlap was checked among all omics features that associated with either chronological age, predicted age (BLUP and Levine) and age acceleration (BLUP and Levine).**

### Age acceleration

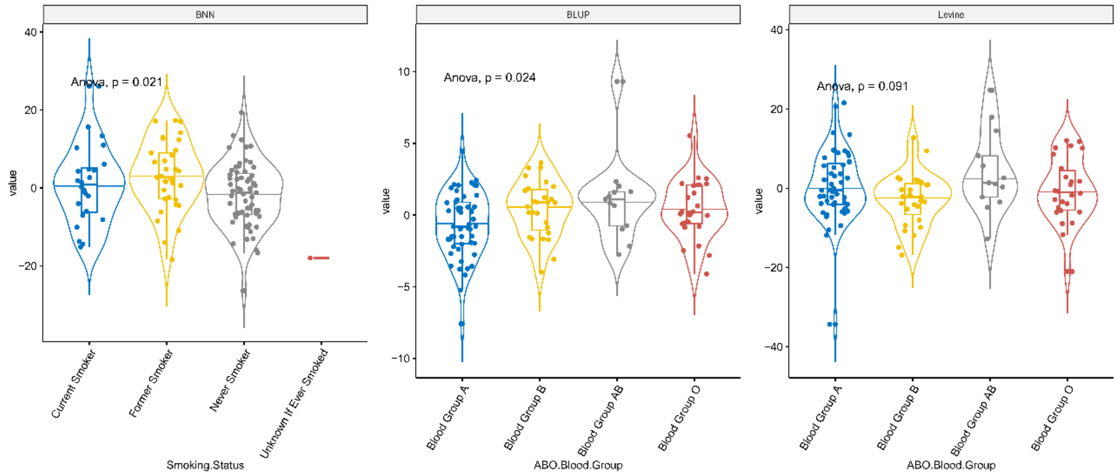

**Figure S36. Phenotypic groups compared on age acceleration levels.**

### Epigenetic clock Levine

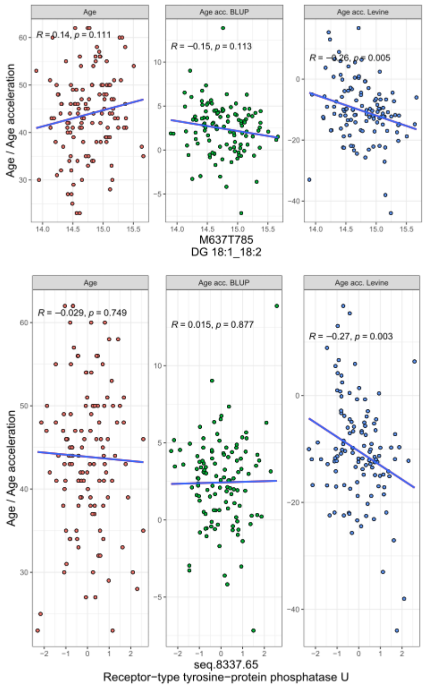

**Figure S37. Individuals –omics features associating with the age acceleration based on Levine clock.**

#### Epigenetic clock BLUP

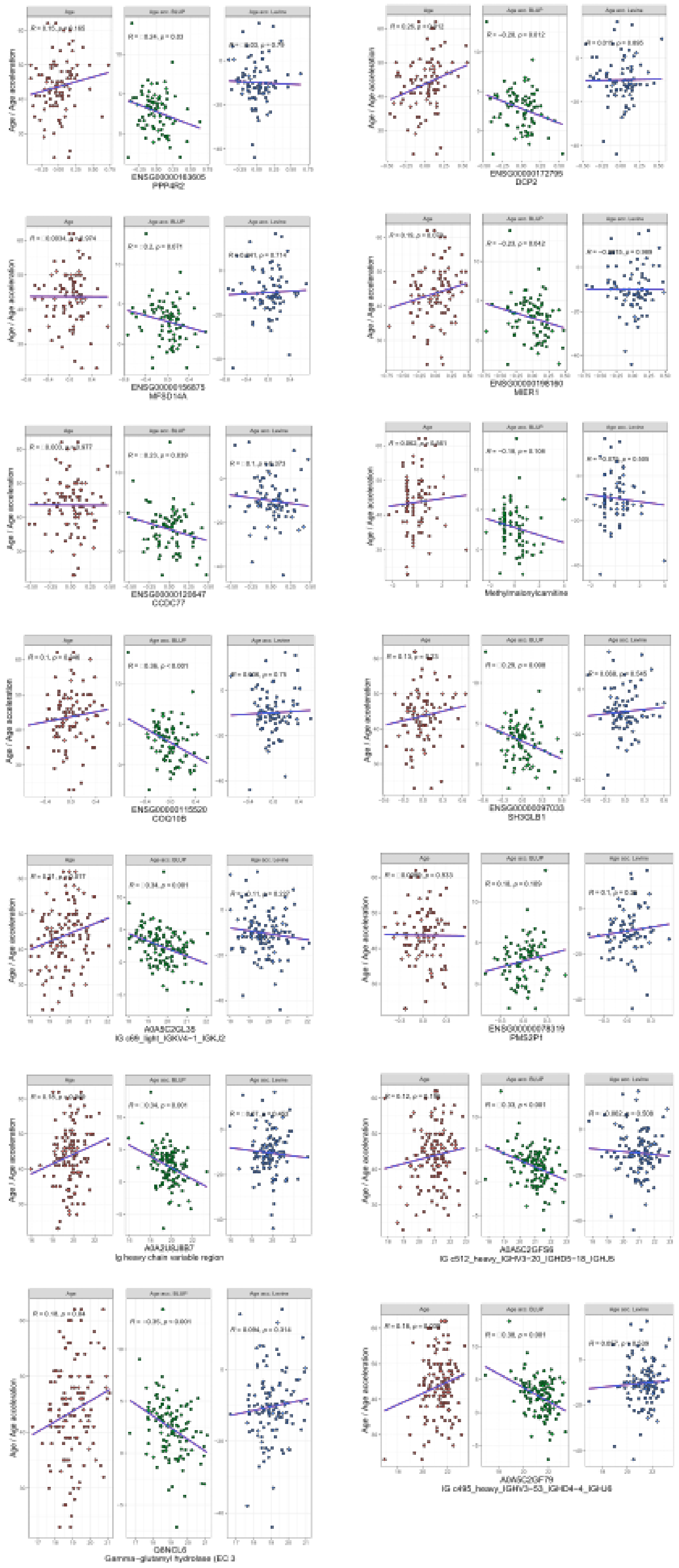

**Figure S38. Individuals –omics features associating with the age acceleration based on BLUP clock.**

### Batch effect adjustment

#### Transcriptomics (mRNA-seq)

Batch effect adjustment was performed using *ComBat* from R library *sva* (version 3.44.0). The parametric method using sex as covariate was applied. Sample correlations were visualized using a heatmap (Fig S1), and Principal Component Analysis was applied to visualize the differences before and after batch-effect adjustment (Fig S2)

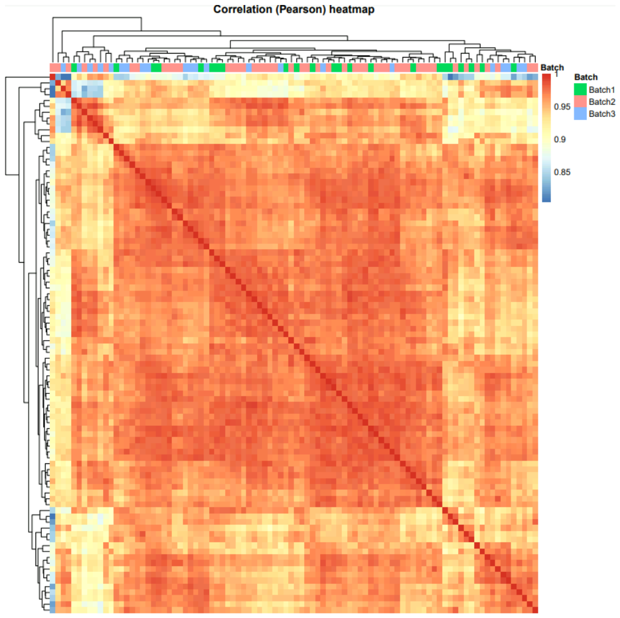

**Figure S39. Correlation heatmap. Batches are indicated by colors.**

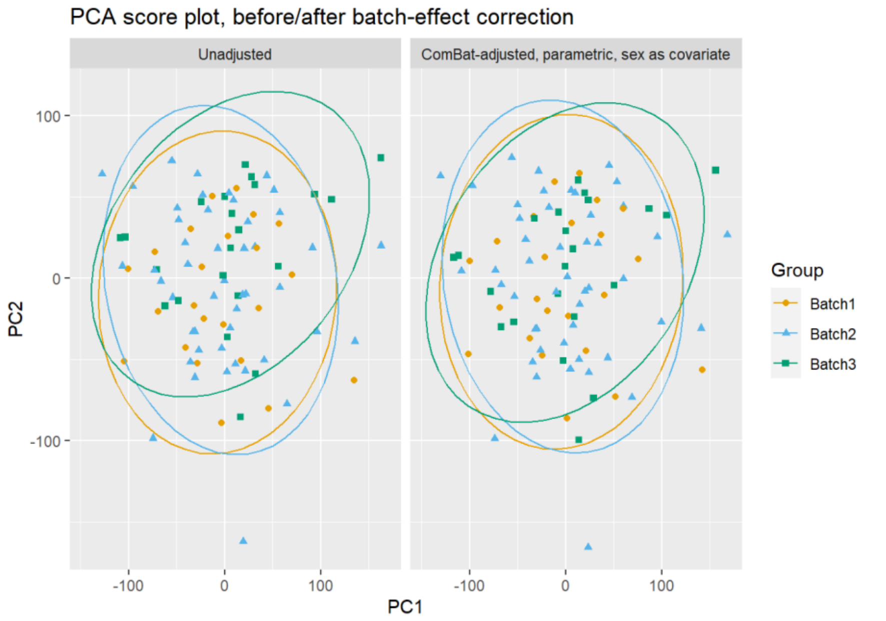

**Figure S40. Principal Component Analysis showing correction for batch-effect.**

#### Transcriptomics (microRNA-seq)

Batch effect adjustment was performed using *ComBat* from the R package *sva* (version 3.44.0). The parametric method using sex as covariate was applied. Batch 6 consists of only one single sample, through which it is not possible to include in the batch-correction. Sample correlations were visualized using a heatmap (Fig S3), Principal Component Analysis was applied to visualize the differences before and after batch-effect adjustment (Fig S4).

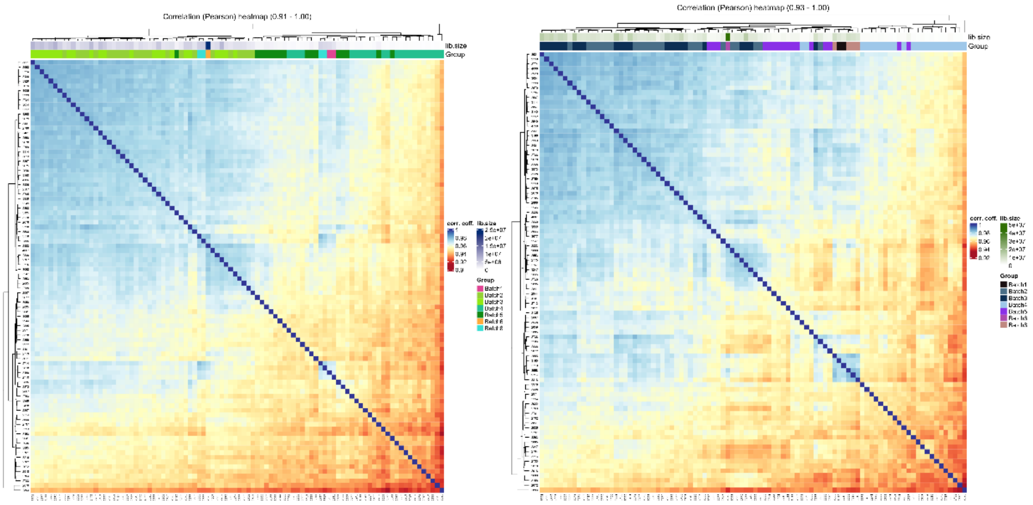

**Figure S41. Sample correlation heatmap for hairpin (left) microRNAs and mature microRNAs (right)**

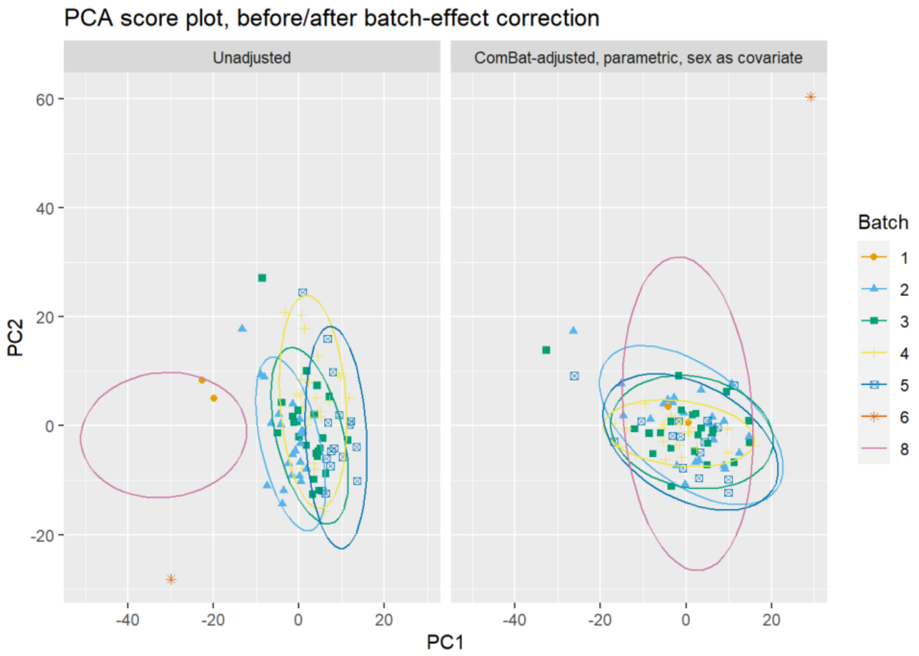

**Figure S42. Principal Component Analysis showing correction for batch-effects.**

#### Transcriptomics (microRNA qRT-PCR)

To remove the batch effect, the Combat method (sva R package) was used. To test the effect of the batch effect removal, the PERMANOVA test was performed again. After batch-adjustment, both RT and extraction were not associated with the microRNA levels (Fig S5).

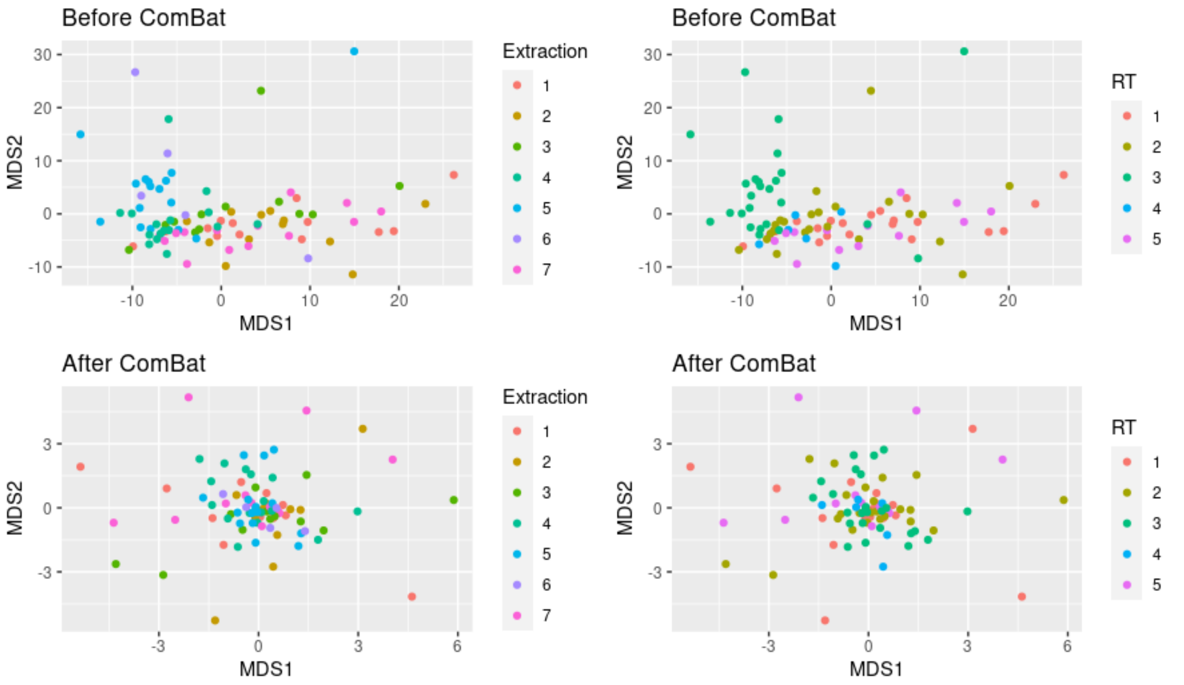

**Figure S43. Correction for batch-effects using Combat.**

#### Proteomics

Batch uncorrected dataset with missing values was used as an input of batch correcting process. Partial redundancy analysis (RDA), implemented in vegan R package (Oksanen J et al., 2022), estimated a proportion of constrained variability (Sex factor) as 5.3 % while a proportion of conditioned variability (Batch) as 37.6 %. This strong batch effect is also visible in graphical representation of batch uncorrected data in form of PCA plot (for PCA the dataset with imputed values was used) with confidence regions colored by batch.

Three methods of batch correction – 1) the feature-level median method, implemented in proBatch R package (Cuclina, J., 2018), 2) the ComBat method and 3) the Combat method with Sex factor as a covariate, both implemented in sva R package, were used for batch correction. The best results were obtained for the last method – a proportion of constrained variability (Sex factor) 2.3 % and a proportion of conditioned variability (Batch) 0.3 % (Fig S6, S7).

In the case of unbalanced batch design using univariate models with adjustment to batch factor might be more appropriate than batch correction of the data but it is not convenient to use batch uncorrected data for multi-omics integration and analyses. To choose the best batch corrected dataset the power of discrimination between male and female samples for each of the four datasets with usage of Support Vector Machines (SVM) models, implemented in e1071 R package (Meyer D et al., 2023), and LASSO models, implemented in glmnet R package (Friedman J et al., 2010), was explored.

Samples were divided into 5 folds and for each i-th fold, i = 1, 2, ..., 5, the separate models of sex and batch were fitted on the rest 4/5 of samples (training data), tested on the i-th fold samples (test data) and expressed as error rates. For both multivariate modelling methods – SVM and LASSO, the best behavior was observed for ComBat with Sex covariate batch corrected dataset. Models based on this dataset fitted sex well (SVM models, test error rates = 8 %, 21 %, 16 %, 0 %, 0 %; LASSO models, test error rates = 16 %, 13 %, 16 %, 7 %, 13 %) while batch with high error rates: SVM = 100 %, 100 %, 100 %, 100 %, 96 %, LASSO = 76 %, 71 %, 81 %, 59 %, 71 % which means that the dataset kept the Sex factor influence while the batch effect has been removed.

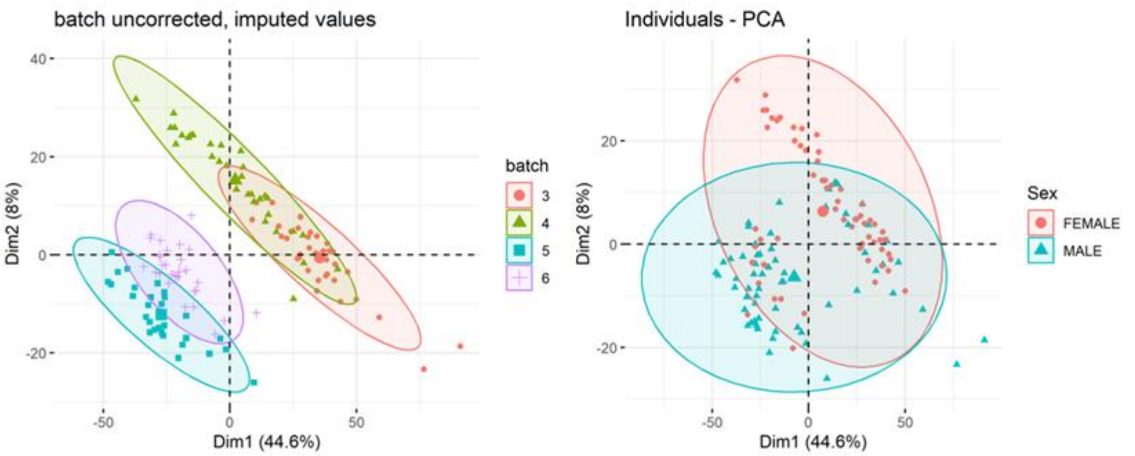

**Figure S44: PCA plot (first two components) for batch uncorrected dataset with imputed missing values; confidence regions colored by batch factor and Sex factor.**

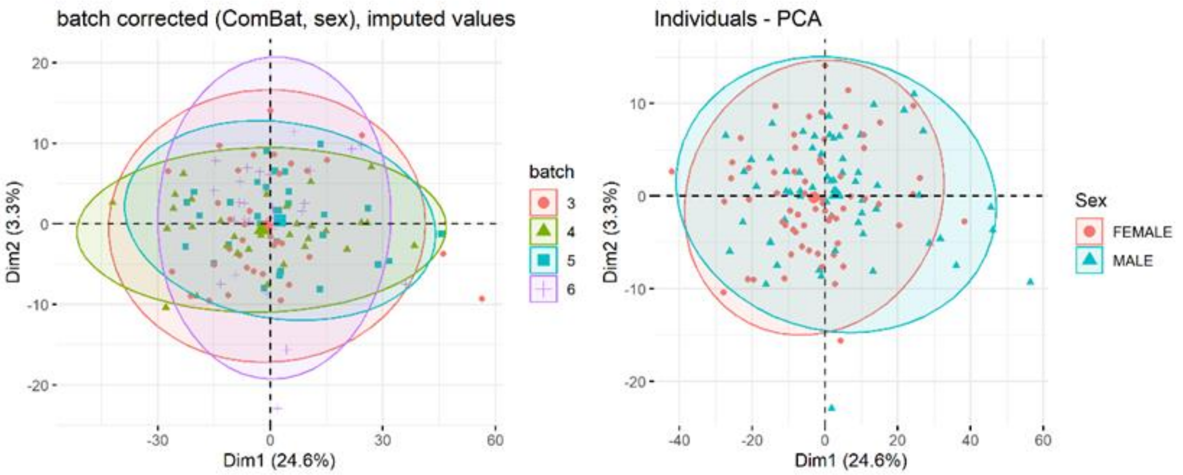

**Figure S45: PCA plot (first two components) for ComBat with Sex as covariate batch corrected dataset with imputed missing values; confidence regions colored by batch factor and Sex factor.**

#### Metabolomics

The presence of batch effects was assessed visually with sample correlation plots (Fig S8) and principal component analysis (PCA) (Fig S9), and statistically using analysis of variance (ANOVA) and redundancy analysis (RDA) (Table S1-2). Sample correlations were calculated in order to detect the presence of potential batch effects. Pearson correlation coefficient was calculated using complete pairwise observations. Batch effects were detected using Analysis of variance (ANOVA) by comparing single metabolite levels of the different batch groups. This was performed for each metabolite measured by the different assays (ACRN, AA and VLCFA).

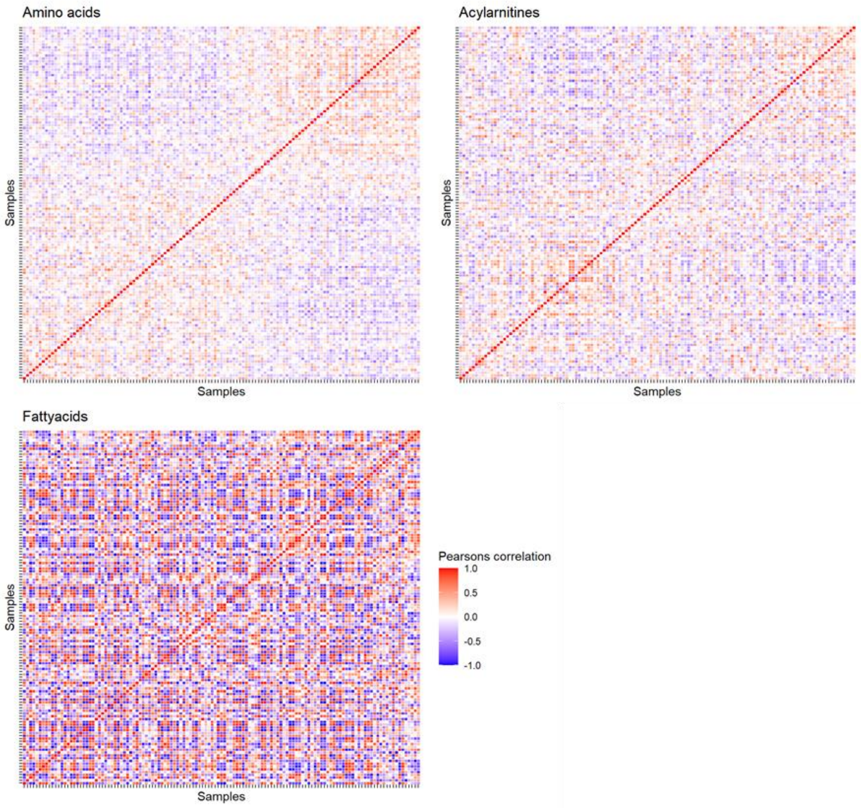

**Figure S46: Sample correlations of matrices of the different targeted metabolomics assays.**

The multivariate constrained ordination method Redundancy Analysis (RDA) was performed to determine if variation in the metabolite levels (response variables) can be explained by the measurement date (batch, explanatory variable). Additionally, RDA was performed with sex as a conditional variable to determine if any effect of batch is still present when taking sex into account. For amino acids, it was additionally tested if differences in metabolite levels can be explained by the number of times samples were frozen and thawed. This was done with and without batch as conditional variable. Analyses were conducted using the R package *vegan* (Oksanen, et al. 2022) .

Based on the RDA results, it was decided to perform batch effect adjustment but include sex as a covariate to account for the unbalanced batch design. To choose between the parametric and non-parametric ComBat methods, prior probability distribution plots were assessed. The parametric method was deemed suitable as empirical kernel density estimates of batch effects and parametric estimates of batch effects overlap.

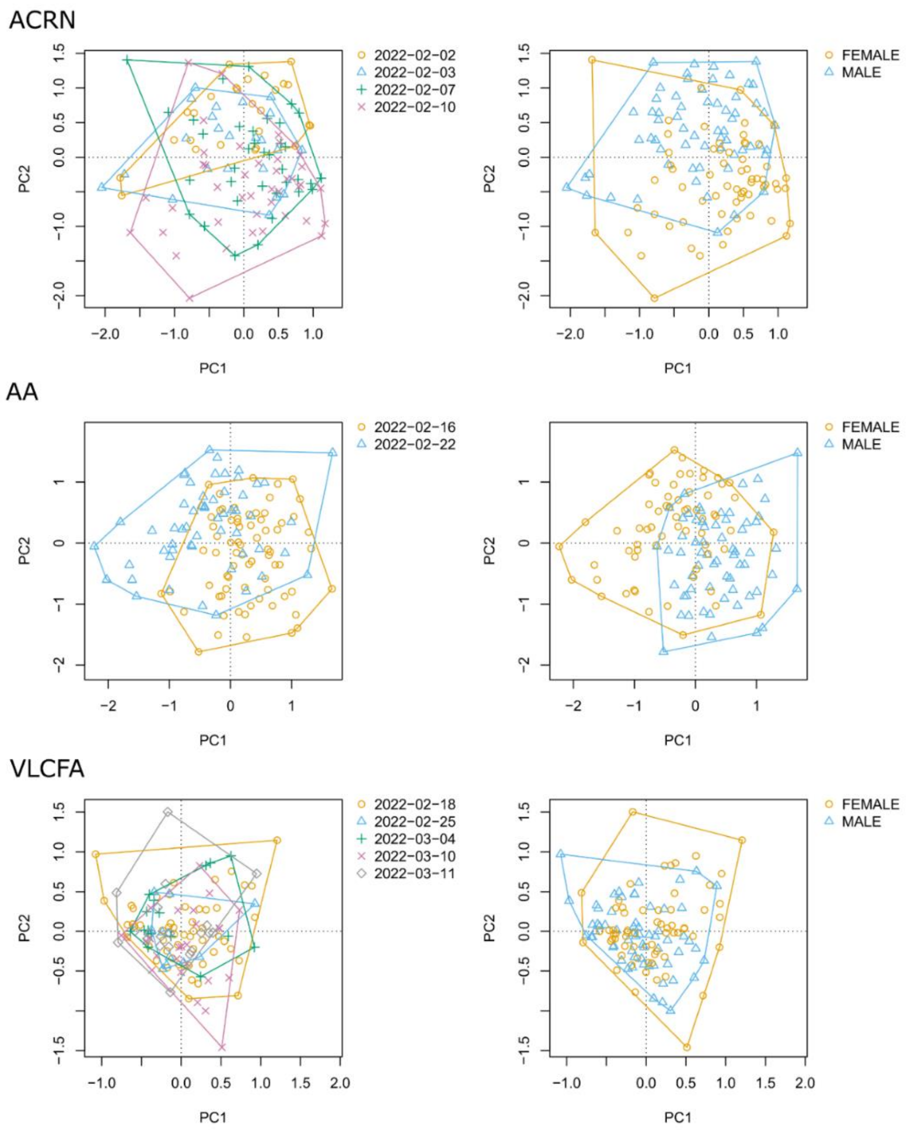

**Figure S47: Principal Component Analysis score plots of targeted metabolomics assays (acylcarnitines, ACRN; amino acids, AA; very long chain fatty acids, VLCFA) before batch effect adjustment.** Features with missing values were excluded from the analysis. Metabolite levels were log-transformed and standardized. Samples are colored by measurement date (left) and sex (right).

### Missing value imputation

#### Metabolomics

Missing values were found in the metabolomics data across the different assays and batches (Fig S29; Table S3). While some analysis methods can handle missing values other methods require complete observations. Therefore, missing values were imputed. Since the metabolomics assays are quantitative targeted measurements, the true values are assumed to be between 0 µM and the lower limit of detection. This means that values are missing not at random (MNAR). Several imputation methods were applied including both methods recommended for values missing at random (MAR) and MNAR. The distributions of imputed values were visually compared to choose a suitable method. Criteria were that imputed values should be in the lower range of the observed values but not distorting the overall value distribution.

*Random Forest*

Missing value imputation with Random Forest was performed using the R package missForest (Stekhoven and Buhlmann 2012, Stekhoven 2022) . Previously, this method was reported to be suitable for data with values MAR or missing completely at random (MCAR) (Wei, et al. 2018) . The number of trees per forest was set to $ntree=1000$ and the maximum number of iterations was set to $maxiter=10$. Out-of-bag (OOB) mean squared errors (MSE) were returned per variable (metabolite). Normalised root MSR (NRMSE) were calculated per variable as $NRMSE=MSEmean(observed)$.

*k-nearest neighbours (kNN)*

The R/Bioconductor package impute (Hastie, et al. 2022) was used to impute missing values using k-nearest neighbours averaging.

*Quantile regression imputation of left-censored data (QRILC)*

QRILC is implemented in the R package imputeLCMD (Lazar and Burger 2022) . This method was previously described to be applicable for values that are MNAR (Wei, et al. 2018) . To avoid distortion of the value distribution values are randomly drawn from a truncated normal distribution (Wei, et al. 2018) .

*No-skip kNN*

No-skip kNN is a modified version of kNN developed for MNAR (Lee and Styczynski 2018) . The internal function nsKNN from the R package MAI (Dekermanjian, Shaddox, et al. 2022a) was used to impute missing values.

*Single imputation*

This method is an alternative method for imputing values MNAR (Dekermanjian, Shaddox, et al. 2022) . The internal function kapImpute2 from the R package MAI (Dekermanjian, Shaddox, et al. 2022a) was employed.

*Low abundance resampling (LAR)*

Missing values were imputed by randomly sampling from the 5% observed values with lowest abundancy.

*Mechanism-aware imputation (MAI)*

This two-step approach implemented in the R package MAI (Dekermanjian, Shaddox, et al. 2022a) first estimates the pattern of missingness (MNAR, MCAR) for each variable (Dekermanjian, Shaddox, et al. 2022) . Based on the estimated missingness pattern, Random Forest is used for MCAR and no-skip-kNN is used for MNAR.

Summary

Bee swarm plots (Figure S47, Figure S48) show that both Random Forest (OOB NRMSE ranged between 0.07 for citrulline and 0.91 for argininosuccinic acid) and MAI produced imputed values that range around the mean of the measured values. kNN produced imputed values higher than any measured values for some metabolites (5-aminolevulinic acid, saccharopine). QRILC clearly distorts the value distribution by producing extremely low values. No-skip kNN produced a mixture of values ranging around the mean of the measured values and low values. Both single imputation and LAR produced low values, but the values imputed by LAR have a higher spread especially for beta-alanine and ethanolamine.

| Assay | Number of analysed compounds | Number of compounds not detected in any sample | Number of compounds with ≥30% (and <100%) missing values | Number of compounds after filtering |
|---|---|---|---|---|
| ACRN | 43 | 6 | 4 | 33 |
| AA | 53 | 0 | 7 | 46 |
| VLCFA | 5 | 0 | 0 | 5 |

**Table S4: Overview of numbers of measured metabolites before and after filtering based on proportion of missing values.**

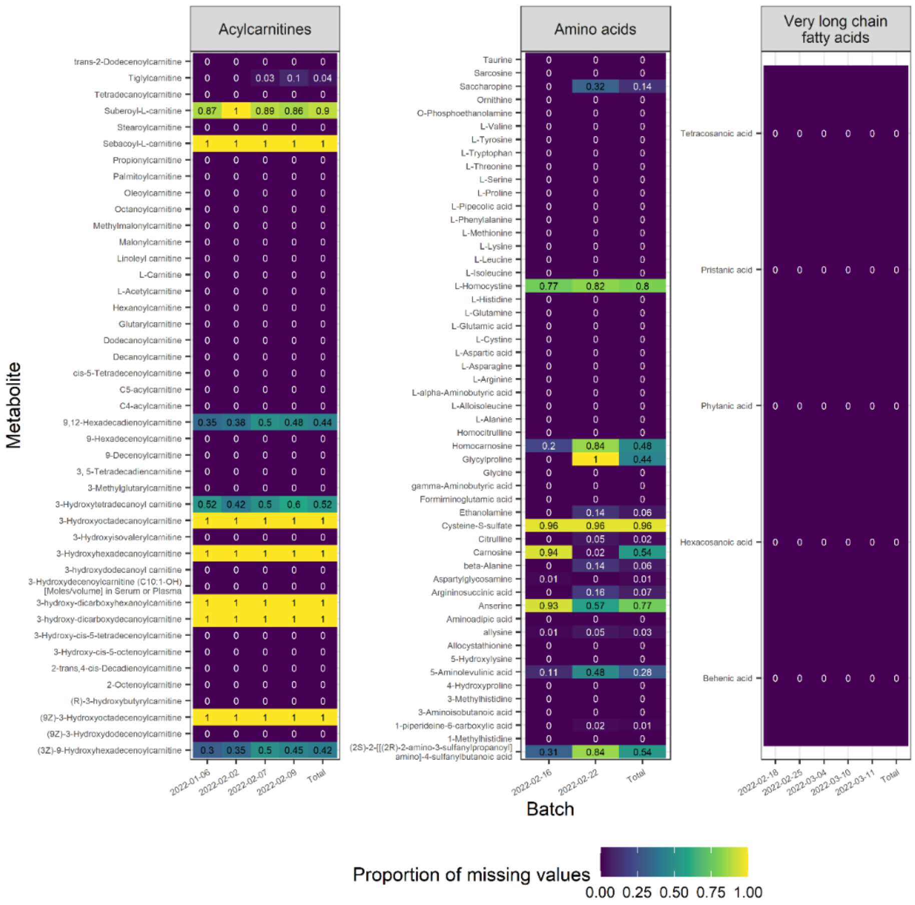

**Figure S48: Proportion of missing values per metabolite and measured batch as well as across all batches (total) for targeted metabolomics assays acylcarnitines, amino acids and very long chain fatty acids.**

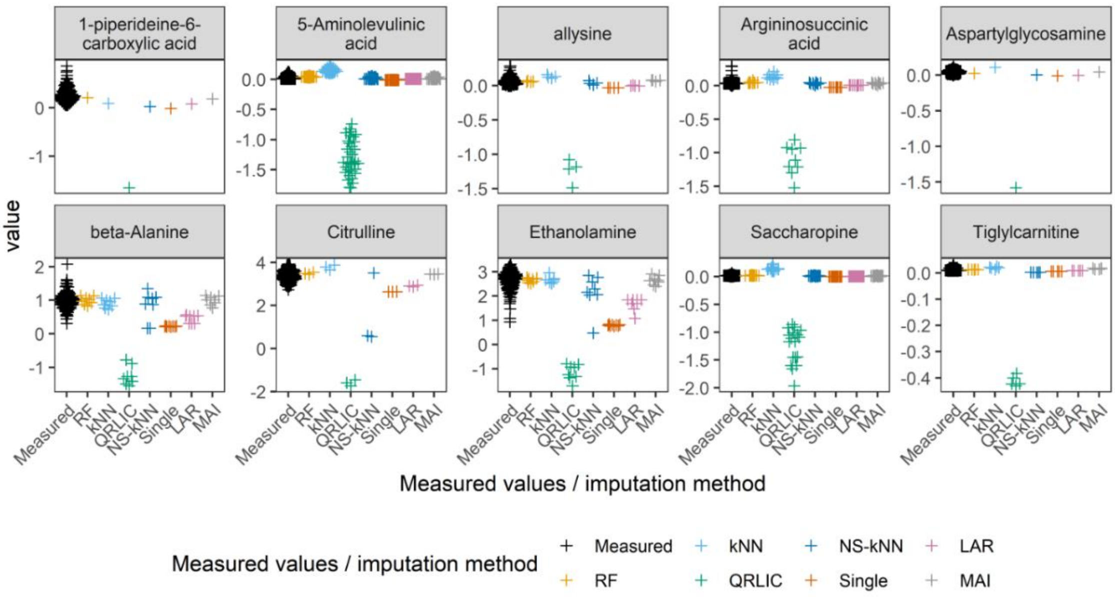

**Figure S49: Distribution of measured and imputed metabolite levels.** Prior to missing value imputation, metabolites were filtered, levels were log-transformed and batch effect-adjusted using the parametric ComBat method with sex as covariate. Imputation methods: RF, Random Forest; kNN, k-nearest neighbour; QRLIC, quantile regression imputation for left-censored data; NS-kNN, no-skip-kNN, Single, single imputation, LAR, low abundance resampling (5%), MAI, mechanism-aware imputation.

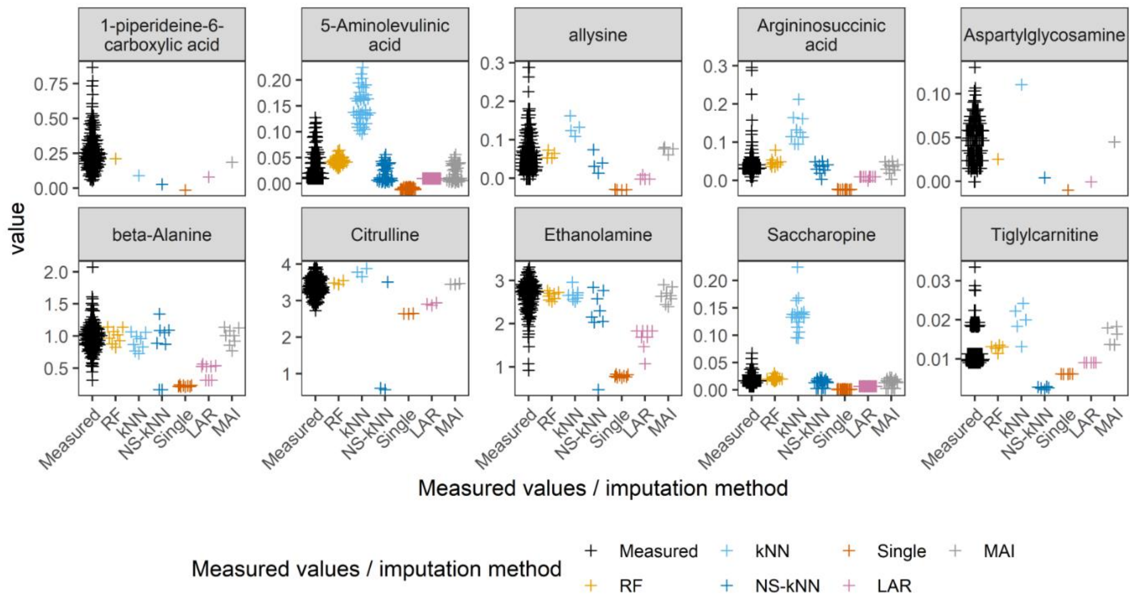

**Figure S50. Distribution of measured and imputed metabolite levels.** Prior to missing value imputation, metabolites were filtered, levels were log-transformed and batch effect-adjusted using the parametric ComBat method with sex as covariate. Imputation methods: RF, Random Forest; kNN, k-nearest neighbour; NS-kNN, no-skip-kNN, Single, single imputation, LAR, low abundance resampling (5%), MAI, mechanism-aware imputation.

| Assay | Batch | Number (percentage) of samples from female individuals | Number (percentage) of samples from male individuals |
|---|---|---|---|
| ACRN | 2022-01-06 | 3 (13%) | 20 (87%) |
| | 2022-02-02 | 4 (15%) | 22 (85%) |
| | 2022-02-07 | 23 (64%) | 13 (36%) |
| | 2022-02-09 | 34 (81%) | 8 (19%) |
| AA | 2022-02-16 | 19 (27%) | 52 (73%) |
| | 2022-02-22 | 45 (80%) | 11 (20%) |
| VLCFA | 2022-02-18 | 10 (19%) | 44 (81%) |
| | 2022-02-25 | 5 (45%) | 6 (55%) |
| | 2022-03-04 | 11 (69%) | 5 (31%) |
| | 2022-03-10 | 18 (69%) | 8 (31%) |
| | 2022-03-11 | 20 (100%) | 0 (0%) |

**Table S5: Number of samples from males and females per batch and assay.**

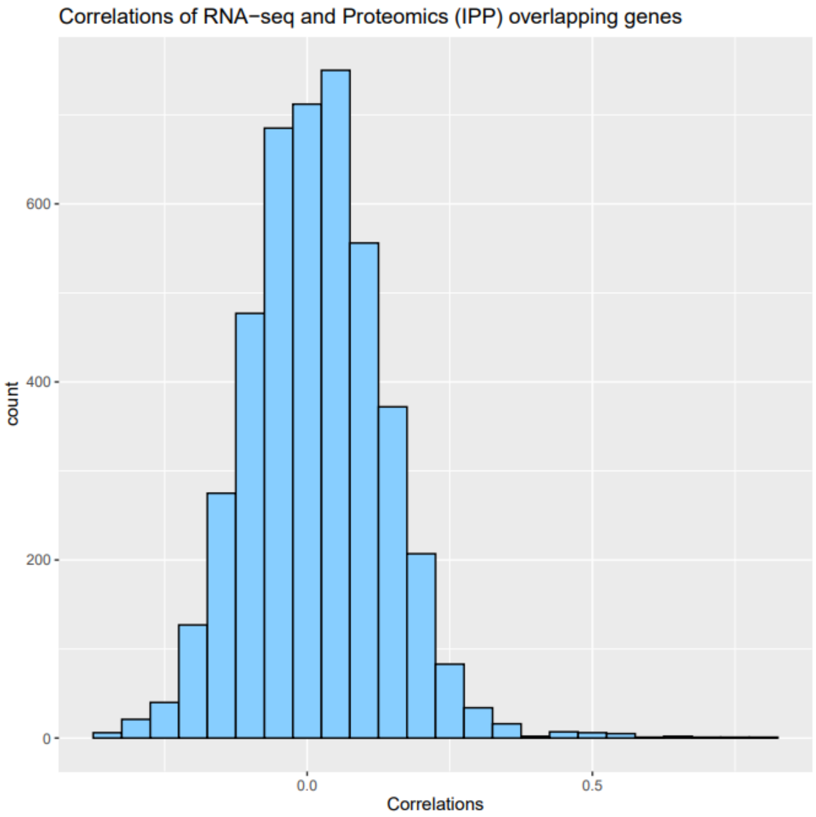

**Figure S51: Correlations of protein and RNA levels of the same genes.**

#### Lipidomics compounds discoverer

Table S4 summarises the number of detected compounds in the dataset according to their identification confidence level. MSI ID Level 2 was done by comparing MSMS data to spectral database, followed by the assigning LipidMAPS ID (LMID) – this dataset will be used for following multi-omics analyses.

We have performed principal component analysis (PCA) to evaluate the human plasma samples (n=127) data in positive ionisation mode (Figure S33) and negative ionisation mode (Figure S34). Only putatively annotated compounds (positive ionisation mode, n=196; negative ionisation mode, n=175) were used to create PCA models. Main differences were observed based on gender segregation. The tight clustering of quality control samples in the middle of the PCA confirms a very low analytical variance in the data, leaving the rest of the clustering to biological influence.

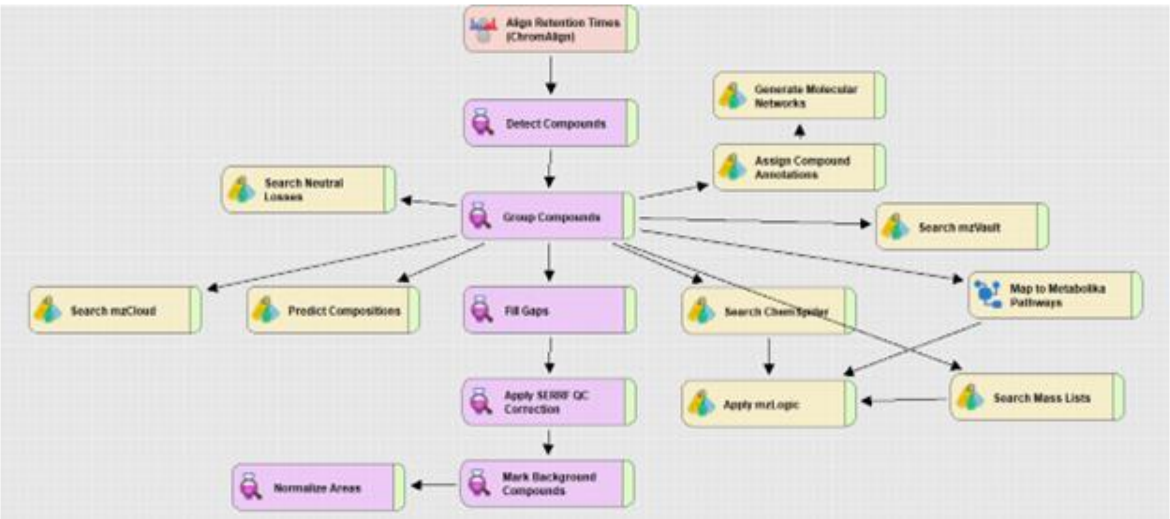

**Figure S52: Schematic workflow applied in Compound Discoverer 3.3 SP1**
